# Local translation drives glioblastoma heterogeneity and tumor invasion

**DOI:** 10.64898/2026.05.02.722387

**Authors:** Nikolas Layer, Niklas Wißmann, Marc Philipp Dehler, Maximilian F. Lasotta, Madlena Yan Ly, Mahammad Bashirov, Tim Wageringel, Jiajie Liu, Robin S. Dietl, Ekin Reyhan, Marc C. Schubert, Svenja Kristin Tetzlaff, Nirosan Sivapalan, Yiheng Tang, Janus Mosbacher, Yeunjung Ko, Tongil Ko, Rangel Lyubomirov Pramatarov, Stella J. Soyka, Yvonne Yang, Yahaya Yabo, Atefeh Pourkhalili Langeroudi, Obada T. Alhalabi, Robert Denninger, Michael Botz, Vivek Kumar Sahu, Nathalie Wilke, Isabell Bludau, Benjamin Deneen, Ana C. deCarvalho, Thomas V. Wuttke, Stefan Pfister, David T. W. Jones, Bogdana Suchorska, Ashok Kumar Jayavelu, Dirk Trauner, Felix Sahm, Dieter Henrik Heiland, Erin M. Schuman, Varun Venkataramani

## Abstract

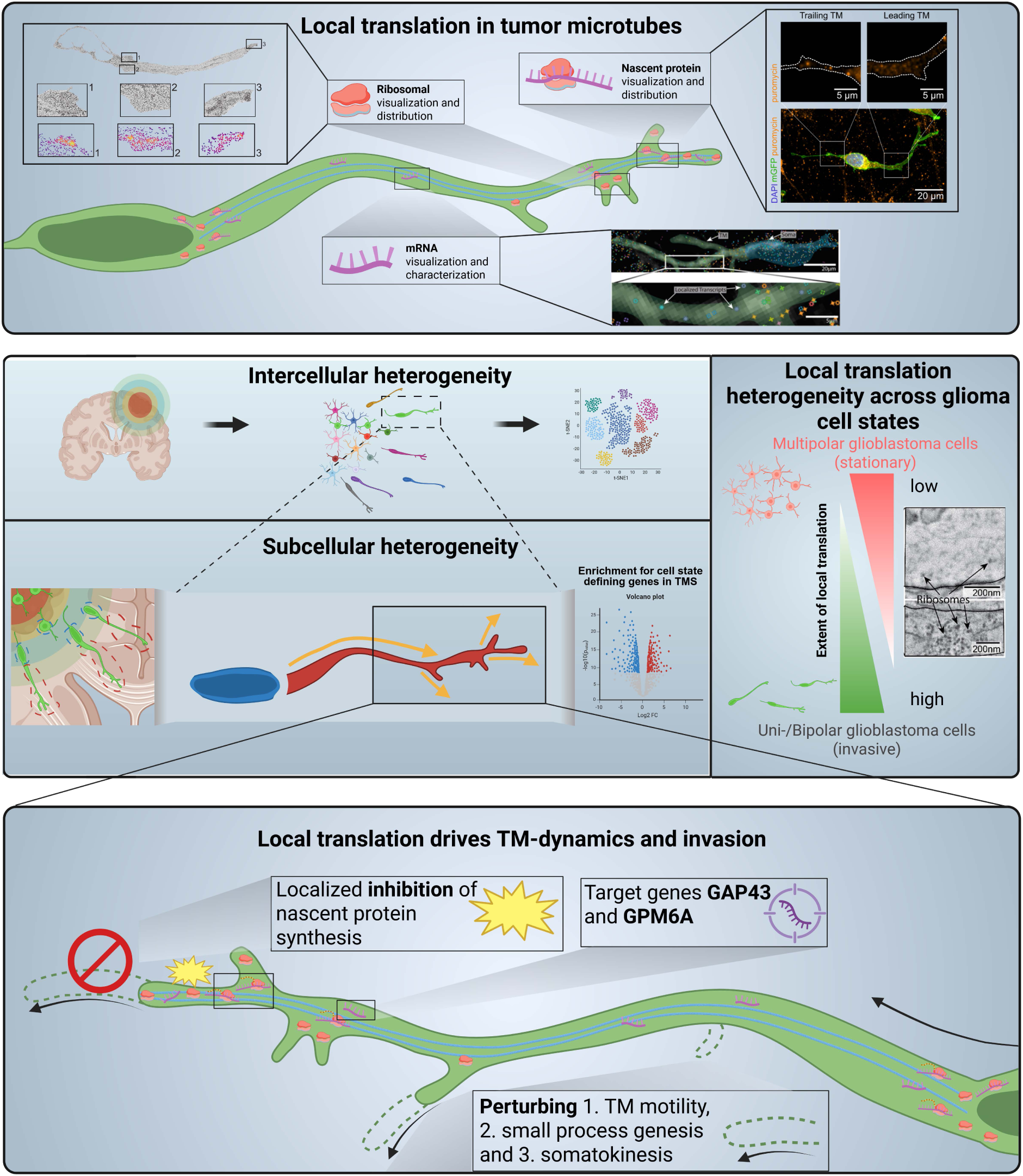

Glioblastoma is characterized by diffuse brain invasion, yet the subcellular mechanisms enabling this aggressive behavior remain poorly understood. A subpopulation of glioblastoma cells forms invasive tumor microtubes (TMs), neurite-like extensions that drive whole-brain colonization. Here, we establish local protein translation as a fundamental driver of TM dynamics and invasive cell states. Developing a subcellular transcriptomics approach - integrating subcellular organelle organization with spatially resolved transcriptomics and functional readouts - we reveal that TM gene expression drives cell state identity. Invasive cells further exhibit significantly elevated local translation in protruding TMs, directly linking subcellular protein synthesis to functional invasive states associated with neurodevelopmental programs of axonal growth cones. Targeted disruption of TM-localized translation via photoswitchable puromycin, and specific knockdowns of the TM-enriched proteins GPM6A and GAP43, impaired TM dynamics, suppressed invasion, and reduced tumor growth. Together, these findings define local translation as a key determinant of tumor heterogeneity and glioblastoma invasion.

## Main Text

Glioblastoma (GB) is the most aggressive and lethal primary brain tumor in adults, characterized by pronounced heterogeneity at both the cellular and molecular levels (*1–4*). This cellular and molecular plasticity contributes to infiltrative behavior and resistance to therapy (*4–15*). One hallmark of GB pathophysiology is its highly invasive nature mediated through dynamic remodeling of neurite-like membrane protrusions called tumor microtubes (TMs) (*8*, *16–19*). These highly motile structures not only facilitate invasion into the surrounding brain parenchyma but also support intercellular communication and resistance to therapy (*16*). Importantly, TM motility is a key driver of GB cell invasion (*20–23*). Local translation, the spatially and temporally restricted synthesis of specific proteins from mRNAs by ribosomes, has emerged as a key regulator of rapid, localized cellular responses in polarized or morphologically complex cells in the central nervous system (*24–26*). In the context of neuronal development and function, local translation regulates precursor cell development (*26*, *27*), axon guidance (*28–30*), enables cells to respond to microenvironmental cues (*31*, *32*) and contributes to synaptic plasticity (*33*, *34*). However, whether glioblastoma cells, which share morphological features and gene expression programs with neuronal and glial precursor lineages (*2*, *10*, *35*, *36*), exploit this neural mechanism of local translation to promote tumor invasion and cellular remodeling remains unknown (*37*). Another fundamental but unresolved question is whether glioblastoma cell state identity - the molecular programs that define distinct tumor cell behaviors - is determined at the level of the whole cell or within specific subcellular compartments such as TMs.

In this study, we dissected the structural and activity-dependent subcellular heterogeneity within GB cells and demonstrated that local translation within TMs promotes invasion, whole-brain colonization and can be therapeutically leveraged.

### All hallmarks of local translation are present in neurite-like structures of GB cells

Glioblastoma cells exhibit remarkable plasticity through constant formation and retraction of highly dynamic TMs (*7*). Comparing the dynamic behaviour of TMs and cell somata *in vivo*, we found that TMs displayed a significantly increased surface area change and motility during brain invasion compared to somatic compartments (Fig. 1C-E, Movie S1). We therefore hypothesized that these subcellular dynamics would require local protein synthesis at sites of active cellular remodeling - a mechanism well-established in neuronal development (Fig. 1A-B) (*38*). Subsequently, we used correlative light and electron microscopy (CLEM) in combination with a bespoke semi-automated ribosome detection pipeline (fig. S1A-B, see Materials and methods) and detected ribosomes in TMs and somata of GB xenograft mouse models (PDX) (Fig. 1F, Movie S2) and across patient tissue (Fig. 1H, fig. S1C). To demonstrate the presence of mRNA as a second hallmark of local translation within GB TMs, we investigated GB patient tumors and patient-derived model systems (Fig. 1G) with spatial transcriptomics, consistently revealing RNA transcripts within both GB cell somata and TMs. To assess the site-specific translation at ribosomes localized within TMs, we used a puromycilation assay to visualize translational activity (*39*), which enables an antibody-based detection of nascent polypeptide chains through incorporation of puromycin (fig. S1D, see Materials and methods)as an orthogonal readout of local nascent protein synthesis (*40*, *41*). These assay demonstrated nascent protein synthesis along TMs and smaller neurite-like processes of GB cells across model systems (Fig. 1I, fig. S1E-F), indicating active local translation (fig. S1G) (*34*, *42*). We observed all major hallmarks of local translation across 14 different patient-derived GB model systems, PDX models and multiple resected primary tissue from GB patients, (table S2, fig. S2, S3). Notably, local translation was consistently observed across five patient-derived pediatric glioma models, extending the relevance of this mechanism beyond adult glioblastoma (fig. S3). Based on the quantitative ribosomal distribution, we further found heterogeneous hotspots regions in the TM associated with dynamic TM remodelling. As previously described, GB cell architecture comprises both larger diameter primary TMs and small processes with particularly high turnover rates (*10*). Intravital two-photon imaging of S24 GB cells (Fig. 1J, Movie S3) revealed a significantly increased rate of protrusion formation and retraction in small processes compared to primary TMs (Fig. 1 K-L). Correlating with this dynamic behaviour, both scanning electron (Fig. 1M-P) and light microscopy (fig. S5) demonstrated elevated ribosomal density at branch points, sprouting segments, and distal TM tips, implicating increased translational demand at these dynamic subregions and indicating functional microdomains of enhanced translational capacity. To probe whether local protein synthesis in GB cells had an impact on cell behavior and TM motility, we next selectively perturbed translation within individual TMs, employing photoactivation of puroswitch, a photoswitchable puromycin analog (*43*) in organotypic slices of resected human tissues (*44*, *45*) injected with patient-derived GB cells and neuron-tumor cultures (Fig. 1Q-R, fig. S4). Light-based activation of puroswitch in single primary TMs synchronized to inactivation in the remaining GB cell induced arrest of TM motility or retraction, resulting in an overall reduction of morphologic TM plasticity (Fig. 1S-T, fig. S4C). This demonstrates that local translation is a mechanistic driver of glioblastoma invasion and opens new therapeutic avenues. The spatially restricted inhibition of translation using photoactivatable puromycin impaired GB cell TM dynamics, validating TM-localized translation as a targetable vulnerability that may extend beyond glioblastoma to other invasive cancers with morphologically complex cellular protrusions. Together, all hallmarks of local translation in neurite-like protrusions were identified across patient tissue and patient-derived model systems (*46*).

**Fig. 1.**
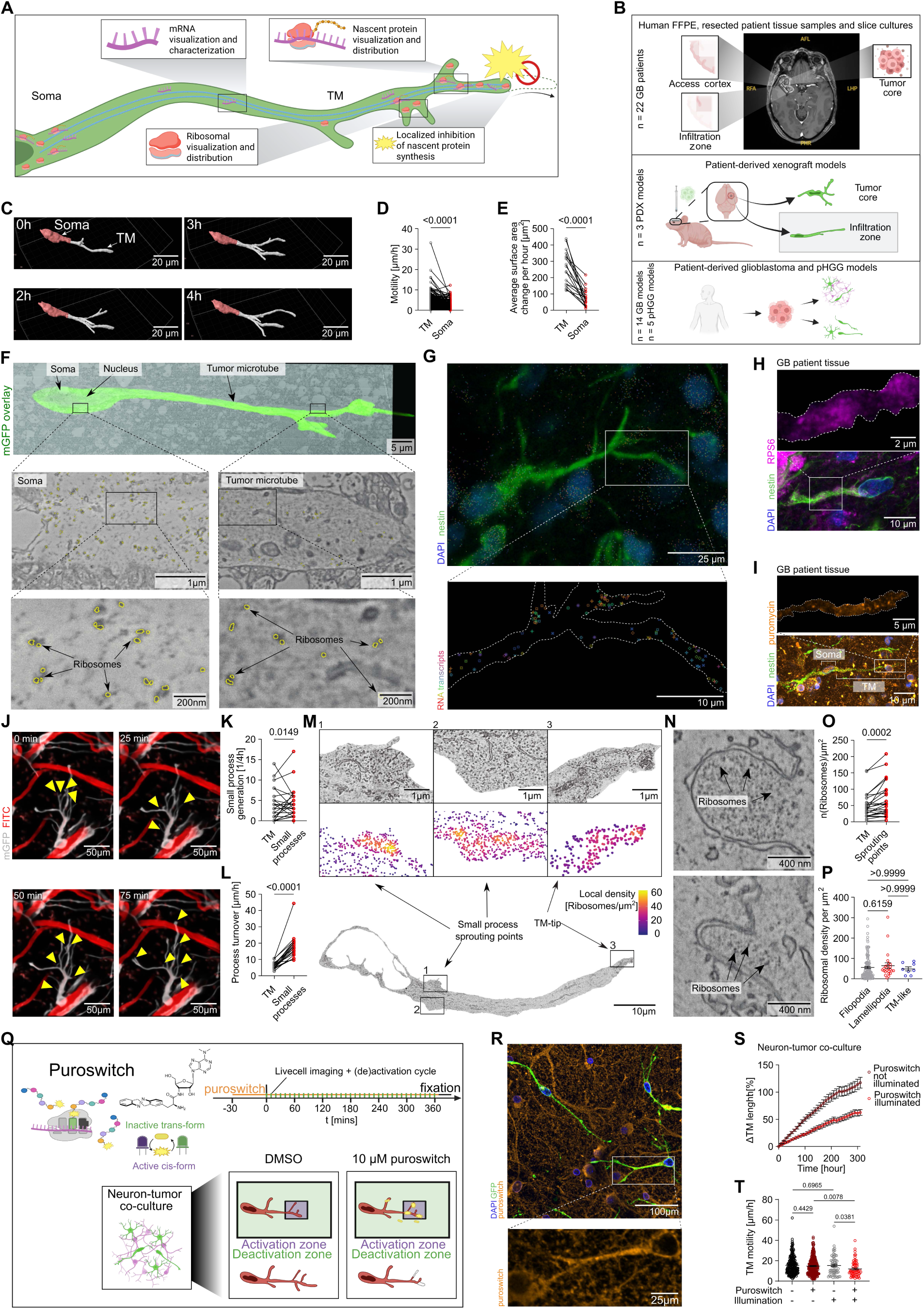
Local translation is a hallmark feature of tumor microtubes in glioblastoma cells. (**A**) Schematic illustration showing hallmarks and specific targeting of local translation in TMs. (**B**) Model systems of high-grade glioma to investigate the role of local translation used in this study (n = 22 GB patients, n = 3 PDX models, n = 14 patient-derived glioblastoma models and n = 5 patient-derived pediatric diffuse high-grade glioma (pHGG) models). (**C**) *In vivo* two-photon microscopy 3D rendering of a GB cell in PDX tissue during brain invasion, showing segmented somatic (red) and tumor microtube (TM, white) compartments. (**D**) Quantification of average displacement of TMs and paired somata over the time frame of 4-hour in vivo two-photon microscopy recordings (n = 351 GB cells), determined by manual tracking using MTrackJ (ImageJ). (**E**) Quantification of cell membrane surface change in TMs and somata of GB cells in PDX tissue during invasion using 4 hour in vivo two-photon microscopy (n = 18 GB cells). (**F**) Electron microscopic image overlaid with the corresponding confocal image of a S24 GB cell (green) in PDX tissue, revealing ribosomes in both the somatic and TM compartments. (**G**) Representative visualization of spatially correlated RNA in GB patient FFPE sections via the Xenium platform, stained for DAPI (blue) and nestin (green). Zoom-in displays different transcripts (multicolor) in the TM with a crop-out of the GB cell outline (dashed line).(**H**) Representative confocal IHC image (standard deviation based projection) of a GB cell in a GB patient brain section stained for DAPI (blue), nestin (green) and RPS6 (magenta). Zoom-in displays RPS6 signal in the TM with a nestin signal based mask of the cell outline (dashed line). (**I**) Representative confocal IHC image of a S24 GB cell in an acute GB patient brain section, stained for DAPI (blue), nestin (green) and puromycin (orange). Zoom-in displays puromycin signal in the TM with a nestin signal based mask of the cell outline (dashed line). (**J**) Representative TM motility of a complex S24 GB cell in PDX tissue (white) over a time frame of 4-hours during intravital two-photon imaging combined with an angiogram (red) (post-processed with enhance.ai in NIS Elements for representation, maximum intensity based projection). Yellow arrow heads indicate transient sprouting of new small processes. (**K**) Quantification of small process genesis events compared to formation of new TMs of GB cells in PDX tissue during 4 hour in vivo two-photon imaging (n = 21 cells). (**L**) Quantification of small process turnover speed compared to TM motility of GB cells in PDX tissue during 4 hour in vivo two-photon imaging (n = 21 cells). (**M**) Electron microscopic representation of ribosomal hotspots at TM tips and sprouting points of GB cells in PDX tissue. Color coding displays the local ribosomal density in selected cellular subcompartments according to the legend. (**N**) Representative EM image of ribosomes in morphologically different small processes of a S24 GB cell in PDX tissue. (**O**) Quantification of average ribosomal density in EM images of sprouting points compared to the overall ribosomal density in the corresponding TM of GB cells in PDX tissue (n = 183 small process sprouting points in n = 25 cells). (**P**) Quantification of average ribosomal density in EM images of different morphologic small process classes in S24 GB cells in PDX tissue (n = 119 filopodia-like, n = 24 lamellipodia-like and n = 8 TM-like small processes).(**Q**) Workflow for spatial inhibition of local translation by optical activation of puroswitch within singular TMs, combined with simultaneous inactivation of paired cell somata in both human organotypic brain slice cultures and neuron-tumor co-cultures. (**R**) Anti-puroswitch staining using the anti-puromycin antibody (orange) of S24 GB cells in neuron-tumor co-culture treated with puroswitch and illuminated for 6 hours. Zoom-in displays puromycin signal in the TM (**S**) Quantification of the relative mean cumulative change of TM length over time in puroswitch-treated, illuminated compared to puroswitch-treated, non-illuminated S24 GB cells in neuron-tumor co-cultures (n = 278 puroswitch non-illuminated TMs, n = 65 puroswitch treated illuminated TMs). (**T**) Quantification of TM motility in illuminated and non-illuminated puroswitch treated S24 GB cells in neuron-tumor co-cultures (n = 209 DMSO non-illuminated TMs, n = 278 puroswitch treated non-illuminated TMs, n = 60 DMSO illuminated TMs, n = 65 puroswitch treated, illuminated TMs).

### Subcellular TM gene expression determines GB cell state

GB is characterized by extensive transcriptomic and morphological heterogeneity (*2*, *4*, *36*, *49*). We therefore investigated whether this heterogeneity manifests at the subcellular level, with TMs and somata maintaining distinct RNA landscapes that could underpin compartment-specific local translation. (Fig. 2A). For this purpose, we isolated soma- and TM-enriched RNA for bulk sequencing, using an insert-based assay that physically restricts somatic cell passage while permitting TM outgrowth into a second accessible compartment (Fig. 2A, see Supplementary Materials and methods) (*50*). Additionally, we developed an ultrastructural transcriptomics approach using MERFISH (*51*, *52*) to segment and analyze TM and somatic regions separately in GB cells with both invasive and non-invasive cell morphology (Fig. 2B, table S3). Differential gene expression and principal component analysis between somatic and TM compartments in both bulk and spatial transcriptomics revealed a distinct transcriptome in TMs compared to somata (Fig. 2F, fig. S7A-I). This subcellular compartmentalization was further supported by MERFISH analyses of glioblastoma patient tissue (*51*) comparing nuclear and cytosolic transcript pools (fig. S7J-M). Housekeeping and nuclear-enriched transcripts localized predominantly within the nuclear area, whereas cell state signature genes were enriched in non-nuclear compartments (fig. S7K). Benchmarking of TM-enriched gene signatures against cell state programs revealed that the predictive accuracy of TM signatures was lower in single-nucleus compared to single-cell data, indicating distinct subcellular RNA heterogeneity across the nucleus, cell somata, and TMs within GB cells (Fig. 2F, fig. S7L-M). GO-term enrichment analysis integrating both MERFISH and bulk RNA-seq revealed TM-enriched transcripts to be coherently associated with GO-terms of migration, growth cones and polarized growth, suggesting that the TM transcriptome is associated with functional programs of invasion (Fig. 2C-E). Further, an enrichment of ribosomal GO terms showed a potential connection to local biogenesis of ribosomes in TMs (Fig. 2D) (*53–55*). Deeper subclustering of TM transcript abundance from MERFISH segmentation uncovered further heterogeneity within TMs themselves (Fig. 2G), which correlated with cellular invasiveness as assessed by GB cell morphology (Fig. 2H). A gene module built from the top PC1-loading genes - the principal component most strongly driving inter-TM cluster differences - showed distinct expression levels across Neftel-classification-based cell states (*2*) in PDX single-cell RNA sequencing data (Fig. 2I, J).

**Fig. 2.**
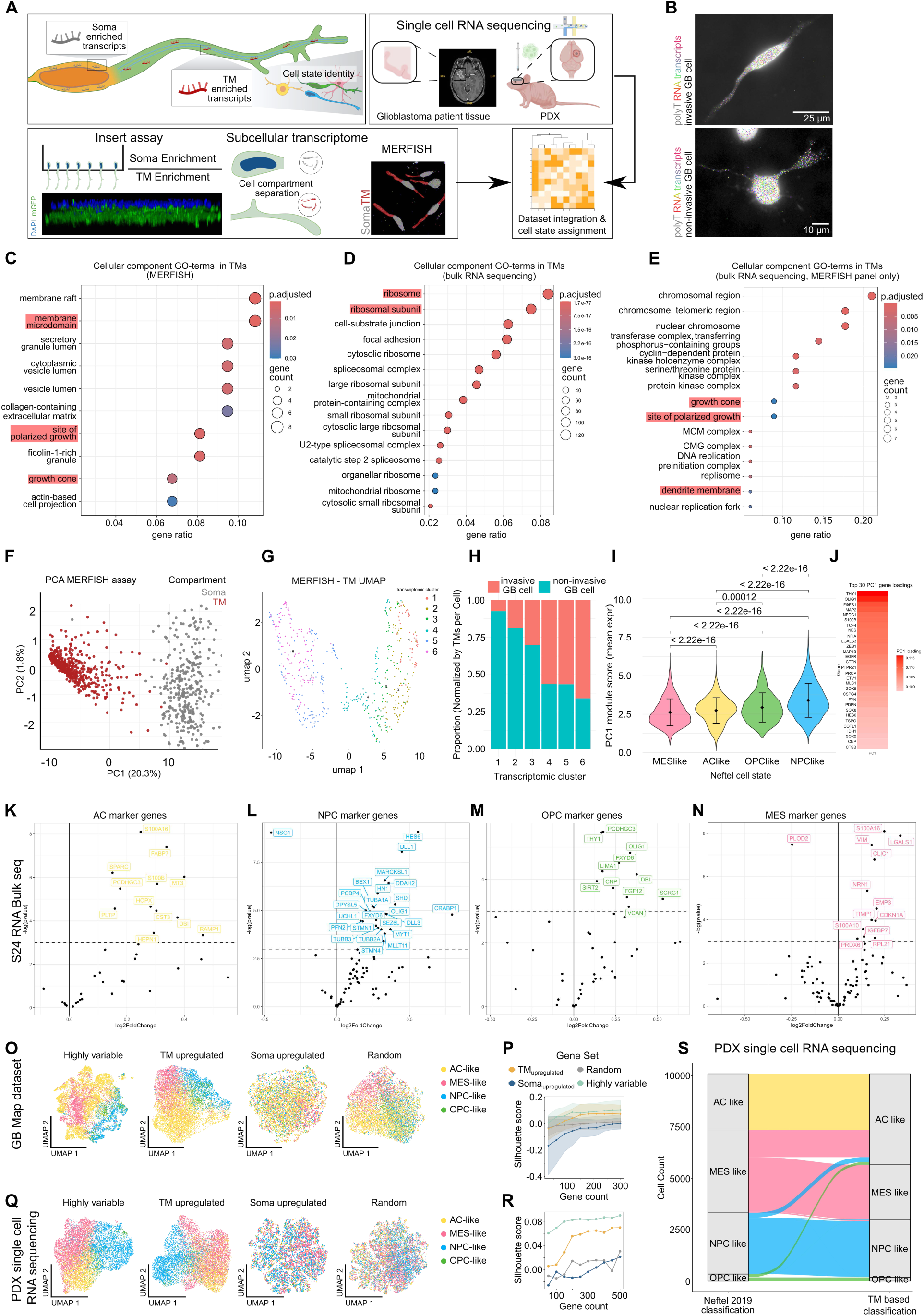
Subcellular transcriptomics of TMs reflects molecular heterogeneity of GB. (**A**) Schematic workflow of soma- and TM-enriched transcriptomic data integration with single-cell transcriptomic datasets from GB patient models and PDX models. (**B**) Representative MERFISH image of a GB cell with invasive compared to non-invasive morphology in mono-culture showing polyT signal (gray) and spatial distribution of RNA transcripts (multicolor). (**C**) Cellular component GO terms identified as associated with RNA transcripts enriched in TMs of S24 GB cell cultures in the MERFISH dataset.(**D**) Cellular component GO terms identified as associated with RNA transcripts enriched in the TM-enriched fraction of S24 GB cell cultures in the insert RNA-bulk sequencing dataset. (**E**) Cellular component GO terms identified for RNA transcripts enriched in the TM fraction of S24 GB cell cultures in the insert RNA bulk sequencing dataset, considering only transcripts being part of the MERFISH dataset. (**F**) PCA of transcripts located in TMs (red) and somata (grey) of S24 GB cell mono-cultures in the MERFISH dataset including 358 invasive GB cells manually segmented into TMs and soma. (**G**) UMAP displaying 6 clusters calculated on a PCA of transcripts located in the TM of invasive and non-invasive GB cells in mono-culture in the MERFISH dataset including 88 invasive and 98 non-invasive GB cells (dots represent single TMs). (**H**) Stacked bar plot showing proportions of TMs belonging to invasive (red) vs non-invasive (blue) cells over the TM-based cluster from (G), proportions normalized by the number of TMs of the respective cells. (**I**) Violin plot displaying a module score calculated from the top 30 genes contributing to the PC1 gene loadings of the TM-based PCA. Scores are shown for single cells from the S24 PDX scRNA dataset grouped by their cell state according to Neftel et al. (2) cell state classification (error bars = ±1 SD) (**J**) Heatmap showing the top 30 genes contributing to the PC1 loadings from the TM-based UMAP (G), along with their relative PC1 weights contributing to TM clustering.. (**K**) Volcano plot displaying differentially expressed genes (DEGs) from bulk sequencing in the Neftel et al. (*47*) meta-modules’ marker genes for AC-like cells with TM-vs-soma log₂-fold change on the x-axis and –log(p-value) on the y-axis. Genes with p-value <0.05 are labeled. (**L**) Volcano plot of meta-modules’ marker genes for NPC-like cells. (**M**) Volcano plot of meta-modules’ marker genes for OPC-like cells. (**N**) Volcano plot of meta-modules’ marker genes for MES-like cells. (**O**) UMAPs of 10,000 randomly sampled cells of the GB Map dataset (*48*), based on the principal component analysis (PCA) using 150 genes from one of four gene sets: the most highly variable (GB Map dataset), TM upregulated (bulk sequencing dataset), soma upregulated (bulk sequencing dataset), or random genes. For plotting, only those cells expressing > 5 genes were retained. (**P**) Plot comparing silhouette scores between the different gene sets from (O). Silhouette score (see materials and methods) quantifies the distinctness of GBM cell state clustering in the GB Map dataset and is displayed for each of the four gene sets across varying gene counts. Dots denote the average silhouette score across all samples calculated on the top 10 principal components after a sample-wise PCA. Shades represent the standard deviation across samples. (**Q**) UMAP representations of S24 GB cell clustering of from single-cell RNA PDX sequencing based on one of four gene sets: the most highly variable (GB Map dataset), TM upregulated (bulk sequencing dataset), soma upregulated (bulk sequencing dataset), or random genes. (**R**) Plot comparing the mean silhouette score between the different gene sets from (Q) analogue to O. (**S**) Riverplot visualizing S24 cluster assignment in the PDX scRNA dataset before (Neftel et al. (*2*) cell state classification) and after gene set reduction according to S24 GB cell TM-upregulated genes.

To independently validate the observation that the TM-enriched transcriptome reflects glioblastoma cell state identity, we examined the distribution of established cell state signature genes within TM-enriched bulk RNA-seq data (*2*). TM-enriched fractions showed significant upregulation of genes associated with astrocyte-like (AC), oligodendrocyte progenitor-like (OPC), neuronal progenitor-like (NPC), and mesenchymal-like (MES) cell states compared to soma-enriched fractions (Fig. 2K-N). UMAP clustering revealed that globally highly variable gene sets produced the strongest cell state separation - as quantified by silhouette scores - consistent with their variability across cell states. Among targeted gene sets, TM-upregulated genes achieved markedly improved cluster separation compared to soma-upregulated and randomly selected gene sets from the GBMap dataset (Fig. 2O, P). Furthermore, re-clustering of PDX single-cell RNA sequencing data using TM-enriched transcripts fully preserved established cluster identities (Fig. 2Q-S).

Collectively, these findings demonstrate that cell state signature transcripts are predominantly compartmentalized to TMs rather than somata, and that TM-enriched mRNA signatures faithfully reconstruct established glioblastoma cell state classifications. This establishes the transcriptomic landscape of TMs as a primary determinant of glioblastoma cell state identity.

Having established that TM transcriptomes are linked to cell state and invasive behavior, we next asked whether the observed transcriptomic heterogeneity between invasive and stationary GB cells is also reflected in the spatial distribution of ribosomes and translational activity (Fig. 3A). To comprehensively assess the translational machinery, we characterized ribosomal subcellular localization associated with organelles in TMs (*56*). A subfraction of ribosomes was bound to (55) the endoplasmic reticulum (ER) (fig. S6A-B), which is associated with translation of membrane and secreted proteins (*57–59*). Additionally, TM ribosomes were preferentially organized as monosomes rather than polysomal clusters (fig. S6C-E) potentially reflecting a preference for the translation of a subset of differentially expressed transcripts (*56*). Additionally, we compared ribosomal abundance at the ultrastructural level between two distinct glioblastoma cell populations: invasive, morphologically simpler cells bearing fewer than three primary TMs (*10*, *60*), and more stationary, highly branched cells with complex morphology as confirmed with timelapse *in vivo* imaging of PDX models (Fig. 3B-D). Importantly, MERFISH-based assessment of transcript density (Fig. 3E) revealed increased RNA abundance in TMs of invasive GB cells and electron microscopy analysis demonstrated a significantly higher ribosomal density in TMs of invasive cells, both relative to TM volume (Fig. 3F) and somatic ribosome density (Fig. 3G). To determine whether these differences in ribosomal density and RNA translated into differential nascent protein production, we performed immunofluorescence labeling of puromycin and ribosomal marker RPS6 in PDX tissue sections, followed by super-resolution confocal microscopy (Fig. 3H-I). Across patient-derived glioblastoma models (fig. S8A-C), invasive tumor cell TMs exhibited both increased levels of local translation (Fig. 3J-K) and significantly elevated ribosomal density (Fig. 3L-M). Moreover, we found a strong correlation between ribosomal and puromycin signal intensities in TMs (Fig. 3N-O) across PDX models, suggesting that ribosomal abundance can serve as a surrogate marker for mRNA translation in glioblastoma. The correlation between increased ribosomal density in TMs and cell invasiveness across glioblastoma cell states (*2*, *61*) indicated that local translation is particularly critical for tumor invasion, establishing a direct link between biological function and nascent protein synthesis in glioblastoma.

**Figure 3.**
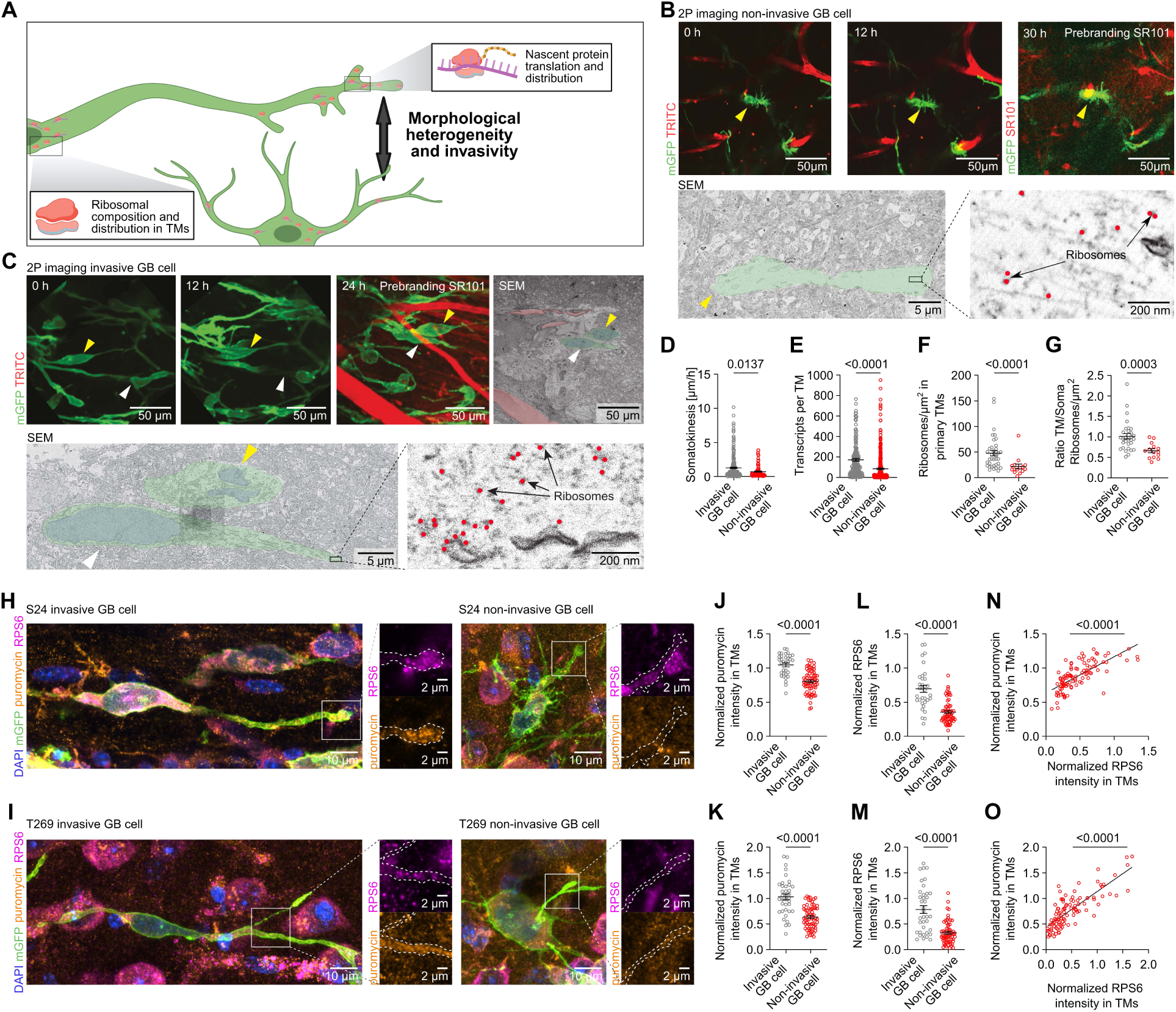
Invasive cell states are enriched for local translation. (**A**) Schematic illustration highlighting the differences in aspects of local translation between cells with distinct morphologies and invasive behaviors. (**B**) Representative TM motility of a stationary, non-invasive S24 GB cell (yellow arrowheads) in PDX tissue, during intravital two-photon imaging over a time frame of 24 hours, combined with a fluorescent angiogram (red) and the correlative SEM image showing the ribosomal density in the TM. (**C**) Representative TM motility of highly motile, invasive S24 GB cells (white and yellow arrowheads) in PDX tissue, during intravital two-photon imaging over a time frame of 24-hours combined with an angiogram (red) and the correlative SEM image showing the ribosomal density in the TM. (**D**) Quantification of somatic motility of S24 GB cells with invasive compared to non-invasive morphology in PDX tissue during intravital two-photon imaging over a time frame of 24 hours (n = 224 invasive TMs, n = 85 non-invasive TMs). (**E**) Quantification of number of transcripts per TM of S24 GB cells with invasive compared to non-invasive morphology (n = 161 invasive GB TMs of 88 GB cells, n = 350 non-invasive GB TMs of 98 GB cells). (**F**) Quantification of ribosomal density in primary tumor microtubes of GB cells with invasive compared to non-invasive morphology in ex vivo SEM images (n = 33 invasive TMs, n = 67 non-invasive TMs). (**G**) Ratio of ribosomal density in TMs to ribosomal density in the soma of GB cells with invasive compared to non-invasive morphology in ex vivo SEM images (n = 30 invasive cells, n = 14 non-invasive cells). (**H**) Representative confocal IHC image of S24 GB cells in PDX tissue with invasive and non-invasive morphology, stained for DAPI (blue), mGFP (green), puromycin (orange) and ribosomal protein S6 (RPS6, magenta). Zoom-ins display RPS6 or puromycin signal in the TM with a mGFP signal based mask of the cell outline (dashed line). (**I**) Representative confocal IHC image of T269 GB cells in PDX tissue with invasive and non-invasive morphology, stained for DAPI (blue), mGFP (green), puromycin (orange) and ribosomal protein S6 (RPS6, magenta). Zoom-ins display RPS6 or puromycin signal in the TM with a mGFP based mask of the cell outline (dashed line). (**J**) Quantification of soma-normalized puromycin signal intensity between TMs of S24 GB cells in PDX tissue with invasive compared to non-invasive morphology (n = 33 invasive TMs, n = 67 non-invasive TMs). (**K**) Quantification of soma-normalized puromycin signal intensity between TMs of T269 GB cells in PDX tissue with invasive compared to non-invasive morphology (n = 37 invasive TMs, n = 65 non-invasive TMs). (**L**) Quantification of soma-normalized RPS6 signal intensity between TMs of S24 GB cells in PDX tissue with invasive compared to non-invasive morphology (n = 33 invasive TMs, n = 67 non-invasive TMs). (**M**) Quantification of soma-normalized RPS6 signal intensity between TMs of T269 GB cells in PDX tissue with invasive compared to non-invasive morphology (n = 37 invasive TMs, n = 65 non-invasive TMs). (**N**) Correlation of soma-normalized RPS6 and puromycin intensity in the TMs of S24 GB cells in PDX tissue (n = 100 TMs). (**O**) Correlation of soma-normalized RPS6 and puromycin intensity in the TMs of T269 GB cells in PDX tissue (n = 102 TMs).

### Neuron-glioma synapses and calcium events correlate with local translation

Given the functional connectivity of GB cells through neuron-glioma synapses (NGS) (62–67) and multicellular networks (68) leading to calcium events within GB cells, we next investigated whether calcium events are correlated with translational activity within TMs (Fig. 4A).

**Figure 4.**
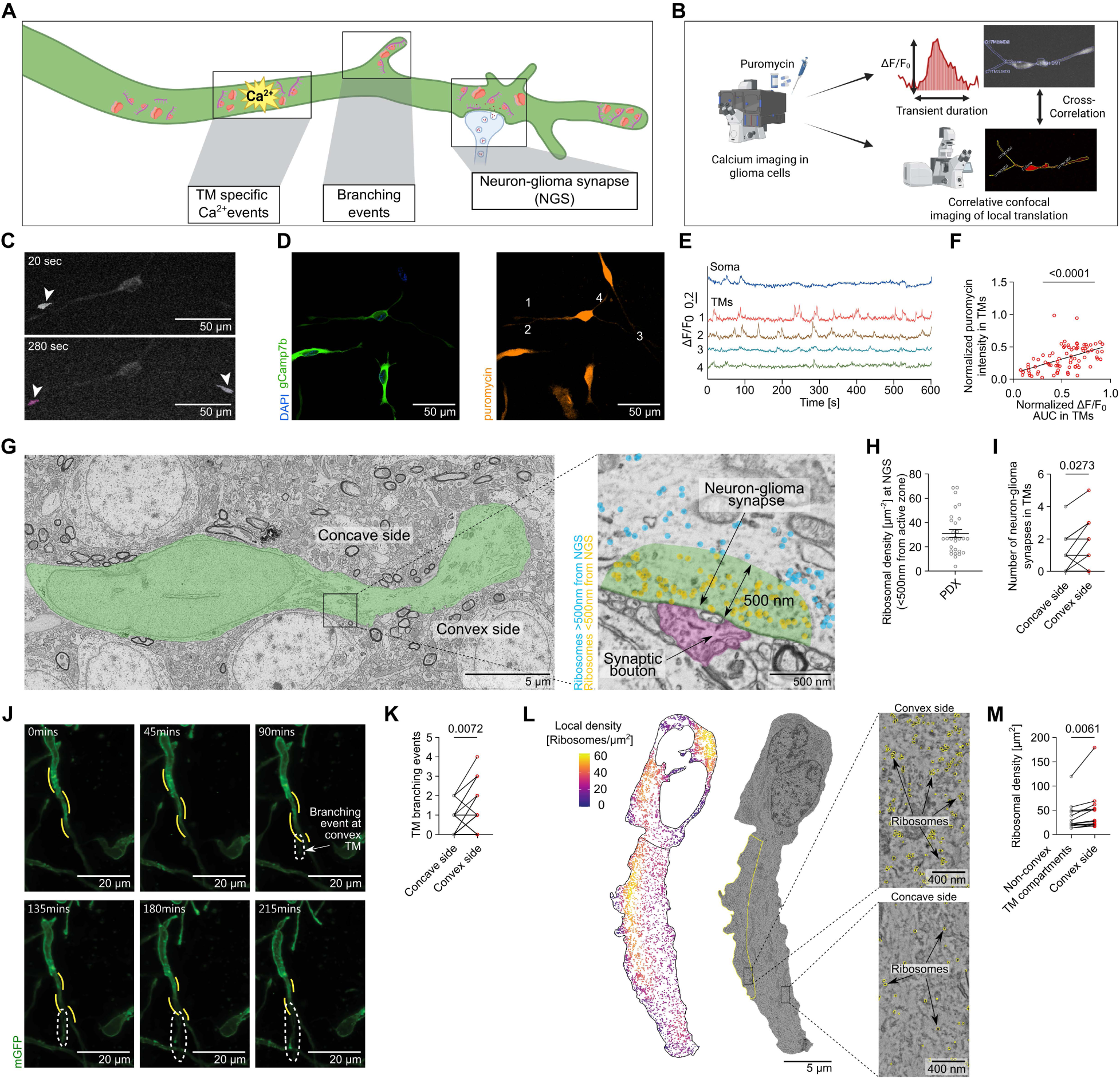
Correlation of functional activity and local translation in TMs. (**A**) Schematic illustration demonstrating the involvement of local translation in TM-localized calcium events, neuron-to-tumor synaptic connectivity and branching events. (**B**) Workflow linking calcium events to local translation through cross-correlation with confocal imaging and puromycin staining in a patient-derived glioblastoma model. (**C**) Visualization of representative Ca²⁺ events in a TM of a S24-gCamp7b tdTomato GB cell in a neuron-tumor co-culture. (**D**) Representative confocal ICC images of the GB cell in (C), stained for DAPI (blue), gCamp7b (green) and puromycin (orange), displaying nascent proteins translated during Ca²⁺ imaging. (**E**) Representative baseline-normalized ΔF/F0 fluorescence Ca²⁺ signals in soma and TMs of the GB cell shown in (C). (**F**) Correlation plot of puromycin signal intensity (normalized to mean soma signal) and Ca²⁺ event area under the curve (AUC) in TM-localized microdomains (normalized to whole-cell activity) of S24-gCamp7b tdTomato GB cells in neuron-tumor co-culture during live-cell imaging (n = 78 TMs). (**G**) Electron microscopic image of a non-invasive GB cell (green overlay) in PDX tissue with neuron-glioma synapses. Zoom-in displays a magnified view of neuron-glioma synapse. Ribosomes located in the postsynaptic region (green, <500nm distance to the synaptic cleft) are marked with a yellow overlay, ribosomes located outside this region (>500nm distance to the synaptic cleft) marked with a blue overlay. The presynaptic region is marked with a purple overlay. (**H**) Quantification of ribosomal density in the postsynaptic area indicated in (G) (n = 25 NGS). (**I**) Quantification of number of neuron-glioma synapses at the convex side of the TM compared to the concave side of GB cells in PDX tissue (n = 15 GB cells). (**J**) Representative TM branching event (indicated by dashed line) at the convex side of the TM (indicated by yellow line) of a GB cell in PDX tissue during intravital two-photon imaging over a time frame of 215 minutes (maximum intensity based projection). (**K**) Paired quantification of branching events on the convex compared to the concave sides of TMs of GB cells in PDX tissue (n = 37 branching events at n = 13 GB cells). (**L**) Representative electron microscopic image of a GB cell in PDX tissue with a bending TM curvature. On the left, density map of cytosolic ribosomes with the density coloured according to the legend. On the right, an EM image of the same cell cropped at its cell membrane, with the convex side outlined in yellow. Representative magnified views of the convex and concave regions are shown, with ribosomes highlighted in yellow. (**M**) Paired quantification of ribosomal density in the curvature-defined convex TM-regions in PDX tissue compared to the remaining TM (n = 13).

For this purpose, we combined calcium imaging with a correlative puromycin readout of the same GB cells (Fig. 4B). This analysis revealed a heterogeneous occurrence of calcium events across somata and TMs of single GB cells (Fig. 4 C-E). TMs with overall higher calcium trace AUCs exhibited higher signal intensity for puromycin (Fig. 4F), indicating a correlation between calcium signaling and local translation in TMs.

These findings raised the question of whether this coupling is spatially organized along TMs. Consistent with this idea, ultrastructural quantification revealed that TM ribosomes are preferentially localized at the cell periphery adjacent to the plasma membrane - rather than within the central core (fig. S6F-G) (69, 70) - precisely the subcellular surface most exposed to extracellular synaptic signals.

To further explore the link between functional connectivity and local translation, we investigated the distribution of NGS along TMs. Quantification of ribosomes in proximity to the synaptic cleft demonstrated a reproducible accumulation of ribosomes within the postsynaptic compartment of NGS (Fig. 4G, H) and a higher number of NGS at the dominant convex TM sides compared to concave TM segments (Fig. 4I). On the ultrastructural level, these convex TM areas displayed significantly higher ribosomal density compared to concave TM regions (Fig. 4L, M) supporting the idea that neuronal signalling through NGS affects TM sprouting and GB cell motility through spatially compartmentalized translation within TMs. In line with this finding, branching of new processes occurred more often at the convex side (Fig. 4J, K).

Taken together, these findings suggest that neuronal input through NGS is spatially coupled to local translation at defined microdomains along TMs. The convergence of postsynaptic ribosome accumulation, elevated translational activity, and preferential process branching at convex TM surfaces points to a model in which calcium-dependent signaling from NGS locally activates protein synthesis at sites primed for structural remodeling - providing a mechanistic link between neuron-tumor connectivity and the dynamic invasion of GB cells through the brain parenchyma.

### Local translation promotes cytoskeletal turnover at sites of TM invasivity

Given the observed enrichment of local translation in TMs of invasive GB cells, we next examined whether local translation affects TM development and dynamics required for cellular remodelling and cell state transition (Fig. 5A). We interrogated both bulk RNA Sequencing of TM-enriched fractions and our MERFISH dataset and revealed an enrichment of RNAs encoding cytoskeletal proteins in TM compartments, including MAP1B, ß-III-tubulin and nestin (Fig. 5B-C, Table S3) which are important molecular components for neurite protrusion (*71*) and invasive migration in cancer cells (*72*). Given that transcripts for nestin, an intermediate filament protein associated with neural stemness and poor prognosis in glioblastoma (*73*), were enriched consistently in TMs for both transcriptomic datasets (Fig. 5B-C), this strongly suggests local synthesis of cytoskeletal proteins within TMs rather than exclusive somatic production followed by protein transport.

**Figure 5.**
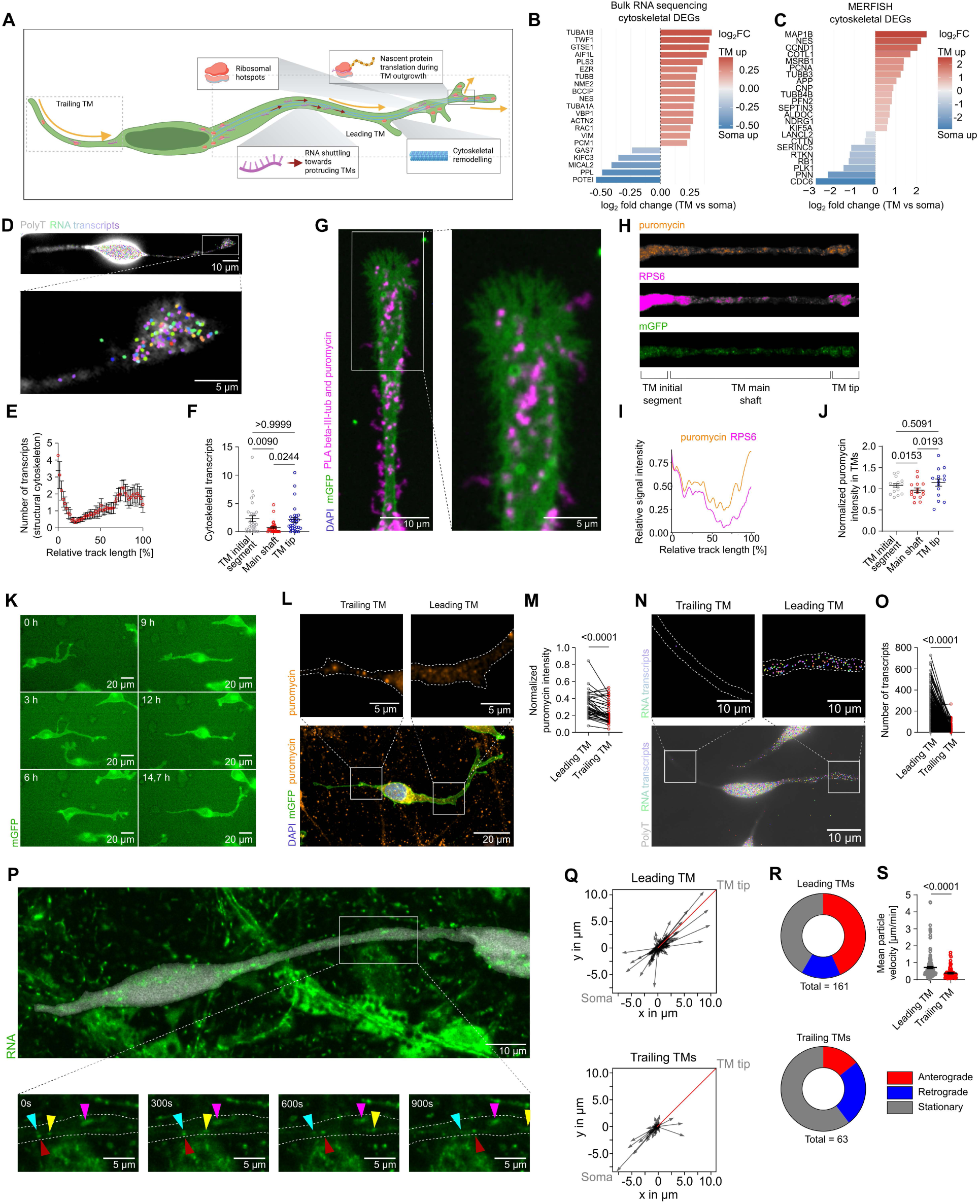
Local translation promotes cytoskeletal turnover in leading TM structures. (**A**) Schematic illustration highlighting aspects of local translation relevant for invasive glioblastoma biology: ribosomal hotspots, nascent protein translation during TM outgrowth, RNA shuttling towards protruding TMs and cytoskeletal remodelling. (**B**) Differentially expressed genes encoding for cytoskeleton-associated proteins (see Materials and methods) in the TM-enriched fraction from the bulk RNA sequencing insert assay data of S24 GB cells (Fig. 2A). Top 21 genes are displayed by their TM-vs-soma log₂–fold change (x-axis). TM-enriched genes are displayed in red, while soma-enriched genes are displayed in blue. (**C**) As in (B) but differentially expressed genes (Padj < 0.05) encoding for cytoskeleton-associated proteins in the TMs from the MERFISH data of S24 GB cells (Fig. 2A). (**D**) MERFISH image of a S24 GB cell in mono-culture with polyT signal (gray) and spatial distribution of RNA transcripts (multicolor). Zoom-in displays distribution of transcripts in a growth cone-like structure at the TM tip. (**E**) Quantification of cytoskeleton-associated transcripts along the TM of S24 GB cells in mono-culture normalized to the total TM length (n = 30 cells)(**F**) Quantification of cytoskeleton-associated transcripts in the TM initial segment, TM main shaft and TM tip of S24 GB cells in mono-culture (n = 30 cells) (**G**) Representative confocal image of the TM of a S24 GB cell in monoculture labeled for mGFP (green) and PLA of puromycin and beta-III-tubulin representing nascent beta-III-tubulin with a zoom-in on the TM tip. (**H**) Straightened mGFP (green), RPS6 (magenta) and puromycin (orange) signal intensities along the TM of a representative invasive S24 mGFP GB cell in PDX tissue. (**I**) Mean intensity profiles of RPS6 (magenta) and puromycin (orange) signal along the normalized TM length of S24 GB cells in PDX tissue (n = 15 TMs) (**J**) Quantification of soma-normalized puromycin signal intensities between the TM segments of S24 GB cells in PDX tissue (n = 15 cells). (**K**) Representative motility of a S24 GB cell (green) over the course of 14.7 hours during live cell imaging. (**L**) Post-hoc correlative ICC image of a representative S24 GB cell from (J), stained against DAPI (blue), puromycin (orange) and GFP (green) in the leading and trailing TM with a mGFP based mask of the cell outline (dashed line). (**M**) Quantification of normalized puromycin intensity in the leading TM compared to the trailing TM of GB cells in neuron-tumor co-cultures (n = 38 cells). (**N**) Representative MERFISH image of a S24 GB cell in mono-culture showing polyT signal (gray) and spatial distribution of transcripts (multicolor) in soma and leading versus trailing TM with a polyT based mask of the cell outline (dashed line). (**O**) Quantification of transcripts based on MERFISH imaging in leading versus trailing TMs of S24 GB cells in mono-culture (n = 358 cells). (**P**) Representative motility of RNA particles visualized by SYTO RNASelect Green Fluorescent Cell Stain (green) in a S24 GB cell (grey) in neuron-tumor co-culture over the course of 15 minutes during confocal live cell imaging. Zoom-ins display the motility of RNA particles (green) in the TM indicated by colored arrowheads. (**Q**) Vector plots displaying RNA particle trajectories in leading (upper, n = 163 particles) and trailing (lower, n = 63 particles) TMs of S24 tdTomato GB cells in neuron-tumor co-cultures (n = 10 cells). Each vector represents the displacement (direction and distance) of a single RNA particle in relation to the orientation of its TM, which is normalized to the orientation of the red line. (**R**) Pie charts of RNA particles classified by their motility trajectory in leading TMs and trailing TMs of S24 tdTom GB cells in neuron-tumor co-cultures. (**S**) Quantification of mean RNA particle motility in leading and trailing TMs of GB cells in (P) (n = 161 particles in leading TMs, n = 63 particles in trailing TMs).

Strengthening the hypothesis that filament associated programs are primary targets of local translation in TMs, cytoskeletal transcripts abundance was not uniformly distributed along TMs: Instead, we identified the TM initial segment positioned proximally near the soma and the distal endpoint of TMs (TM tip) as hotspots of high transcript density (Fig. 5D-F). Accordingly, we identified nascent ß-III-tubulin via proximity ligation assay (PLA) against puromycin and ß-III-tubulin in the TM tip (Fig. 5 G) as well as increased puromycin incorporation and RPS6 signal intensity in the TM initial segment and TM tip (Fig. 5H-J). Furthermore, TMs sub-compartments positive for drebrin, an actin-binding protein and cytoskeletal remodeling marker (*74–77*), showed increased puromycin signal (fig. S8G-H), indicating elevated translational activity during active cellular migration. This enrichment of enhanced local translational activity at TM tips (Fig. 5E, J) is a correlate to growth cones in developing neurites, which similarly depend on localized translation for directional extensions (*71*, *78*, *79*).

Especially invasive GB cells display a similar asymmetric TM morphology (*10*, *80*), which led us to examine if there were fundamental differences of local translation between single TMs of a cell associated with invasive behaviour. We performed correlative live-cell imaging and post-hoc quantification of puromycin incorporation after puromycin-pulsing in neuron-tumor co-cultures (Fig. 5K). Puromycin intensity analysis revealed significantly higher local protein synthesis in leading TMs of GB cells compared to trailing TMs (Fig. 5L-M, Movie S4). This pattern was consistently observed across multiple invasive GB cell lines (Fig S8 D-F). Furthermore RNA transcript density was likewise significantly increased in leading TMs relative to trailing ones (Fig. 5N-O), suggesting coordinated RNA localization and translation at the invasive front. To understand the dynamics underlying these RNA asymmetry within TMs, we examined RNA trafficking in the TMs. Live imaging using an RNA-selective fluorescent dye revealed dynamic bidirectional RNA granule transport along TMs (Fig. 5P). Notably, leading TMs displayed a higher fraction of anterograde transport toward distal compartments (Fig. 5Q-R), increased RNA particle velocity (Fig. 5S) and pronounced transcript accumulation at TM tips (Fig. 5E, Q). In contrast, trailing TMs showed relatively more retrograde RNA movement (Fig. 5R). This polarized trafficking pattern mirrors mechanisms described in developing neurites, where RNA transport toward growth cones enables spatially restricted translation programs supporting directional outgrowth (*78*, *81*).

Taken together, these data reveal that TMs harbor a spatially resolved and directionally organized gene expression program, in which cytoskeletal transcripts are selectively enriched and locally translated at sites of active structural remodeling contributing to cellular plasticity.

### Targeted disruption of local translation decreases tumor invasion and whole-brain colonization

Having established that local translation contributes to TM dynamics and outgrowth, we next aimed to identify candidate effectors mediating structural remodeling within TMs and leading to increased invasivity. Bulk RNA sequencing of TM fractions from S24 cells revealed a TM-enrichment of transcripts associated with neuronal growth cones and programs of process extensions (Fig. 6 B). Among these, *GAP43* and *GPM6A* were prominently represented within TMs, consistent with their established roles in membrane protrusion dynamics and neurite outgrowth. Sh-RNA mediated knockdown of *GAP43*, a well-established mediator of axonal growth cone dynamics and implicated in tumor cell networks (*82–86*), significantly reduced TM overall motility during intravital imaging of PDX mice (fig. S9 A, B) indicated by lower invasion speed(Fig. 6C) and a decreased number of TM branching events(Fig. 6D) supporting the hypothesis that TM invasion hijacks conserved growth-cone machinery. We next focused on GPM6A, a membrane glycoprotein implicated in filopodia formation and synaptic plasticity (*87–89*) with a potential role in TM protrusion dynamics. Colocalization of GPM6A immunolabelling and puromycin-based visualization of nascent proteins in TMs shown via a PLA and superresolution STED microscopy suggested local translation of GPM6A at TM tips (Fig. 6E-F). Functional knockdown of GPM6A achieving RNA reduction of approximately 50% (Fig. 6I) resulted likewise in reduced TM turnover and motility in neuron-tumor co-cultures (fig. S9 C-G). Next, we explored the role of GPM6A knockdowns in PDX models. Tumor growth was drastically reduced as reflected in smaller tumor areas in *ex vivo* assessments (Fig. 6G, H, J), and stagnation of tumor cell count over the time course of four weeks using longitudinal *in vivo* two-photon microscopy (Fig. 6K-M) as compared to control PDX mice. GPM6A knockdown in GB cells further displayed a reduced invasion speed and TM turnover (Fig. 6N-Q) as well as significantly decreased TM branching events (Fig. 6R-T) compared to controls indicating impaired dynamic remodeling of invasive processes. In line with previous studies defining GAP43 and GPM6A as regulators of neuronal growth cone dynamics (*87*, *88*, *90–92*), our findings reveal striking parallels between TM biology and neuronal morphogenetic programs. GPM6A emerges as a key regulator of TM plasticity as well as tumor growth suggesting that TM-enriched targets of local translation govern a functional importance for glioblastoma invasivity which can be molecular targets for therapy of intractable brain tumors.

**Figure 6.**
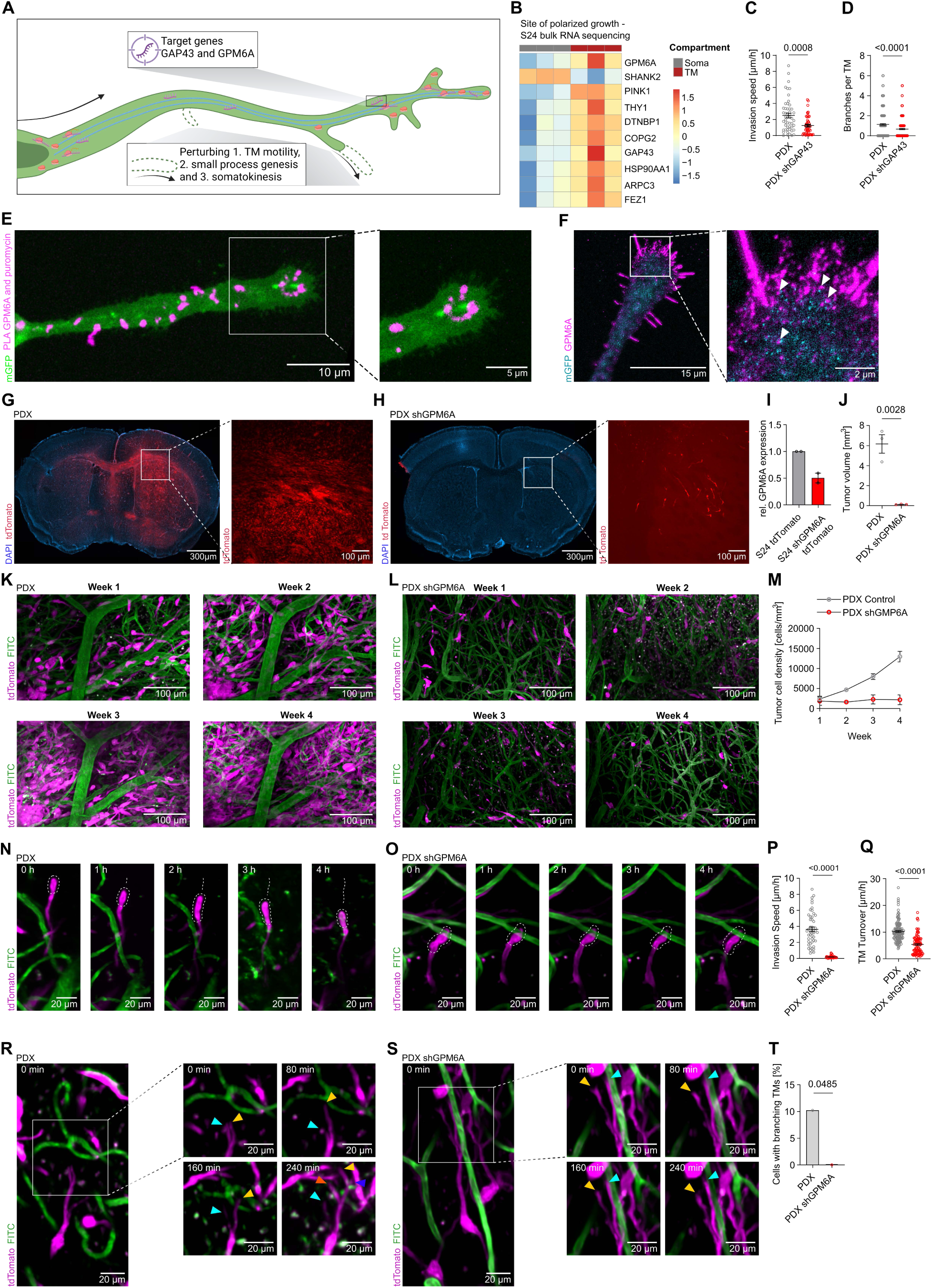
Targeting local translation perturbs TM formation and tumor cell invasion. (**A**) Schematic illustration depicting target genes involved in local translation and the functional consequences of their perturbation by genetic knockdown on TM motility, small process motility and somatokinesis. (**B**) Heatmap of variance-stabilized (vst) expression values from DESeq2 for the top 10 genes from the GO term “site of polarized growth” (GO:0030427) with the lowest adjusted p-values, ordered accordingly (top = most significant) in Soma vs TM enriched fractions of RNA bulk sequencing of S24 GB cells. Expression values are scaled per gene (row Z-score), and colours represent relative expression in the corresponding samples. (**C**) Quantification of invasion speed over a time frame of 24 hours during intravital two-photon imaging from fig. S9A-B of S24 GB cells compared to S24 shGAP43 GB cells in PDX tissue (n = 31 S24 GFP cells, n = 51 S24 shGAP43 GFP cells). (**D**) Quantification of branches per TM over a time frame of 24 hours during intravital two-photon imaging from fig. S9A-B in S24 GB cells compared to S24 shGAP43 GB cells in PDX tissue (n = 31 S24 GFP cells, n = 51 S24 shGAP43 GFP cells) (**E**) Representative confocalimage of the TM of a S24 GB cell in mono-culture labeled for mGFP (green) and PLA of GPM6A and puromycin representing nascent GPM6A proteins (magenta)with zoom-in on the TM tip.. (**F**) Representative STED image of the TM tip of a S24 GB cell in mono-culture labeled for puromycin (cyan) and GPM6A (magenta) with a zoom in on the outgrowing edge. Arrowheads highlight colocalization of nascent proteins (puromycin) and GPM6A. (**G**) Representative overview of PDX brain slices, stained for DAPI (blue) and tumor cells (red) eight weeks after striatal injection of S24 tdTomato GB cells. Zoom-in displays the native tumor cell signal. (**H**) Representative overview of PDX brain slices, stained for DAPI (blue) and tumor cells (red) eight weeks after striatal injection of S24 shGPM6A tdTomato GB cells. Zoom-in displays the native tumor cell signal. (**I**) Relative abundance of GPM6A RNA in S24 shGPM6A tdTomato cells measured by qPCR normalized to housekeeping gene abundance and S24 tdTomato control cells. (n = 2 biological replicates with n=3 technical replicates each) (**J**) Quantification of tumor volume of S24 compared to S24 shGPM6A PDX models eight weeks after tumor injection (n = 3 S24 control brains, n = 3 S24 shGPM6A brains). (**K**) Representative intravital two-photon imaging of tumor growth in the same region over 4 weeks of S24 tdTomato GB cells (magenta) in PDX, combined with an angiogram (green) (post-processed with enhance.ai in NIS Elements for representation). (**L**) Representative intravital two-photon imaging of tumor growth in the same region over 4 weeks of S24 shGPM6A tdTomato GB cells (magenta) in PDX, combined with an angiogram (green) (post-processed with enhance.ai in NIS Elements for representation). (**M**) Quantification of tumor volume over 4 weeks of S24 tdTomato PDX compared to S24 shGPM6A tdTomato PDX models based on intravital imaging (n = 3 regions from one S24 shGPM6A tdTomato PDX, n = 3 regions from one S24 tdTomato PDX). (**N**) Representative somatic movement (indicated by dashed lines) of S24 tdTomato GB cells (magenta) in PDX tissue over a time frame of 4 hours during intravital two-photon imaging (maximum intensity-based projection) combined with an angiogram (green) (post-processed with enhance.ai in NIS Elements for representation). (O) Representative arrest of somatic movement (indicated by dashed lines) in S24 shGPM6A tdTomato GB cells (magenta) in PDX tissue over a time frame of 4 hours during intravital two-photon imaging (maximum intensity-based projection) combined with an angiogram (green) (post-processed with enhance.ai in NIS Elements for representation). (**P**) Quantification of invasion speed over the time frame of 4 hours during intravital two-photon from from (M) and (N) in S24 tdTomato and S24 shGPM6A tdTomato cells in PDX mice (n = 49 GBC from one S24 tdTomato PDX vs. n = 31 GBC from one S24 shGPM6A tdTomato PDX). (**Q**) Quantification of TM turnover over the time frame of 4 hours during intravital two-photon from (M) and (N) S24 tdTomato cells compared to S24 shGPM6A tdTomato cells in PDX mice (n = 131 TMs from n = 50 GBC in one S24 tdTomato PDX vs. n = 93 TMs from n = 29 GBC in one S24 shGPM6A tdTomato PDX). **(R)** Representative TM dynamics of S24 tdTomato GB cells (magenta) in PDX tissue over a time frame of 4 hours during intravital two-photon imaging (maximum intensity-based projection) combined with an angiogram (green) Zoom-ins display the TM genesis and dynamics (indicated by colored arrowheads) (post-processed with enhance.ai in NIS Elements for representation). **(S)** Representative TM dynamics of S24 shGPM6A tdTomato GB cells (magenta) in PDX tissue over a time frame of 4 hours during intravital two-photon imaging (maximum intensity-based projection) combined with an angiogram (green) Zoom-ins display the TM dynamics (indicated by colored arrowheads) (post-processed with enhance.ai in NIS Elements for representation). **(T)** Quantification of cells with branching TMs over a time frame of 4 hours during intravital two-photon imaging (Q) and (R) of S24 tdTomato cells and S24 shGPM6A tdTomato cells in PDX mice (n = 131 TMs from n = 50 GB cells in one S24 tdTomato PDX vs. n = 93 TMs from n = 29 GB cells in one S24 shGPM6A tdTomato PDX, tested by Chi-squared test).

It is becoming increasingly clear that invasive glioblastoma cells exploit fundamental neuronal mechanisms to drive brain infiltration (*8*, *10*, *63*, *67*, *93–95*). The findings presented here established that glioblastoma cells have co-opted the neural mechanism of local translation (*46*) in TMs to achieve rapid and spatially restricted invasion and colonization through the brain. By integrating ultrastructural analyses, spatial transcriptomics, and functional translation assays, we revealed that all canonical hallmarks of local translation - ribosomal presence, mRNA transport, and active protein synthesis are present in invasive glioblastoma TM (*96*, *97*). While neurons and glial cells utilize local translation for diverse functions including axon guidance, myelination, and synaptic plasticity (*34*, *46*, *98–105*) glioblastoma cells have repurposed this machinery for driving molecular heterogeneity and whole-brain invasion. Taken together, this study establishes local translation as a fundamental mechanism driving GB brain invasion and GB cell heterogeneity.

## Acknowledgments

We gratefully acknowledge the data storage service SDS@hd, supported by the Ministry of Science, Research and Arts Baden-Württemberg (MWK). During the preparation of this manuscript, the authors used ChatGPT 3.5, 4.0,5.0, 5.1 and 5.2 (OpenAI), Claude Sonnet 4.5 (Anthropic) and Composer 1.5 (Cursor) for coding support and for improving the clarity and readability of the text. The authors reviewed and edited the content as needed and take full responsibility for the content of the published article.

We thank Manuela Brom, Damir Krunic, Marko Lampe and Felix Bestvater from the Light Microscopy Core Facility of the German Cancer Research Center and Karsten Richter of the EM Core Facility of the German Cancer Research Center for their support. We thank F. Blum, T. Rubner, M. Eich, K. Hexel, and S. Schmitt from the DKFZ Flow Cytometry Core Facility for their help with FACS experiments. Further, we acknowledge the support of the Nikon Imaging Center Heidelberg Core facility by Ulrike Engel and Christian Ackermann. For support in project associated laboratory work we thank Andrea Rojas-Martínez, Verena Buchert, Daniela Pimonov and Marvin Porsiel. We thank Niklas Schwarz (Hertie-Institut Tübingen) for providing human CSF for the human organotypic slice cultures. Finally, we thank K. Selke, M. Kempfert, K. Dell, A. Riedasch, K. Schmidt, A. Berdel and K. Becker for their support with animal care and the design of animal experiments.

## Funding

European Center for Neuro-Oncology, EZN (VV)

German Research Foundation DFG: VE1373/2-1 (VV)

Else Kröner-Fresenius-Stiftung: 2020-EKEA.135 (VV)

Physician-Scientist-Program, Krebs- und Scharlachstiftung, Medical Faculty of Heidelberg University (VV)

Schwiete Stiftung (VV)

Wilhelm-Sander-Stiftung (VV)

Aventis Bridge Foundation (VV)

Research Seed Capital (RiSC) from the Ministry of Science (project number VE1373/2-1516), Research and the Arts Baden Württemberg and the Health + Life Science Alliance Heidelberg Mannheim (VV)

Deutsche Krebshilfe/German Cancer Aid (Mildred-Scheel-Scholarship for MD students) (E.R., S.K.T., Y.Y., V.K.S., R.S.D.)

German Research Foundation (DFG: SFB 1389, project ID 404521405) (V.V., N.L., Y.Y., S.K.T., A.P.L. and E.R.)

Heidelberg University Medical Faculty Clinician Scientist Program (O.T.H.)

## Author contributions

Conceptualization: NL, NW, VV Data curation: NL, NW, VV

Software: NL, NW, MPD, MFL, MYL, MCS, JM, RDi, DHH, VV

Validation: NL, NW, VV Methodology: NL, NW, VV

Investigation: NL, NW, MPD, MFL, MB, TW, MYL, JL, ER, MCS, SKT, NS, YT, JM, YK, TK, RLP, SJS, YvY., YaY., RD, APL, OTH, RD, MiB, VKS, AJ, DHH and VV

Visualization: NL, NW, MPD, MFL, MYL, MB, TW, ER, MCS, YT, DHH and VV

Formal analysis: NL, NW, MPD, MFL, MB, TW, MYL, JL, ER, MCS, SKT, YT, RLP, YvY, AJ, DHH and VV

Resources: TK, BD, AdC, TVW, BS, AJ, DT, FS, DHH and VV

Funding acquisition: VV

Project administration: NL, NW, VV Supervision: VV

Writing – original draft: NL, NW, MPD, MFL, MB, ML and VV Writing – review & editing: NL, NW, BD, DHH, ES and VV

## Data, code, and materials availability

This study did not generate new unique reagents. Materials described in this paper are available from the corresponding author upon reasonable request. Code used for ultrastructural ribosomal detection analysis is available at https://github.com/venkataramani-lab/. Any additional information required to reanalyze the data reported in this paper is available from the lead contact upon request.

## Materials and Methods

### Animal models

All animal procedures were performed in accordance with the institutional laboratory animal research guidelines following approval of the Regierungspräsidium Karlsruhe, Germany. Efforts were made to minimize animal suffering and reduce the number of animals used according to the 3R principles. Male NMRI nude mice older than 8 weeks (sourced from Charles River and Janvier) were used for all animal studies involving patient-derived glioblastoma models. Mice were routinely checked for clinical endpoint criteria and if they showed marked neurological symptoms or weight loss exceeding 20%, experiments were terminated. No maximum tumor size was defined for the invasive brain tumor models.

Cranial window implantation in mice was performed by incorporation of a custom-made titanium or teflon ring to allow for painless head fixation during imaging. Tumor cell implantation involved the stereotactic injection of 100,000 tumor cells into the cortex or striatum of the mouse brain at an approximate depth of 500 µm. In some cases, tumor cell injection was performed concurrently with cranial window implantation in a single procedure or without any cranial window implantation at all.

Following 3-4 months of tumor growth, xenografted mice were anesthetized with an intraperitoneal barbiturate overdose and transcardially perfused with 4% PFA for non-puromycin based readouts and postfixed overnight at 4°C. Following fixation, brains were sliced into 80 μm thin sections using a vibratome (VT1200S, Leica Germany). For puromycilation assays, animals were intracardially perfused with carbogenized ice-cold NMDG-ACSF (in mM: NMDG 135, KCl 1, KH_2_PO_4_ 1.2, MgCl_2_ x 6 H_2_O 1.5, CaCl_2_ x 2 H_2_O 0.5, Choline Bicarbonate 20, Glucose 12.95), decapitated and brains were immediately removed. Acute coronary brain slices were generated using a vibratome (VT1200S, Leica Germany) with 250-350 μm thickness. Immediately following cutting, slices were incubated in 50 μM puromycin in carbogenized NMDG-ACSF for 2 minutes or 10 minutes at room temperature to minimize the stress experienced by tumor cells and to ensure that puromycin binding occurred locally by limiting its time for diffusion. After puromycin incubation, the slices were fixed in 4% PFA overnight.

### Human tissue models

Human material used for experimental investigation was obtained after approval of the local regulatory authorities (ethical codes 23-1233-S1, 23-1234-S1, S-005/2003, 23-1175-S1 and 057/2021BO2) after pseudonymization. Human neocortical tissue was freshly resected from diagnosed glioblastoma, low-grade glioma or brain metastasis patients. Pathological areas were classified as access cortex, invasion zone or tumor core based on MRI tumor signatures and extracted using Brainlab neuro-navigation system. Resected tissue was submerged in 4% PFA for immediate fixation overnight at 4°C or in carbogenated, ice-cold NMDG-aCSF for acute or cultured human slice preparation. Fixed tissue blocks were subsequently transferred to PBS and cut into 80 μm thin sections using a vibratome (VT1200S, Leica Germany). Fresh tissue was instead cut into 250-300 μm thin sections after microsurgical removal of the arachnoid using a vibratome (Leica Germany) constantly oxygenated before incubation in 50µM puromycin for 10 minutes and subsequent fixation (invasion zone, tumor core). For human brain slice cultures, access cortex sections were cultured on Millicell membrane inserts (Merck) with human cerebrospinal fluid as culture medium purified from hydrocephalus patients (*45*). Culture medium was exchanged twice a week. For inoculation, target cells were prepared as previously described 24 hours after slice culturing, undergoing accutase incubation, centrifugation, and resuspension in hCSF at 20.000 cells/μl. Cells were inoculated into tissue sections using a 10 μl Hamilton syringe to deliver 0.5 μl cell suspension at a 90° angle.

### Cell culture models

Primary tumor cell lines derived from resected glioblastoma specimens were cultured as previously described (*8*, *16*, *20*, *67*) using DMEM/F-12 medium in a serum-free, non-adherent environment designed to support ‘stem-like’ growth. This medium included B27 supplement without vitamin A, insulin, heparin, epidermal growth factor, and fibroblast growth factor. DNA methylation profiling at over 850,000 CpG sites was performed regularly on all GB cell lines using the Illumina Infinium Methylation EPIC array kit, following the manufacturer’s instructions at the Genomics and Proteomics Core Facility of the German Cancer Research Center, Heidelberg, Germany, or the Institute for Neuropathology, Heidelberg, Germany. Glioblastoma stem-cell lines were transduced with lentiviral vectors carrying membrane-bound GFP (pLego-mGFP) or tdTomato (pLego-T2) for labeling, pLego-T2-GCaMP7b-tdTomato for calcium imaging or shRNA carrying lentiviral constructs. Transduced cells were routinely sorted by fluorescence-activated cell sorting (FACS) using either a FACSAria Fusion 2 or FACSAria Fusion instrument. GFP was detected using a BL 530/30 filter, while tdTomato was detected using a YG 586/15 filter. GB cell lines and their molecular diagnosis used within the experiments of this study are listed in table S1.

### Neuron-tumor co-cultures and GB cell mono-cultures

Primary rat cortical neuronal cultures were generated as described before (*64*), E19.5 rat embryos were harvested (62) from pregnant Wistar rats. The skull was opened, the brain taken out and the meninges removed. Subsequently, cortices were dissected, cut in small pieces and incubated with 0.5% trypsin for 20 minutes. For further separation, cells were gently triturated with 20G and 25G needles before passing the cell suspension through a 100 µm pore filter mash. Primary cultures were seeded on poly-L-lysine coated glass coverslips or glass bottom plates at 55.000 cells per cm^2^ in DMEM high glucose medium supplemented with 10% FBS and 250mM L-glutamine. 6 hours after seeding medium was exchanged to neurobasal medium supplemented with B27 and 150mM L-glutamine. Cortical cultures were used from 7 days after initial cell seeding for neuron-tumor co-cultures.

Neuron-tumor co-cultures were generated as described before (*8*, *10*): GB cells were dissociated by accutase incubation at 37°C for 5 minutes. For each cell line, 1000 cells were seeded per well onto 24-well plates containing glass coverslips or a glass bottom optical plate for live cell imaging. For 96-well glass bottom optical plates 250-500 cells were seeded per well instead based on the experiment. Cultures were held for at least 5 days before imaging or fixation. Tumor cell mono-cultures were instead cultured on Matrigel-coated surfaces for at least 7 days before further processing, until the formation of TMs was observed. For puromycilation assays, cells were treated with 5 µM puromycin and incubated at 37°C for 10 minutes. Anisomycin was added at 10 µM for 10 minutes before puromycin incubation as negative control in some samples. 10 minutes after puromycin treatment, the medium was removed and 0.5 ml of 4% PFA were added to each well for fixation. After 10 minutes at room temperature, PFA was removed, and wells were briefly rinsed with 0.5 ml PBS. Subsequently, 1 ml PBS was added to each well, and plates were sealed with parafilm and stored at 4 °C until further processing. For metabolic labeling of nascent proteins, cells were incubated in methionine-free DMEM high glucose (DMEM high glucose w/o Met, Cys, L-Glut, Sodium Pyruvate; supplemented with 31.5 mg/l Cysteine, 25 mg/l Sodium Pyruvate, 0.5 mM Glutamine and 20 µl/ml B 27 + Vit.A) supplemented with 200 µM AHA (4-Azido-L-homoalanine HCl) for 5–10 min at 37 °C. Following the pulse, cells were washed once with PBS and fixed with 4% PFA for 10 min at room temperature (RT). Fixed samples were washed twice with PBS.

### In vitro live-cell time-lapse imaging of neuron-tumor co-cultures

S24 mGFP, T269 mGFP, and P3XX mGFP GB cells were used in this experiment. 5-7 days after seeding on neuronal cultures, tumor cells were imaged via an inverted Nikon Ti2 microscope equipped with a 20x NA 0.75 multi immersion objective every 10 minutes over the course of 14 hours maintaining plate focus via the Nikon perfect focus system. Temperature was maintained at 37°C and 5% CO_2_ by a Tokai Hit incubation chamber. Immediately following imaging, cells were incubated for 10 minutes in 5 µM puromycin in the culture medium. Afterwards the medium was removed and cells were fixed with 4% PFA for 10 minutes.

### In vivo multiphoton laser scanning microscopy (MPLSM) and image analysis

#### *In vivo* multiphoton imaging

*In vivo* MPLSM imaging commenced three weeks post-implantation using a Zeiss 7MP microscope, a Zeiss LSM 980 with Airyscan2, and a TriM Scope II microscope (LaVision BioTec GmbH), all equipped with pulsed Ti:Sapphire lasers (Chameleon II ultra; Coherent). Imaging of GFP, tdTomato, FITC-dextran and TRITC-dextran was performed at excitation wavelengths of 850 nm and 960 nm, respectively. Emitted fluorescence was separated using a 560 nm dichroic mirror. For the Zeiss 7MP and Zeiss LSM 980 systems, bandpass filter sets of 500–550 nm and 575–610 nm were used. The TriM Scope II system utilized a 500–570 nm bandpass filter and a 590 nm longpass filter. Imaging with the Zeiss setups was conducted using a 20x, 1.0 NA, apochromatic water immersion objective with a 1.7 mm working distance. The TriM Scope II setup included a 16x, 0.8 NA, apochromatic objective with a 3 mm working distance, and a 25x, 1.1 NA, apochromatic objective with a 2 mm working distance (both Nikon). Fluorescence signals were detected using low-noise, high-sensitivity photomultiplier tubes. Anesthesia was induced using 3–5% isoflurane in 100% oxygen and maintained at 0.5–3%, with levels adjusted based on the mouse’s respiratory rate. Eye cream was applied after induction to prevent corneal drying. Throughout imaging, mice were maintained at 37°C using a temperature-controlled heating plate and monitored via a temperature sensor. For visualization of blood vessels, FITC-dextran (2,000,000 g/mol; fluorescein isothiocyanate-dextran) or TRITC-dextran (500,000 g/mol; tetramethylrhodamine isothiocyanate-dextran) was prepared at 10 mg/ml in 0.9% NaCl. Immediately prior to imaging, 100 µl of the solution was injected into the lateral tail vein. Image timelapse stacks were acquired either every 7-8 days (tumor cell density), every 12 hours (correlative 2P-EM data), every 8 hours up to 24 hours (shGAP43 somatokinesis), or every 5 minutes for up to 4 hours total (TM and small process turnover, cell somatokinesis data of main TM curvature).

To analyze tumor growth and invasion in S24 shGPM6A tdTomato compared to S24 tdTomato control, *in vivo* 3D stacks were acquired in the same field of view (FoV; FoV = 606.09 µm x 606.09 µm) every 7-8 days over 4 weeks (0.59 µm pixel size, 2.05 µs pixel dwell time), resulting in a total of 4 time points. 3 different regions per mouse were imaged, selected by similar tumor cell density at the first time point. The laser power was chosen at 90% of the optimal amount according to the built-in range indicator function of ZEN (Carl Zeiss Jena) to reduce laser power and avoid phototoxicity. After imaging, mice were observed until awake.

The 4 stacks for each time point were hyperstacked and manually registered in ImageJ/Fiji (*106*) via the vessel signal.

To analyze invasion speed, TM turnover and branching events in S24 shGPM6A tdTomato compared to S24 tdTomato control, a FoV with a volume of 606.09 µm x 606.09 µm x 100 µm (0.59 µm pixel size, 0.77 µs pixel dwell time) was selected and the image stack was repetitively imaged over 4 hours every 5 min resulting in a total of 48 time points. Laser power was chosen as described above. Visible drift during time-lapse imaging was corrected manually after image acquisition of each stack in xyz directions, if necessary. A mean respiratory rate of 90-100/min was aimed for and the respiratory rate was monitored after each stack. The isoflurane concentration was adapted accordingly. After imaging, mice were observed until awake.

The 48 stacks for each time point were hyperstacked and registered in ImageJ/Fiji via the vessel signal and by using the plugin Correct 3D drift (*107*).

#### *In vivo* image analysis and quantification

The tumor cell density was analyzed after image registration by defining a fixed volume (approx. 520µm x 520µm x 50µm) for each region imaged over 4 weeks. Cells were counted manually at each time point within this volume and expressed as tumor cell density (cells per mm^3^).

Cells were cropped and registered as described above. For each cell, a maximum intensity projection (MIP) of the z-stack, encompassing its entirety, was used for further analysis.

Measurement of somatokinesis was performed in ImageJ/Fiji. The cell somata were outlined and the center point of the selection was determined using the centroid function. The distance between the center points at time point 0 and 4 hours was divided by the experimental observation time to calculate the invasion speed.

Measurement of TM turnover was performed by manual tracking of TM tips over time in MTrackJ (*108*). Subsequently, the distances between each time point of these tracks were extracted, added up and divided by the experimental observation time to determine TM turnover per single TM, summarizing the dynamics of single TMs.

For the quantification of TM branching events, the fraction of TM tracks initiating after the first time point was calculated relative to the total number of TMs. To compare the surface turnover in somata and TMs, three-dimensional masks generated via pixel-classifier in Ilastik were manually segmented, separated by cell compartments and quantified over time in Arivis.

For visualization, image restoration for *in vivo* imaging data was performed using the enhance.ai tool within the Nikon NIS-Elements AR Analysis software v5.42.03 (Nikon GmbH Germany/Laboratory Imaging) with a pretrained model (*10*).

### Live cell imaging of RNA dye stained neuron-tumor co-cultures

5-7 days after seeding on neuronal cultures, cells were labeled with 500 nM SYTO ® RNASelect™ Green Fluorescent Cell Stain at 500 nM concentration in prewarmed neurobasal medium supplemented with L-glutamin and B27: Cell culture medium was removed from the wells designated for staining and transferred to a separate 24-well plate for temporary storage. 0.5 ml of the prewarmed labeling solution (500 nM) was added to each well and incubated at 37°C for 20 minutes. The labeling solution was removed and rinsed twice with PBS, then the original cell culture medium from the other 24 well plate was reapplied. Prior to live cell imaging, cells were recovered for at least one hour in the incubator at 37°C. Live cell imaging of RNA motility was performed with a pixel size of 0.13 µm and a z-step size of 0.7 µm every 15 seconds with a 63x oil immersion objective (NA = 1.4) at a Zeiss 780 spinning disc confocal microscope or with a pixel size of 0.07 µm without z-stacks every 10 seconds with a 40x water immersion objective (NA = 0.7) and a 1.5x optical zoom at an inverted Nikon A1R, alternating between excitation with a 488nm and 561nm laser diode.

RNA particles visible for more than five consecutive time frames were tracked using the mTrackJ (*108*) plugin in Fiji. The resulting trajectory coordinates were analyzed in Python to compute displacement vectors between the first and last positions of each trajectory. To define the TM direction, a TM axis vector was generated from the center of the soma to the tip of the corresponding TM (leading or trailing). All TM axis vectors were then rotationally aligned such that they were oriented toward a 45° reference vector (normalized vector [1, 1]). The required rotation angle was derived from the dot and cross products between the TM axis vector and the reference vector. For each TM axis vector, the corresponding rotation matrix was applied to all RNA particle vectors associated with that TM.

Aligned vectors were visualized in Matplotlib, with the reference vector shown in red and RNA particle displacement vectors shown in black, preserving their relative magnitudes and orientations.

RNA particles with a displacement vector smaller than 1 µm were considered stationary. All particles with a displacement vector larger than 1 µm were manually classified as either anterograde (moving away from the soma toward the TM-tip) or retrograde (moving away from the TM tip toward the soma).

### Puroswitch assay

Neuron-tumor co-cultures in 24-well optical plate were pre-incubated with a photoswitchable puromycin (puroswitch 10 µl) (*43*) on DIV5-7 for 30 minutes before imaging. For spatial illumination of TMs a FRAP set-up built on an inverted Nikon Ti2 microscope equipped with a 20x multi immersion objective (NA = 0.75) and an optical zoom of 1.5 (30x magnification in total) was used. Cultures were maintained within 37°C and 5% CO2 over the course of the experiment. A list of points was generated, each centered on the TM of a different GB cell. Every 10 minutes for every point a picture was taken (560 nm excitation), the activation zone (65 µm*65 µm) was illuminated with an UV-laser (405 nm) for 10 seconds to isomerize the incorporated photoswitchable puromycin towards the active form and an outer safety margin with a thickness of 65µm was subsequently illuminated with a 560 nm-laser for 10 seconds to instantly deactivate diffused activated puromycin. This activation-deactivation-process was repeated every 10 minutes for 6 hours.

For photoactivation in human organotypic slices, slices were removed carefully from the membrane insert by applying a stream of hCSF from the side with a microliter pipette. Cultures were preincubated in 10 µM puroswitch and imaged in hCSF with a spinning disc confocal built on an Nikon Ti2 inverted microscope equipped with an Nikon Plan Apo VC 20x objective (NA = 0.75) and a digital mirror device coupled to a Spectra LED laser diode box instead. Stimulated TMs were selected manually as a combined region of interest (ROI) and illuminated for three seconds with 395 nm, followed by an illumination with 550 nm for three seconds with the inverted ROI. This activation-deactivation-process was repeated every 10 minutes for 6 hours.

To visualize puroswitch uptake and incorporation into neurons and GB cells, large fields of view were illuminated for 6 hours with 395 nm for three seconds every 10 minutes instead without inactivation.

### Correlative in vitro calcium imaging in tumor microtubes

5000 S24 gCamp7b-tdTomato GB cells were seeded onto DIV7 primary rat cortical neuronal cultures on glass coverslips in neurobasal medium as described above. After 6-10 days, dishes were imaged with an inverted Zeiss LSM780 spinning disc confocal microscope at 37°C, 5% CO_2_. For imaging of calcium transients, images were acquired every second with a 20x air objective (NA = 0.8) alternating between excitation with a 488nm and 561nm laser diode at a Zeiss 780 spinning disc confocal microscope. Images were acquired with a pixel size of 0.4 µm. After an initial baseline recording period of 5-10 minutes, 5 µM puromycin was added for 10 minutes during continuous image acquisition. Cells were immediately fixed afterwards with 4% PFA for 10 minutes.

### Cell compartment-specific RNA bulk sequencing via an insert assay

Patient-derived GB cells were cultured as described above. Prior to seeding, inserts were coated with poly-L-lysine for 1h at 37°C and filled with neurobasal medium supplemented with 0.25% (v/v) glutamine and 2% (v/v) B27. 2 ml of medium were added to both the inside and outside of the inserts. Approximately 600.000 cells were seeded per insert. Seeded inserts were maintained for up to 4 weeks at 37°C and 5% CO_2_, allowing sufficient infiltration to the second compartment through the membrane by tumor microtubes. Three biological replicates were generated for sequencing with additional samples for imaging. For RNA harvesting of the soma-enriched compartment, the TM-enriched side was removed by scraping the bottom insert surface with sterile swabs; for the TM-enriched compartment, the topinsert surface was scraped. Material was transferred into pre-chilled tubes containing ceramic beads and 1 ml TRIzol reagent. RNA extraction was performed using the NucleoSpin RNA kit (Macherey-Nagel) according to the manufacturer’s instructions, samples with too low RNA concentration were excluded from further processing. RNA concentration was measured using Nanodrop or Qubit and RNA integrity was assessed with the TapeStation (Agilent Technologies). Libraries were sequenced on a NextSeq 550 platform with single-end 75bp reads.

### Quantification of GPM6A knockdown efficiency via qPCR

Total RNA was isolated from 1×10⁶ cells of S24 tdTom and S24 shGPM6A cultures using the NucleoSpin RNA Plus Mini Kit (Macherey-Nagel), which includes genomic DNA removal. RNA quantity and integrity were assessed using an Agilent 4200 TapeStation with a High Sensitivity RNA ScreenTape (Agilent Technologies). Quantitative RT–PCR was performed using the iTaq Universal SYBR Green One-Step Kit (Bio-Rad) on a LightCycler 480 instrument according to the manufacturers’ instructions. GPM6A expression was quantified using gene-specific primers (fwd: AATTCCGTGTACCAGATTCTACT, rev: CCCCAGGCATTTGATACAGC. Expression levels were normalized to the reference genes YWHAZ (primer sequences fwd: ATACTGGTTTGTCCTGGCGC, rev: ATGTGAACCGTTTCTGCCCT), TBP (primer sequences fwd: GAGCTGTGATGTGAAGTTTCC, rev: TCTGGGTTTGATCATTCTGTAG), and PPIA (primer sequences fwd: GTTCTTCGACATTGCCGTCG, rev: CTCAGTCTTGGCAGTGCAGA), and relative expression was calculated by 2^−ΔΔCt^. Analyses were performed on 2 biological replicates with 3 technical replicates each.

### Lentiviral vector generation

To generate lentiviruses expressing a shRNA against GPM6A, we sub-cloned a shRNA based knockdown cassette against the human transcript (shRNA sequence: CAGATGTGTGAGCGCTTGGTTGTGAAGCCACAGATGAACCAAGCGCTCACACATCTG) created by de novo-synthesis (Eurofins genomics) via restriction cloning into the lentiviral pLego-tdTomato expression system. ShRNA expression was controlled by the ubiquitous U6 promoter while tdTomato expression was controlled via a FSSV promoter. Viral particles were produced in HEK293 LentiX strain (Takara) in DMEM high glucose culture medium supplemented with 10% FBS and purified after cell lysis by precipitation by the PEG-it kit (System Biosciences) according to the supplier’s protocol. S24 cells tranduced with lentiviral construct for knockdown of GAP-43 was kindly provided by Prof. Winkler and prepared as described before (*8*).

### Proximity Ligation Assay (PLA)

Newly synthesized GPM6A and βIII-tubulin were visualized using a proximity ligation assay (PLA). Cells were labeled with puromycin (5 µM) for 10 min to mark nascent proteins. Following labeling, cells were fixed with 4% paraformaldehyde (PFA) for 10 min, subsequently permeabilized and processed according to the suppliers protocol.

IN brief, samples were blocked with Duolink blocking buffer for 60 min at 37 °C. Primary antibodies were diluted in Duolink antibody diluent and applied for 1 h at room temperature while gentle shaking. The following primary antibodies were used: anti-puromycin (mouse, 1:500) in combination with either anti-GPM6A (rabbit, 1:250) or anti-βIII-tubulin (rabbit, 1:200).

After primary antibody incubation, samples were washed twice with Wash Buffer A and incubated with PLA probes (anti-mouse PLUS and anti-rabbit MINUS; diluted 1:5) for 1 h at 37 °C. Ligation was performed by incubating the samples with ligation solution containing connector oligonucleotides and T4 DNA ligase for 30 min at 37 °C.

Following ligation, samples were washed in Wash Buffer A and subjected to rolling circle amplification using Phi29 DNA polymerase and fluorophore-labeled detection oligonucleotides (Duolink Detection Reagents Orange) for 100 min at 37 °C. Samples were then washed twice with 1× Wash Buffer B and once with 0.01× Wash Buffer B.

Finally, coverslips were mounted using Duolink PLA mounting medium containing DAPI.

Imaging was performed using a Leica spinning disk confocal microscope equipped with a 40× water immersion objective (numerical aperture [NA] = XX)

### Immunocytochemistry

To permeabilize the cells, 0.5 ml of 0.2% Triton X-100 in PBS was added to each well, and incubated at room temperature (RT) for 10 minutes while shaking. For blocking 0.5 ml of 10% FBS in PBS was added at RT for 10 minutes also shaking. After that 250 µl of primary antibodies (diluted in 10% FBS) were added to each well. Primary antibodies - anti-puromycin (mouse, 1:500), anti-RPS6 (rabbit, 1:200), anti-GFP (chicken, 1:1000), anti-RFP (guinea pig, 1:500), anti-nestin (mouse, 1:2000) - were diluted as indicated. After one hour of shaking incubation at RT, the antibody solution was removed, and wells were washed twice with 0.5 ml of PBS for five minutes each. Then, the secondary antibodies (diluted in 10% FBS) were added at a 1:500 dilution and incubated shaking at RT for one hour. To prevent photobleaching, samples were covered with aluminum foil immediately upon addition of the secondary antibodies and remained covered for the next steps. After one hour, samples were washed with 0.5 ml PBS for five minutes, repeating three times. Samples were stained with DAPI (1:10.000 [w/v] in PBS) followed by mounting with SlowFade^TM^ gold, covered with a cover glass and sealed with nail polish.

### STED microscopy

Super-resolution imaging was performed using STED microscopy on a Leica STELLARIS 8 STED/FALCON system (Leica Microsystems, Mannheim, Germany). The system is equipped with a White Light Laser for excitation and a pulsed 775 nm STED laser for stimulated emission depletion. Images were acquired using an HC PL APO CS2 100x/1.40 oil immersion objective. Fluorescence signals were recorded using hybrid detectors. Samples were fixed and immunostained as described in the immunohistochemistry section. Abberior STAR 580 and Abberior STAR RED were used as flourophores coupled to secondary antibodies. A 638 nm excitation line and the corresponding notch filter were applied for selective excitation and spectral separation. Time-resolved detection (Tau-STED) was applied to further enhance lateral resolution. A temporal gate of 0.5-8 ns after excitation was set to filter out early photon counts corresponding to incomplete depletion, thereby improving spatial resolution and signal-to-noise ratio (*109*).

### Immunohistochemistry

Fixed brain slices (80–300 µm) derived from patient tissue or PDX mouse brains were transferred to PBS and stored at 4°C until further processing. For permeabilization and blocking, sections were incubated for 2 h at room temperature (RT) in blocking buffer containing 5% FBS and 1% Triton X-100 (v/v in PBS). Primary antibodies - anti-puromycin (mouse, 1:500), anti-RPS6 (rabbit, 1:100), anti-GFP (chicken, 1:1000), anti-nestin (rabbit, 1:300) and anti-drebrin (rabbit, 1:500) - were diluted in 1% FBS and 0.2% Triton X-100 (v/v in PBS), and applied overnight at 4°C. For puromycin staining in acute slices, primary incubation was extended to 72 h at 4°C to ensure sufficient antibody penetration.

Following primary incubation, slices were washed three times in 2% FBS (in PBS) for 15 min each, then incubated with appropriate secondary antibodies (ch488, ms568, rb647; all diluted 1:500 in 1% FBS, 0.2% Triton X-100 in PBS) for 4 h at RT. Sections were subsequently washed three times with 1% FBS (in PBS) for 10 min, followed by five washes in PBS for 10 min each. Nuclei were counterstained with DAPI (1:10.000 (w/v) in PBS) for 10 min. Slices were mounted using SlowFade™ Gold and the coverslip edges were sealed with nail polish.

### IHC and ICC Imaging

Confocal imaging was performed at a Zeiss LSM 900 and a Zeiss LSM 980 both equipped with an Airyscan superresolution detector (Light microscopy facility, DKFZ Heidelberg) as well as an Nikon AX Confocal Ti2 microscope equipped with a nSparc superresolution detector (Nikon Imaging Center Heidelberg). Samples were screened using a 20x objective (Zeiss LSM: NA = 0.8; Nikon AX: NA = 0.75). Imaging was done using a 40x oil immersion objective at the LSM900 (NA = 1.3, refractive index = 1.518) and a 40x silicone oil immersion objective (NA = 1.25, refractive index = 1.42) at the Nikon AX. Images were taken as a z-stack with 0.5 µm z-steps and a pixel size of 0.144 µm.

For ICC-based ribosomal cluster analysis, confocal imaging was performed under Nyquist sampling (63 nm pixel size) to approach super-resolution. Z-stacks were acquired at 100 nm intervals across selected TM regions (main shaft, TM tips, small processes), with Nikon’s Perfect Focus System used to minimize drift.

### Immunohistological signal quantification

Z stack images of fluorescently labeled cells were manually cropped to isolate regions containing one or more cells of interest. Signal probability masks were created using a pixel classification based workflow via Ilastik (*110*), with an individual training for each fluorescence channel (83) channel. For Ilastik pixel classification we chose multiscale Gaussian smoothing, edge filters (Laplacian of Gaussian, Gaussian gradient magnitude, Difference of Gaussians), and text features (structure tensor and Hessian eigenvalues) at σ = 0.3-10. Hereafter we refer to this as the standard feature set. In cases where strong signal variation along the z-axis interfered with Ilastik classification, cumulative distribution function (CDF) matching was applied beforehand.Training used the Pixel Classification workflow, which iteratively refines feature-based predictions from user annotations to better distinguish subtle cellular structures and suppress background signal. In some cases, the Autocontext workflow was additionally employed, feeding previously generated probability maps back into the classifier to further reduce small debris artifacts and produce more consistent, refined probability maps. Images were subsequently normalized in case of a signal drop-off in the z-axis by a custom Fiji script: The frame with the highest in-focus signal was selected as a reference, either from the first slice or from the slice with maximal intensity. Normalized images were then reprocessed through Ilastik with the standard feature set.

To address residual signal attenuation across z-slices, a correction function was derived from the background signal of the target channel. Intracellular signals were removed from the raw image using Ilastik probability maps (with the standard feature set) for DAPI and the respective protein channel. The resulting background was quantified slice by slice, excluding the brightest 10% of pixels to reduce artifacts. These measurements were used to plot signal intensity as a function of slice number. An exponential decay fit was applied to this plot, and correction factors derived from the resulting equation were subsequently implemented in a custom Fiji macro to normalize the raw images. Following correction, signal intensity within tumor cells was quantified by masking GFP-positive regions. The GFP probability maps generated by Ilastik were imported into Fiji and converted into binary masks, followed by 16-bit conversion. The corrected RPS6 or puromycin images were multiplied by the binary GFP masks to isolate signal originating within GFP-positive cells. Using the original files as reference, cellular compartments such as tumor microtubes (TMs) and somata were manually segmented using ROI tools in Fiji. Z-stacks were further cropped to exclude any non-target structures above or below the segmented cells. Resulting measurements of gray value and area, using a consistent threshold range, and intensity values for each ROI were extracted using multi-measure functions. The weighted average signal intensity for each ROI was calculated by the following formula:

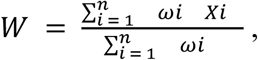

with W as weighted intensity average, Χ as signal intensity per pixel and ω as area per measured region. This resulted in a volumetric mean intensity for signal intensities in each subcellular compartment.

Corrected image stacks were duplicated and merged with the GFP probability maps to create composite images. Tracing of tumor microtubes and somata was performed using neuTube (*111*) and imported into Simple Neurite Tracer (SNT) (*112*), which was also used to quantify and compare signal intensities between different cellular compartments. Quantification was performed via the plot profile analysis function using “Disc” for the shape of each node and “Sum” for the integration metric. Tracing data were refined using fitted radii and node alignment tools, with radius parameters optimized for each structure. To improve interpretability, all traced paths were straightened, enabling direct visual comparison of signal distribution along the processes. Final visualizations and signal profiles were generated using R. Cells were classified as invasive when displaying one to two primary TMs, and as non-invasive when exhibiting more than three primary TMs. Intensity ratios between soma and TMs were compared across different cellular states and experimental conditions.

Residual signal attenuation across z-slices and subsequent correction of puromycin, RPS6 and drebrin intensities were performed as described above. Following neuTube tracing and SNT quantification via plot profile, we refined the resulting “node” values in R to ensure that each pixel was represented by a single intensity value. To classify regions of “high” and “low” drebrin, we defined any continuous stretch of ≥6 pixels (approximately 700 nm) with drebrin intensities above the mean as a high-drebrin region. For each such region, up to 16 immediately adjacent pixels with drebrin intensities below the mean were labeled as low-drebrin regions. Within these defined regions, we quantified the mean puromycin intensity to enable comparisons between high- and low-drebrin domains. Different TM segments (initial segment, main shaft and TM tip) were manually segmented.

### Analysis of tumor area

Tumor area was quantified in every third consecutive 80 µm thick brain slice as a fraction of the total slice area by Ilastik (*110*) based pixel classification training of tumor cell signal. Ilastik probability masks were subsequently imported into Fiji, binarized, thresholded and quantified as mean grey area. Relative tumor fractions were quantified by dividing the resulting tumor cell area by the overall brain slice area. Brain and tumor volumes were approximated by extrapolating the segmented areas across non-analyzed sections, areas were multiplied by the section interval (240µm) and summed.

### Calcium transient analysis and correlation

The channel of the recorded calcium imaging data was isolated and drift-corrected in AquA2 when necessary. Only GB cells displaying clear calcium transients throughout the entire recording were selected for analysis, while those showing signs of phototoxicity were excluded. Calcium transient analysis was performed using Aqua2 in MATLAB (*113*). Regions of interest were manually defined for somata, TMs and the entire cell borders. Data were globally bleach-corrected by a linear fit. The intensity threshold scaling factor was set to 2, and transients were filtered by a minimum duration of 1 s and a minimum size of 30 pixels. Temporal segmentation was enabled with a minimum seed size of 0.01, a Z-score threshold of 3.5 for seed significance, and a maximum dissimilarity of 0.8 for event merging. Spatial segmentation subsequently applied, using a pixel source size of 0.01 and a detection sensitivity of 2. Temporal extension for events was enabled, whereas global signal detection was disabled. Propagation-related metrics and network features were extracted, and decay speed was included in the analysis.

For each region, the area under the curve (AUC) of the ΔF/F₀ signal from detected events was summed and normalized to the whole-cell AUC. The same ROIs defined during calcium imaging were applied to quantify puromycin intensity in confocal image stacks acquired post fixation and immunostaining against GFP and puromycin. The average puromycin intensity per region (TMs), normalized to the average intensity of the soma, was cross-correlated with the corresponding calcium activity parameters obtained from AquA2 analysis by a linear regression fit.

### *Ex vivo* correlative light and electron microscopy

Sample preparation, confocal microscopy (Leica TCS SP8) and analysis for correlative light and scanning electron microscopy (SEM) was conducted as described before (*10*). Further, previously acquired data (*10, 64*) were integrated and analyzed. To approximate invasive potential, cells were stratified by the number of TMs visible per cell. For each cell, the number of TMs was recorded and cells were categorized as (i) unipolar (one TM), (ii) bipolar (two TMs), or (iii) multipolar (≥ 3 TMs). Uni- and bipolar cells were grouped as invasive and multipolar cells grouped as non-invasive GB cells. This categorical scheme was chosen to capture phenotypic differences linked to motility, with unipolar/bipolar morphologies typically representing more invasive phenotypes than multipolar cells (64).

### *In vivo* correlative light and electron microscopy

For conducting *in vivo* correlative light and electron microscopy (CLEM) with infrared branding, we categorized cells based on their uptake of Sulforhodamine 101 (SR101), which was monitored by using two-photon microscopy in live mice over time as described previously (*10*). Subsequently, the mice underwent transcardial perfusion by 40 ml of PBS followed by 40 ml 4% PFA. In order to facilitate the correlation process on a macroscopic level as well as on *ex vivo* confocal and electron microscopy imaging level, approximately 7 ml of Evans Blue were added into the last 20 ml of PFA at the end of the perfusion to visualize blood vessels as fiducial markers *ex vivo*. To preserve the *in vivo* orientation for imaging, the mice were decapitated, leaving the window and the titanium ring on the head, allowing immediate subsequent two-photon microscopy with consistent positioning. For the infrared branding, cells of interest that were chosen before were centered in a field of view of 694 x 694 µm using a 16x objective. A high-resolution z-stack of the area prior to branding was obtained (pixel size: 200nm, z-step-size: 520nm).

To precisely localize the region of interest within the brain *ex vivo*, infrared branding was performed (*114*). Therefore, the following settings were adjusted: The surface of the cortex was identified, the laser wavelength was changed to 800 nm and the laser power was turned to a maximum. The photomultiplier tubes (PMTs) of the microscope were turned down to accommodate the maximum laser power. Initially, a 50 µm by 694 µm segment at the lowest part of the 694 µm x 694 µm area centering the cells of interest was subjected to a two-photon laser at 800 nm wavelength focused on a depth of 50 µm beneath the surface of the cortex for a duration of 45 seconds. Subsequently, this procedure was replicated on the upper side of the target region, so that the cells of interest were then framed by two laser lesion bars that can be seen macroscopically.

This macroscopically visible region providing clear demarcation for the region of interest was cut out of the brain in the form of a cube using a surgery knife. Subsequently, the cube was embedded in agarose in a manner that its surface was parallel to the sectioning blade of the vibratome. Slices of 300 μm thickness were obtained using a Leica VT1000 vibratome.

The sample was then stained with DAPI and imaged with a high-resolution confocal microscope (Leica TCS SP8) providing an intermediate step for the correlation process. Preparation of the sample for electron microscopy was performed as previously described. Then, we captured low-resolution overviews with serial-section scanning electron microscopy with a 30-50 nm pixel size, sufficient to resolve cellular outlines. The cells of interest were unmistakably identified by their cell morphology and by the spatial arrangement of the DAPI-stained nuclei as well as the Evans blue signal as a marker for vascular structures. Lastly, we acquired the cells of interest throughout large z-volumes and reconstructed them in 3D, which enabled us to perform 3D-rendering analysis (Autodesk 3ds max).

Invasive/non-invasive cells were discerned through the colocalization of the SR101 with the intrinsic GFP signal exhibited by the cells.

### Semi-automated quantification of ribosomes

For each dataset we chose a representative section that maximized the TM area. We excluded images with contamination, poor contrast, or other artifacts. Cell boundaries were manually delineated along the plasma membrane; nucleus and outer nuclear membrane were likewise cropped manually to define reference compartments (cell, nucleus, TM, cytosol). Compartment areas were later computed from these contours using the recorded pixel size.

An autocontext workflow in Ilastik (*110*) (Pixel classification with standard feature set; standard feature families (intensity, edge, texture) across multiple Gaussian scales (σ ≈ 0.3–10)) was trained per cell (ribosomes vs. background) to retain human control and maximize accuracy. Training labels were restricted to cytoplasm; nuclei were excluded from learning.

Probability maps were further processed in Fiji with custom-made macros. The ribosome channel was interactively contrast-adjusted and thresholded, then a watershed transform was applied to split touching particles. Ribosomes were detected with the “Analyze Particles” function using size and circularity criteria adapted to acquisition scale (scaled according to pixel size). Ribosomes appear as compact, rounded particles with ∼25nm [∼250Å] in size (*115*). Minimum circularity of 0.5 was chosen to detect rounded particles. To accommodate dataset-specific variation in image resolution, pixel size and preprocessing (including training performance), the size window was set to ∼20-40nm, which reliably captured ribosome particles while excluding smaller noise features or merged larger structures. As a quality check, detections were overlaid on the raw EM image; if unsatisfactory, thresholds/parameters were adjusted and the detection repeated (human-in-the-loop). For spatial analyses, ribosome contour ROIs were converted to centroid point ROIs. Cytosolic, TM, and nuclear areas were derived from the manual contours and pixel size.

ROI files exported from Fiji (.roi/.zip) were imported into R with RImageJROI. We organized each dataset as an S4 object that stores paths to the ROI files, loaded cell/TM/nuclear polygons, ribosome centroids, and derived summaries (compartment areas in µm², cytosolic area, etc.).

Ribosomal density was computed as the number of ribosomes per cytosolic area (µm²). Per TM summaries and additional spatial readouts were computed from the centroid coordinates within the same S4 framework.

For validation the workflow was applied on a section of the dataset jrc_hela-3 (*116*) at OpenOrganelle, an open-access electron microscopy dataset (*117*) and compared with their segmentation (*118*).

For further spatial analysis, ribosomes were assigned to subcellular compartments. We manually annotated lamellipodia-like, filopodia-like and TM-like processes based on morphological criteria (Classification of GB cell TM morphology) in the EM sections. Ribosome centroids within the annotated contours were counted and normalized to compartment area to compare densities across subcellular domains. We subdivided each TM in two regions: a ∼750nm peripheral band (“Border TM”) adjacent to the plasma membrane, and the remaining interior (“Inner TM”). TM segments that bent by more than 15° were labeled as convex. The convex region of interest was defined as a membrane-adjacent band traced along the curvature peak, with its width adapting locally to the membrane contour until the TM straightens again. Ribosome density within this curvature-guided convex band was then compared to the remaining TM area.

### Quantification of ER-bound and free ribosomes

For each ribosome centroid, we measured the minimal euclidean distance to the nearest ER contour; those ≤ 20 nm were annotated as “ER_bound,” whereas all others were considered “free.” Cells with missing or poorly segmented ER contours, or with strong image artifacts in the ER region, were excluded from this analysis.

### Quantification of monosomes and polysomes

To distinguish monosomes from polysomes, pairwise distances between ribosome centroids were computed and spatial clustering was performed. Ribosome centroids separated by ≤40nm were grouped into clusters; clusters containing only a single ribosome were classified as monosomes, while those with ≥2 ribosomes were defined as polysomes.

### Ribosomal cluster density quantification (ICC)

Cellular and ribosomal signals were segmented using 3D machine learning in Ilast(*81*) (standard feature set), generating ch(83)l-specific probability maps. Ribosomal clusters were extracted in 3D with a custom Cellpose model, and ROIs defined in Fiji to restrict analysis. TM-specific ribosomal signals were isolated by overlapping GFP probability maps with Cellpose cluster segmentations. Cluster density (per µm^3^) was quantified using the 3D ImageJ Suite plugin.

### Classification of GB cell TM morphology

Primary TMs were defined as processes originating directly from the GB cell soma, with a length greater than 10 µm; processes emerging from other processes were not considered primary. In live cell experiments, the leading TM was defined as the primary TM which drives the overall cell motility. In images without a live cell correlation, the leading TM was defined as the more prominent or elongated TM, typically exhibiting greater length, thickness, and sometimes a noticeable thickening at the TM tip compared with the opposing TM. In EM images filopodial structures were defined as structures shorter than 10 μm and thinner than 1.25 μm. Structures that were shorter than normal TMs (10 μm), but exhibiting TM diameters from 1.25 μm to 3 μm were classified as TM-like structures. Short structures that were thicker than normal TM (> 3 μm) were classified as lamellipodia-like structures. Sprout points were defined as cell membrane sites where a protrusion emerges from the TM, operationally identified as the transition zone between the flat plasma membrane and the emerging protrusive structure. For quantification the region of interest includes the protrusion itself and an adjacent segment approximately 300nm to both sides and into the cell interior.

### MERFISH image-based transcript quantifications

MERFISH imaging data were processed with nucleus and cytoplasm segmentation masks. Cell centers and nuclear boundaries were identified using DAPI-based segmentation. For each cell, the euclidean distance of every transcript to the cell centroid was calculated. Transcripts located within a defined radius were assigned to the corresponding cell. To ensure accurate distance estimation, only isolated cells were considered. Isolation was defined by nearest-neighbor distances between cell centroids exceeding a 25 µm threshold. For each transcript, the distance to the cell centroid was calculated and compared to the mean nuclear radius derived from DAPI segmentation. Genes with average distances smaller than the mean nuclear radius were classified as nuclear-enriched, whereas genes with distances larger than this threshold were classified as cytosol-enriched. Per-gene mean distances and standard errors were computed across all cells, and genes were ranked accordingly.

For all subsequent analyses, mono- or bipolar cells were selected based on the polyT channel. GB subcompartments were segmented and morphologically classified within the MERSCOPE Visualizer software as soma, leading TM, or trailing TM. In cases where morphological polarity was ambiguous, the cell was excluded. To compare transcript abundance between leading and trailing TMs, expression matrices were exported from the Visualizer, and the total number of transcripts was summed for each group. For analyses of cytoskeletal transcripts, the complete set of genes was screened for genes part of the GO “terms myelin sheath” (GO:0043209), “microtubule binding” (GO:0008017), “kinesin complex” (GO:0005871), “actin cytoskeleton organization” (GO:0030036), and “neuron projection development” (GO:0031175). Only these cytoskeletal transcripts were included in subsequent analyses (structural cytoskeletal genes in table S3).

For the analysis of transcript distribution along the TM, images of selected cells were exported twice: once with the polyT channel only and once with the transcript channel only. PolyT images were imported into neuTube for tracing of the leading TM, with 200 tracing points placed between the TM initial segment and the TM tip, thereby subdividing the TM into nodes. The corresponding transcript-channel image was imported into the SNT plugin of Fiji, using the neuTube tracing points as positional references. Signal intensity was measured for each of the 200 nodes using the “Plot Profile” function selecting “Disc” for the shape of each node and “Sum” for the integration metric and saved in tabular format. To convert intensity to transcript number, the signal of a single transcript was measured, and the total intensity of each segment was divided by this value. TM length was then normalized to a 0-100% scale. For comparisons between TM regions, the initial segment was defined as 0-20% of TM length, the shaft as 20-80%, and the TM tip as 80-100%.

To further characterize the TM transcriptome a principal component analysis was performed using Seurat and a k-nearest-neighbor graph was constructed to perform Louvain clustering. A UMAP was created for visualization.

Cell invasiveness was assigned based on morphology by counting primary TMs. Cells with two or fewer primary TMs were classified as invasive and cells with three or more primary TMs were classified as non-invasive. Based on the classification of the respective cells each TM was assigned an invasiveness label. When calculating the proportions of each invasiveness label in the TM cluster a normalization was applied to account for cells with many TMs possibly dominating an unnormalized analysis: For each TM a weight equal to 1/(number of TMs for that cell) was used, then for each cluster the proportion of invasive vs. non-invasive contribution was computed as the sum of these weights within each invasiveness group divided by the total weighted count in that cluster. By that each cell contributes equally regardless of TM number.

To investigate the correlation of our TM clustering to the Neftel et al. cell state classification (*2*), the top 30 genes ranked by absolute PC1 loading from the TM-based principal component analysis were defined as a module. In our PDX scRNA dataset the Neftel et al. cell state classification (*2*) was applied to the cells and a PC1 module score was computed per cell as the mean expression of the module genes. The mean and SD of the PC1 module score was calculated per cell state and global (Kruskal–Wallis) and pairwise (Wilcoxon test with BH adjustment) significance tests were performed.

### RNA bulk sequencing analysis

The raw reads from the sequencing were at first quality-controlled using FastQC. Transcript-level quantification was performed against the human reference transcriptome (Ensembl GRCh38, version 86) using kallisto, an RNA-quantification program / abundance estimation tool producing estimated counts and abundance metrics. These estimates were imported and summarized to the gene level using tximport (v. 1.34.0) in R. A DESeqDataSet was constructed and modeled regarding the effects of passage and compartment. Genes with a very low expression were filtered out prior to model fitting by retaining only those with a total count across all samples of at least 10. Normalization and differential expression analyses were performed with DESeq2 (v.1.46.0). Genes with adjusted p-value (padj) below 0.05 (and an effect size threshold such as |log2FC| > 0.5) were considered significantly differentially expressed between TM and soma compartments. We projected the published cell-state marker gene sets from Neftel et al. (*2*) onto bulk-RNA-seq differential gene expression results. For each of the eight meta-modules (MES1, MES2, AC, OPC, NPC1, NPC2, G1/S, G2/M), Neftel et al. (*2*) defined all genes whose average single-cell log-ratios exceeded 2, then took the top 50 highest-ranking genes. We merged MES1 and MES2 into a single “MES” set, and NPC1 and NPC2 into “NPC” and removed modules “G1/S” and “G2/M”. For each of the four remaining modules (MES, AC, OPC, NPC) we generated a volcano plot of log2FC (TM vs. Soma) against –log₁₀(p-value) using ggplot2. Differentially expressed gene heatmaps were generated by the package “pheatmap”. Enriched GO-terms for differentially expressed genes were identified by the “Enrich GO”. To retrieve cytoskeleton associated genes the Homo sapiens annotation database (org.Hs.eg.db) was queried for the Gene Ontology (GO) terms the GO:0005856 (“cytoskeleton”) or a combination of GO:0007017 (“microtubule-based process”), GO:0005882” (“intermediate filament”), GO:0005884 (“actin filament”) and GO:0034453 (“microtubule binding”) respectively.

### Comparison of single-cell and single-nuclei sequencing data

For the single-cell RNA-seq data we used the GBMap dataset and for single-cell nucleus data the GBspace data set was used. Raw count matrices were processed using Scanpy (version 1.11.4). Cells with low library complexity and high mitochondrial transcript fractions were excluded. Remaining counts were normalized to a fixed library size per cell and log-transformed. When raw counts were preserved, normalized values were stored in a separate layer to allow consistent downstream scoring. Cell state signatures were quantified according to the procedure originally described by Neftel et al (*2*). For a given gene set each cell was assigned a signature score that reflects the relative abundance of the gene set compared to matched control genes. Relative expression was defined as the deviation of a gene’s expression in a given cell from the average expression of that gene across all cells. Control gene sets were constructed by binning all genes into thirty equally sized groups according to mean expression levels. For each signature gene, one hundred genes were randomly sampled from the same bin, resulting in a control set that was one hundred-fold larger than the original gene set and matched for overall expression distribution. The final signature score was defined as the average relative expression of the signature genes minus the average relative expression of the control set. For the definition of GB cell states, the four major transcriptional programs were considered: mesenchymal, astrocytic, oligodendrocytic progenitor-like, and neural progenitor-like. For each cell, scores for all four programs were calculated as described above. Cell state were assigned according to Neftel et al. as described before (*2*). First, cells were separated into progenitor-like (oligodendrocytic progenitor-like or neural progenitor-like) versus differentiated (astrocytic or mesenchymal) according to which branch displayed the higher maximum score. Within each branch, the final state was determined by comparing the two corresponding program scores, assigning cells to the program with the stronger signal. This approach ensures that each cell was uniquely assigned to one of the four canonical states. For each program, we derived a subset of genes enriched in TMs. TM signatures were scored using cell state signatures established by Neftel et al. (*2*), ensuring comparability with full program scores. To assess the capacity of TM-restricted signatures to recapitulate state definitions, we compared TM-derived predictors against full-signature–based labels in both single-cell and single-nucleus datasets. Ground-truth labels were defined from full program scores using the Neftel el al. described cell state scoring (*2*). TM predictors were constructed as pairwise contrasts within each program branch, for example mesenchymal versus astrocytic or oligodendrocytic progenitor-like versus neural progenitor-like. Performance was evaluated by receiver operating characteristic analysis with full-signature assignments as the reference with the area under the curve as a quantitative output parameter.

### Mapping of transcriptomic TM and soma signatures onto single-cell RNA sequencing datasets and reclustering

Single-cell RNA-sequencing data from S24 PDX (*10*) was used, as well as the “core GBmap” (*48*), which integrated 110 patients and over 330,000 cells from 16 studies. Subsequent data analysis was conducted using Scanpy (version 1.11.4). Among the malignant cells, four cellular states, AC-like, NPC-like, OPC-like, and MES-like, were identified as described by Neftel et al. (*2*). Four gene sets were tested, namely top significantly differentially expressed genes for TM and soma ranked by fold changes, random genes, and highly variable genes as calculated by Scanpy under default parameters. An identical number of genes was selected from the top of each gene list, up to 300 genes. In each sample with over 100 malignant cells, the normalized expression matrix of tumor cells was subset to genes from the aforementioned gene sets, and a principal component analysis (PCA) was performed with 10 principal components (PCs). We then evaluated the performance of the gene sets in separating the cell state clusters via the silhouette score metric on the PC matrix, using the module scikit-learn (version 1.7.1), obtaining one score for each gene set per sample. Additionally, we performed PCA on over 10,000 cells randomly sampled across the integrated dataset, retained cells expressing > 5 genes, and visualized the cell state clusters using Uniform Manifold Approximation and Projection (UMAP) on the top 10 PCs. To represent cell state transitions via riverplots, data matrices only including TM signature genes were reassigned one of the four main Neftel et al. cell states (*2*) as described above.

### Transcript mapping and quantification in patient-derived S24 cells (MERFISH platform)

S24 mGFP mono-cultures were seeded onto poly-L-lysine-coated MERSCOPE non-beaded imaging slides. After five days of culture, cells were fixed with 4% PFA for 10 minutes at room temperature. Following storage in PBS, samples were processed according to the manufacturer’s standard protocol, including permeabilization (*119*) followed by the workflow described in the User Guide (*120*). Samples were imaged sequentially using a custom-designed panel comprising 500 genes selected for relevance to glioblastoma cell biology (custom gene panel; table S3). Acquired image overlays, including transcript signals as well as DAPI and polyT channels, were manually segmented to obtain gene expression matrices corresponding to somata and TMs. Gene counts for each matched soma and TM were subsequently normalized to generate a weighted pseudo-bulk expression matrix per cell and further processed for GO-term enrichment as described above.

### Transcript mapping and quantification in human FFPE sections (Xenium platform)

Glioblastoma patient FFPE sections of the tumor infiltration zone with a thickness of 5µm were processed according to the suppliers instructions (10X genomics) for the commercially available “Xenium Human Brain” panel. After sequencing, sections were deparaffinized by incubating twice in 100% xylene, followed by two washes in 99% ethanol, and then sequentially in 95%, 70%, and 50% ethanol. Finally, the sections were rinsed in MilliQ water, with each step lasting 3 minutes. Antigen retrieval was subsequently performed by immersing the samples in a 20 mM sodium citrate buffer (pH 8.0, prepared in MilliQ water) contained in a glass cuvette. For antibody epitope retrieval, the samples were subsequently heated in a microwave at 360 W for three to four cycles to boiling, allowing brief cooling between each cycle, and then incubated in an oven at 60 °C for 30 minutes and stained for nestin (rb, 568) and DAPI as described above. For image registration, subsequently acquired widefield and confocal microscopy images were manually aligned to the Xenium DAPI reference within Xenium Explorer: Images were imported and manually aligned to the Xenium DAPI signal by iteratively adjusting rotation and mirroring and by placing corresponding anatomical landmarks to achieve precise registration of all microscopy channels to the Xenium coordinate space. This procedure ensured accurate spatial correspondence between high-resolution cellular morphology and the underlying transcript distribution.

### Statistical analyses

Displayed qualitative and quantitative data plots were generated via the ggplot2 package in R or Graphpad Prism 10.6.1. Statistical analyses were performed in Graphpad Prism 10.6.1. Before choosing appropriate tests, all data was checked for gaussian normality distribution via Shapiro-Wilk test. T-test was used to compare normally distributed dataset with two groups, while Mann-Whitney Rank test was chosen for non-normally distributed data. For paired data (puromycin/RPS6 quantification within the same cell, EM based ribosomal quantifications within the same cells or MERFISH/Xenium based quantifications within the same cell), data was compared accordingly with paired t-tests for normally distributed and Wilcoxon Rank test for non-normally distributed data. For datasets with more than two groups, One-way Anova with follow-up tests of Tukey’s multiple comparisons test (normally distributed data) or Dunn’s multiple comparisons test (non-normally distributed data) was used. Data was considered significantly different for p-values < 0.05 or adjusted p-values < 0.05 for differential gene expression analyses taking false discovery rates into account.

---

**Fig. S1.**
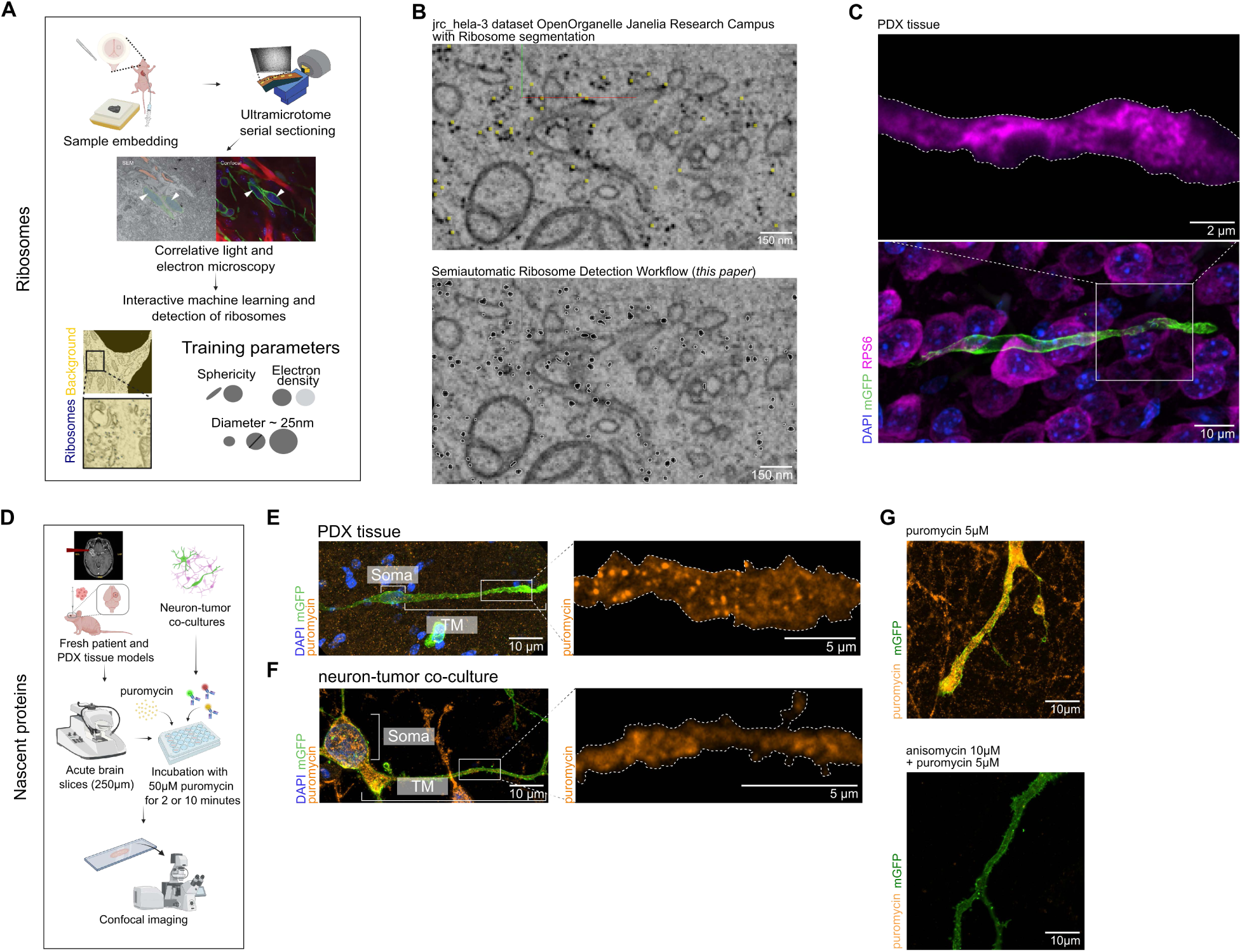
Technical validation and multimodal application of ribosome segmentation and nascent protein visualization via puromycylation assays and AHA labeling. (**A**) Electron microscopy and image segmentation workflow depicting the identification of ribosomes in GB cells in PDX tissue by a supervised machine learning algorithm (for details Materials and Methods). (**B**) Performance comparison of ribosome segmentation on an image out of the jrc_hela-3 dataset from OpenOrganelle of the Janelia Research C(*_87_*). Top: Reference Segmentation by OpenO(89)elle. Bottom: Ribosome segmentation generated using our proposed workflow for the same dataset. (**C**) Representative confocal immunohistochemistry (IHC) image (standard deviation based projection) of a S24 mGFP GB cell in PDX tissue, stained for DAPI (blue), GFP (green) and the ribosomal protein S6 (RPS6, magenta). Zoom-in displays RPS6 signal in the TM with a mGFP based crop-out of the cell outline (dashed line). (**D**) Workflow for visualizing nascent proteins in neuron-tumor co-cultures, GB patient and PDX samples using a puromycylation assay. (**E**) Representative confocal IHC image of a patient-derived mGFP GB cell (S24) stained for DAPI (blue), GFP (green) and puromycin (orange) in PDX tissue. Zoom-in displays puromycin signal in the TM with a mGFP based crop-out of the cell outline (dashed line). (**F**) Representative confocal immunocytochemistry (ICC) image of a S24 mGFP GB cell stained for DAPI (blue), GFP (green) and puromycin (orange) in a neuron-tumor co-culture. Zoom-in displays puromycin signal in the TM with a mGFP based crop-out of the cell outline (dashed line). (**G**) Validation of puromycin signal by preincubation of S24 mGFP GB cells in neuron-tumor cocultures with anisomycin to inhibit puromycin binding. Stained for GFP (green) and puromycin (orange).

**Fig. S2.**
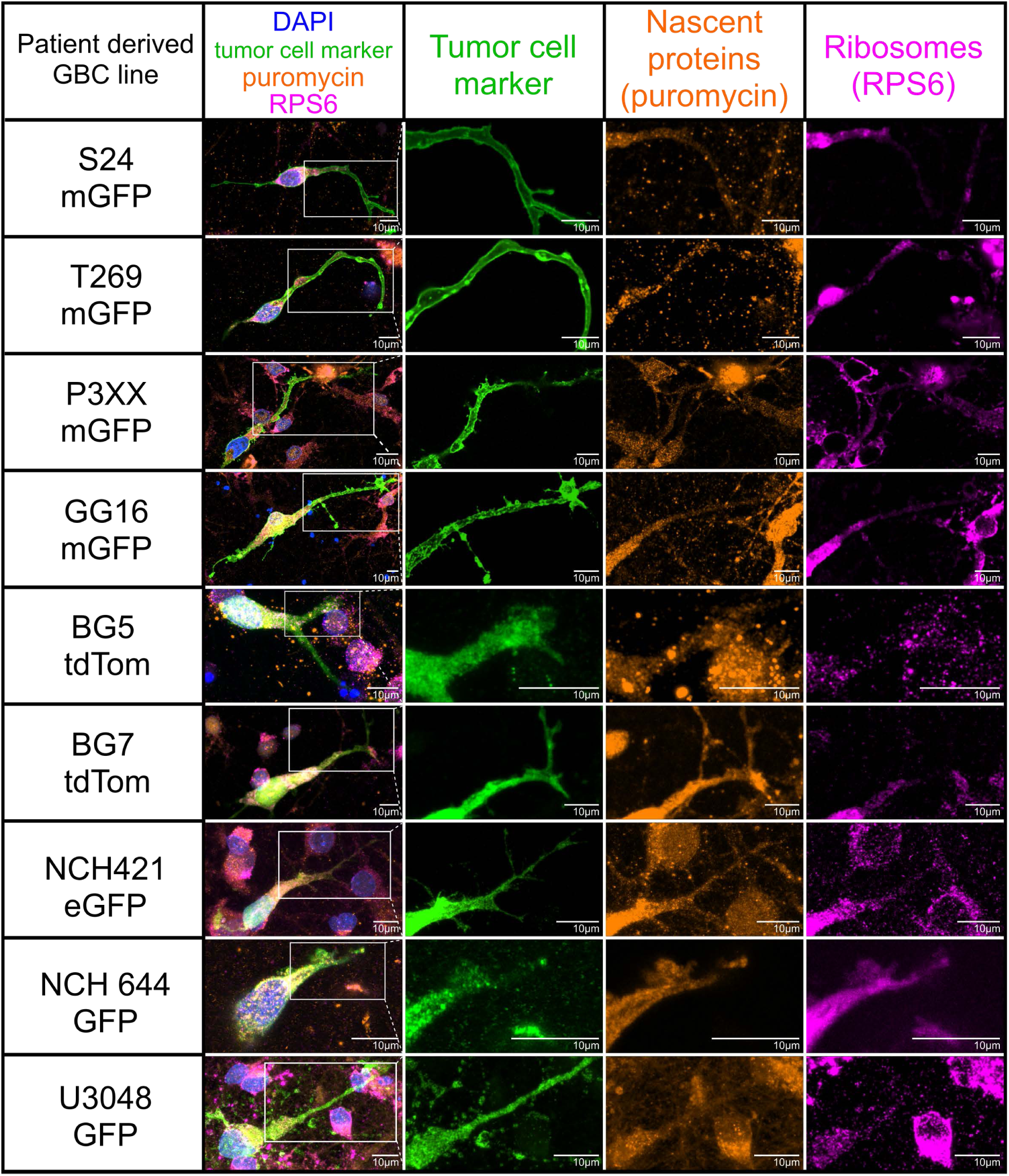
Ribosomal and nascent protein abundance in various patient derived GB models in neuron-tumor co-cultures. Representative ICC images of patient-derived GB models in neuron-tumor co-cultures, stained for DAPI (blue), mGFP, tdTomato, eGFP or GFP (green), puromycin (orange), and ribosomal protein S6 (magenta). Zoom-ins display cell markers, puromycin and RPS6 signal in the TM.

**Fig. S3.**
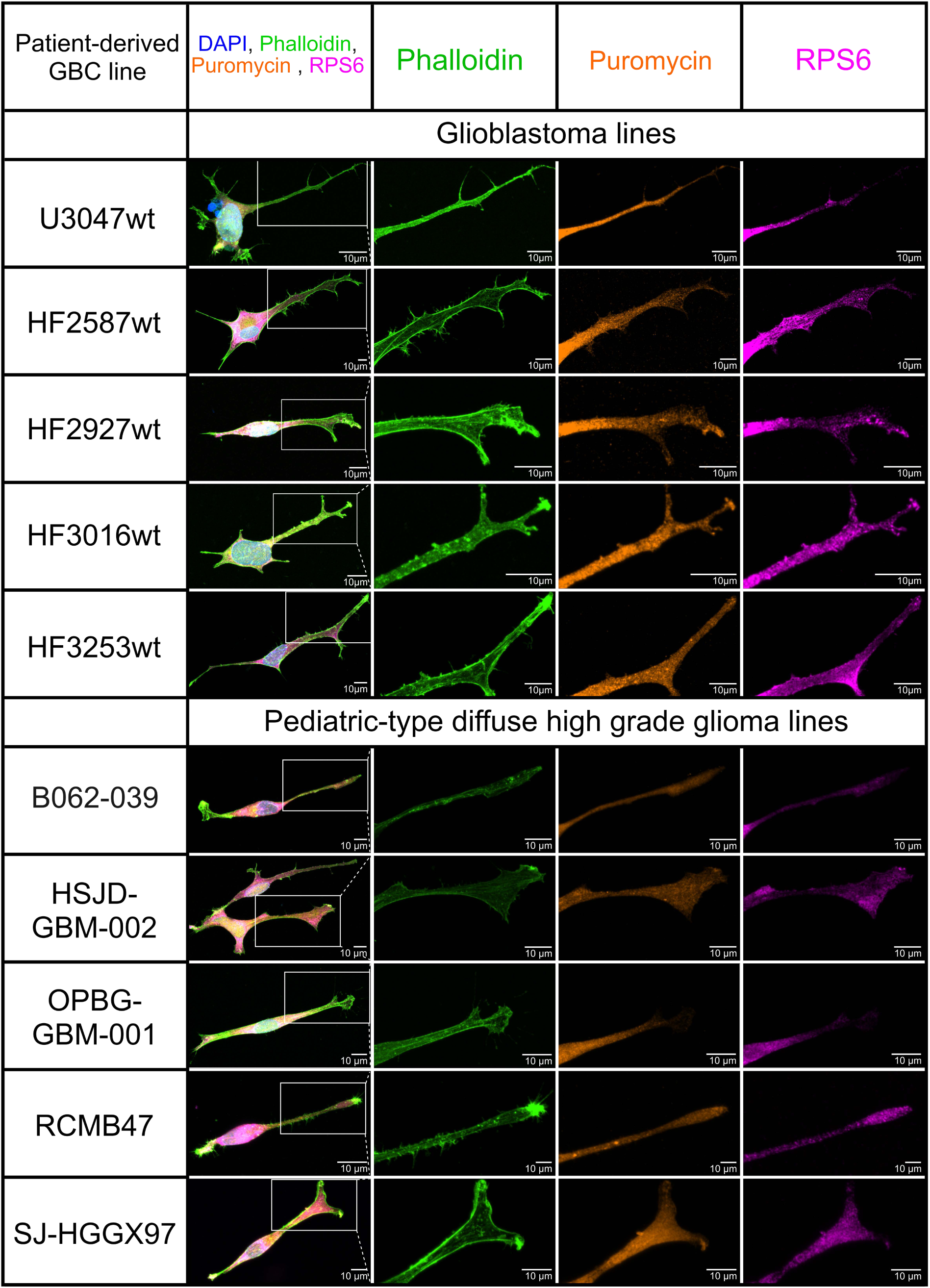
Ribosomal and nascent protein abundance in various patient derived GB models and pediatric-type diffuse high grade glioma models in mono-cultures. Representative ICC images of patient-derived GB models in mono-culture stained for DAPI (blue), phalloidin (green), puromycin (orange), and ribosomal protein S6 (magenta). Zoom-ins display cell marker, puromycin or RPS6 signal in the TM.

**Fig. S4.**
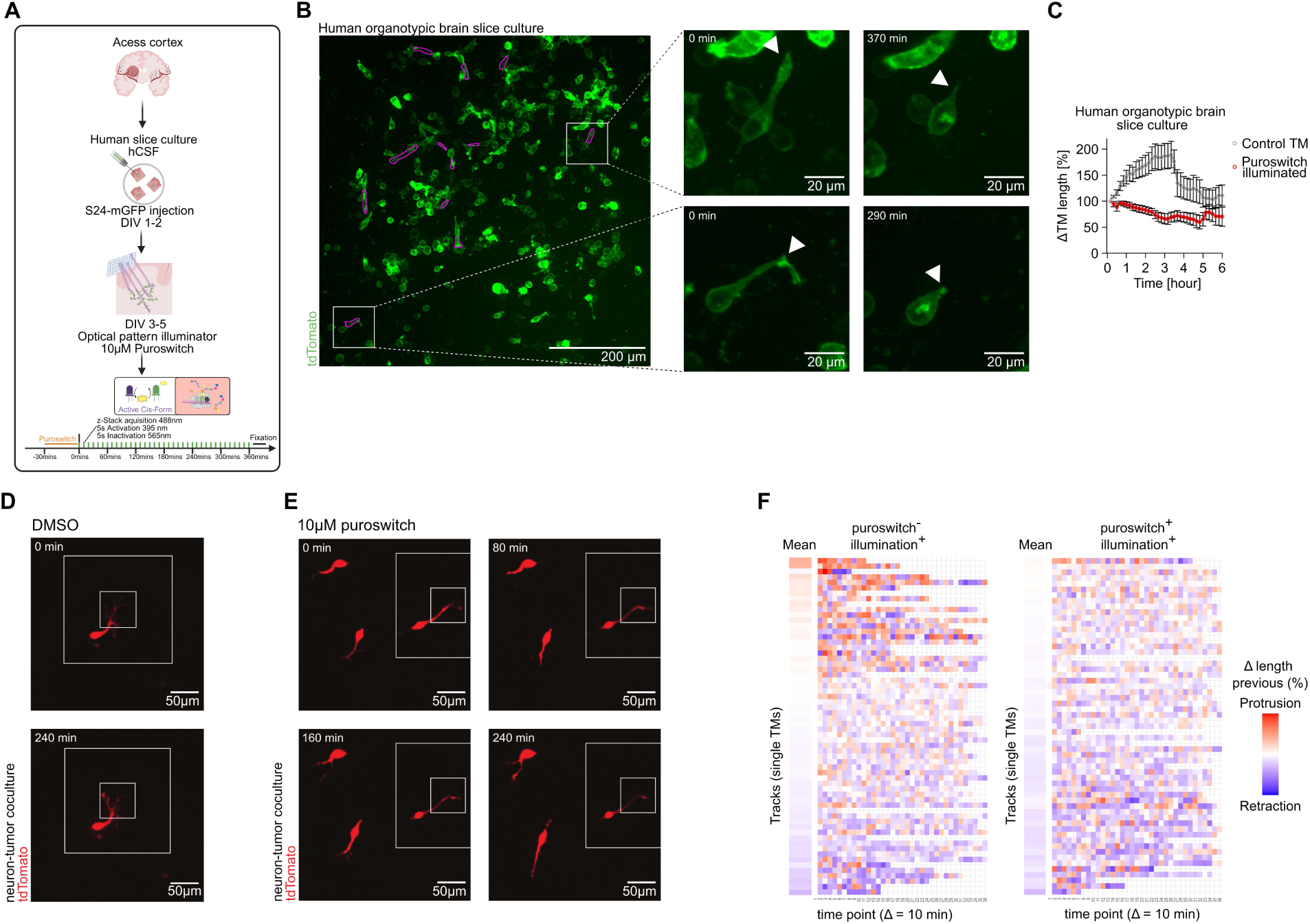
Functional validation of spatial puroswitch-mediated translation inhibition in glioblastoma models. (**A**) Workflow for spatial inhibition of local translation using puroswitch within singular TMs and inactivation simultaneously of associated cell somata in both human organotypic brain slice cultures. (**B**) Left: Representative image of a S24 grafted human organotypic brain slice culture treated with puroswitch, purple outlines indicating illuminated TMs. Right: Two representative S24 GB cells treated with puroswitch before and after illumination leading to TM retraction. (**C**) Quantification of the relative mean cumulative change of TM length over time in puroswitch-treated, illuminated and of non-puroswitch-treated non-illuminated S24 GB cells in human organotypic brain slices (n = 25 control non illuminated TMs, n = 12 puroswitch treated illuminated TMs). (**D**) Representative motility of S24 GB cells in neuron-tumor co-cultures in DMSO treated samples during photostimulation described in Fig. 1Q. The inner, smaller box represents the 405nm activation zone, the outer box represents the 555nm inactivation zone. (**E**) Representative motility of S24 GB cells in neuron-tumor co-cultures in puroswitch treated samples during photostimulation described in Fig. 6B. The inner, smaller box represents the 405nm activation zone, the outer box represents the 555nm inactivation zone. (**F**) Heatmap of TM motility over the course of live cell imaging from (D) and (E) in photostimulated DMSO control TMs and photostimulated puroswitch treated TMs. The color code represents the TM motility (red = protrusion, blue = retraction), rows represent TMs of single cells, columns represent the consecutive 10-minute time bins.

**Fig. S5.**
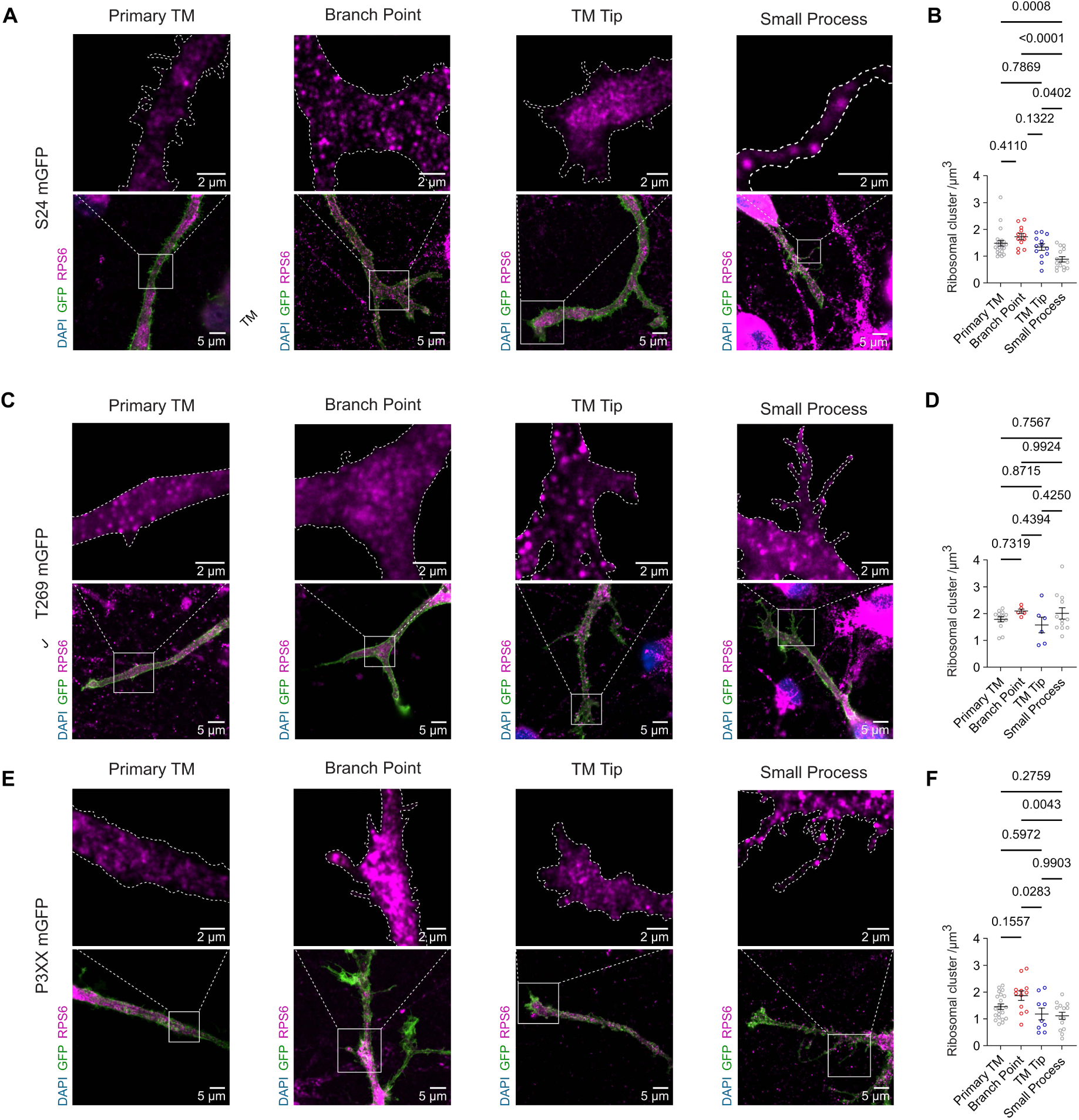
Immunocytochemical quantification of ribosomal clusters at tumor microtube hotspots, related to. **Figure 5**. (**A**) Representative ICC image of a S24 GB cell in neuron-tumor co-culture, stained for GFP (green) and RPS6 (magenta) in a primary TM, TM branch point, TM tip and small process. Zoom-ins display ribosomal clusters with a mGFP based crop-out of the cell outline (dashed line). (**B**) Quantification of RPS6 cluster density in primary TMs (n = 22), branch points (n = 12), TM tips (n = 13) and small processes (n = 15) of S24 GB cells in neuron-tumor co-cultures. (**C**) Representative ICC image of a T269 GB cell in neuron-tumor co-culture, stained for GFP (green) and RPS6 (magenta) in a primary TM, branch point, TM tip and small process. Zoom-ins display ribosomal clusters with a mGFP based crop-out of the cell outline (dashed line). (**D**) Quantification of RPS6 cluster density in primary TMs (n = 13), branch points (n = 5), TM tips (n = 6) and small processes (n = 12) of T269 GB cells in neuron-tumor co-cultures. (**E**) Representative ICC image of a P3XX GB cell in neuron-tumor co-culture, stained for GFP (green) and RPS6 (magenta) in a primary TM, branch point, TM tip and small process. Zoom-ins display ribosomal clusters with a mGFP based crop-out of the cell outline (dashed line). (**F**) Quantification of RPS6 cluster density in primary TMs (n = 21), branch points (n = 12), TM tips (n = 9) and small processes (n = 14) of P3XX tumor cells in neuron-tumor co-cultures.

**Fig. S6.**
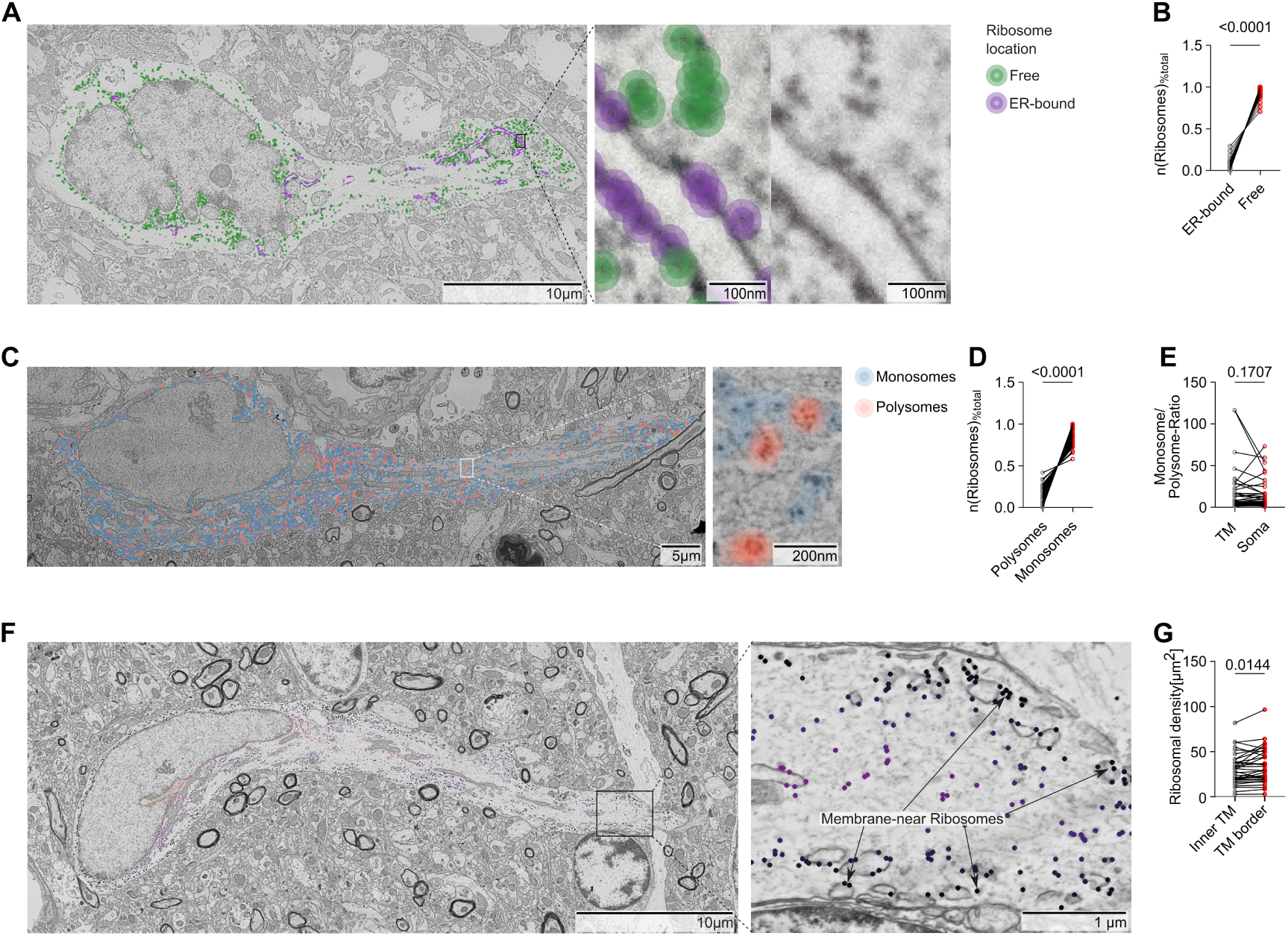
Subcellular distribution of ribosomes in tumor microtubes of patient-derived GB cells. (**A**) Representative electron microscopic image of a GB cell in PDX tissue showing the distribution of free (green) and endoplasmic reticulum (ER)-bound (purple) ribosomes. (**B**) Quantification of the proportion of free and ER-bound ribosomes among all ribosomes of GB cells in PDX tissue (n = 23 cells). (**C**) Representative EM image of a GB cell in PDX tissue showing the distribution of monosomes (blue) and polysomes (red). (**D**) Quantification of monosomes and polysomes in GB cells in PDX tissue via EM as a percentage of total ribosomes. (**E**) Quantification of the monosome-to-polysome ratio in TMs compared with somata of S24 GB cells in PDX tissue. (**F**) Representative EM image of a patient-derived GB cell highlighting membrane-near ribosomes (black) by an overlay. (**G**) Quantification of ribosomal density in the inner TM compared to the TM border (< 750 nm distance to cell membrane).

**Fig. S7.**
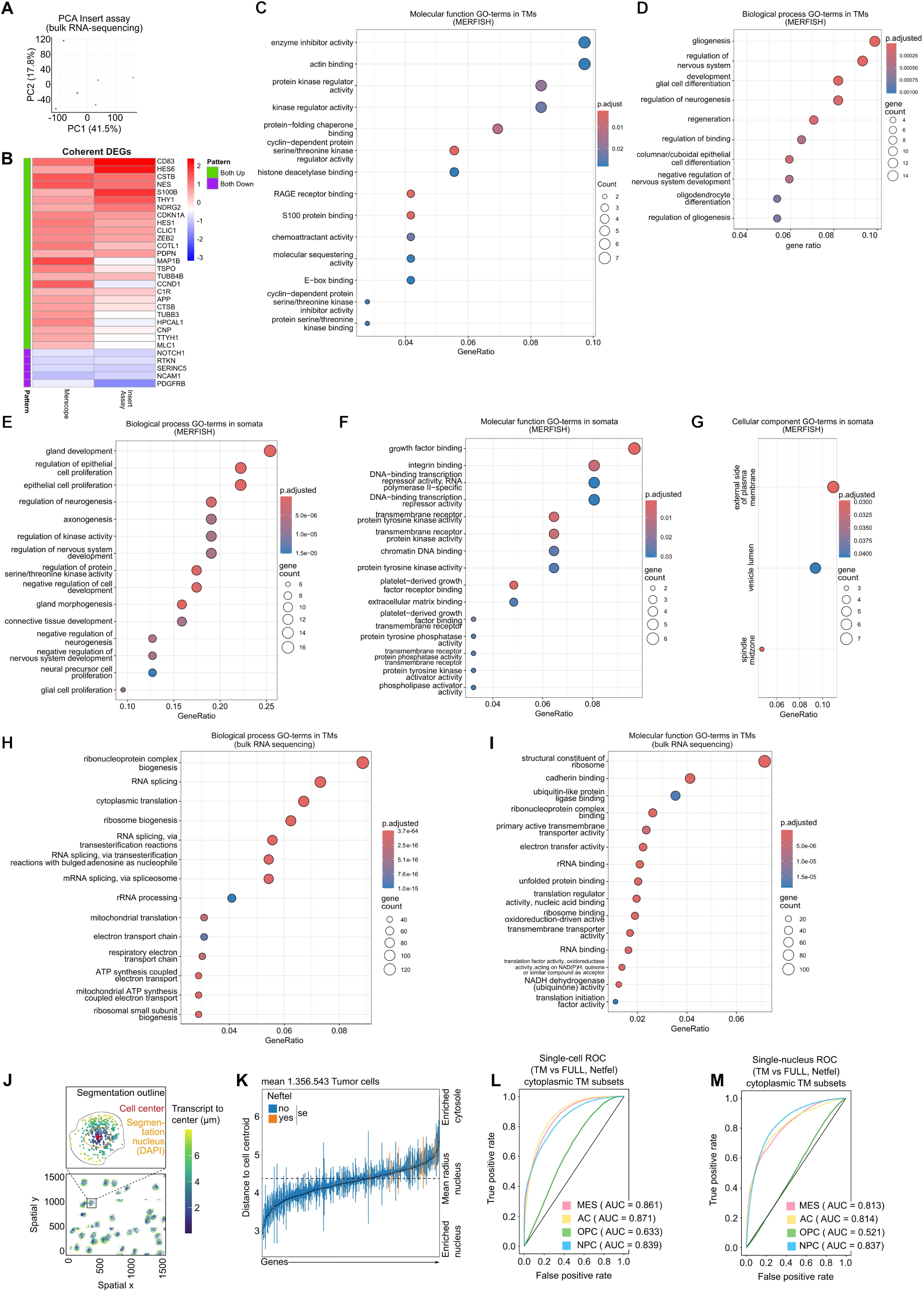
Transcriptomic profiling in subcellular compartments of GB cells. (A) Two dimensional PCA plot of gene expression profiles of TM (red) and soma (gray) enriched samples (insert assay) generated by bulk sequencing of S24 GB cells. (B) TM-upregulated differentially expressed genes (DEGs) shared between Merscope and Insert Assay datasets. (D) Dotplot of identified “Biological process” GO terms associated with RNA transcripts enriched in TMs of S24 GB cell mono-cultures in the MERFISH dataset. (E) Dotplot of identified “Biological process” GO terms associated with RNA transcripts enriched in somata of S24 GB cell mono-cultures in the MERFISH dataset. (F) Dotplot of identified “Molecular function” GO terms associated with RNA transcripts enriched in somata of S24 GB cell mono-cultures in the MERFISH dataset. (G) Dotplot of identified “Cellular component” GO terms associated with RNA transcripts enriched in somata of S24 GB cell mono-cultures in the MERFISH dataset. (H) Dotplot of identified “Biological process” GO terms associated with RNA transcripts enriched in TMs of S24 GB cell mono-cultures in the insert assay-based bulk RNA sequencing dataset. (I) Dotplot of identified “Molecular function” GO terms associated with RNA transcripts enriched in TMs of S24 GB cell mono-cultures in the insert assay-based bulk RNA sequencing dataset.(J) Segmentation of glioblastoma cells via DAPI signal into a nuclear and cytosolic compartment and color-coding of MERFISH transcripts to the cell centroid. (K) Distribution of transcripts enriched in the nuclear and cytoplasmic area for MERFISH data. Neftel et al. (2) cell state signature genes being part of the gene panel are displayed in orange. (L) Benchmarking of TM-derived signatures against the full Neftel et al. (2) programs in single-cell datasets. (M) Benchmarking of TM-derived signatures against the full Neftel et al. (2) programs in single-nuclei datasets. Statistical comparison of AUCs between single-cell RNA and single-nuclei RNA showed significant differences between MES (p = 0.021), AC (p = 0.044) and OPC (p = 0.0024) clusters.

**Fig. S8.**
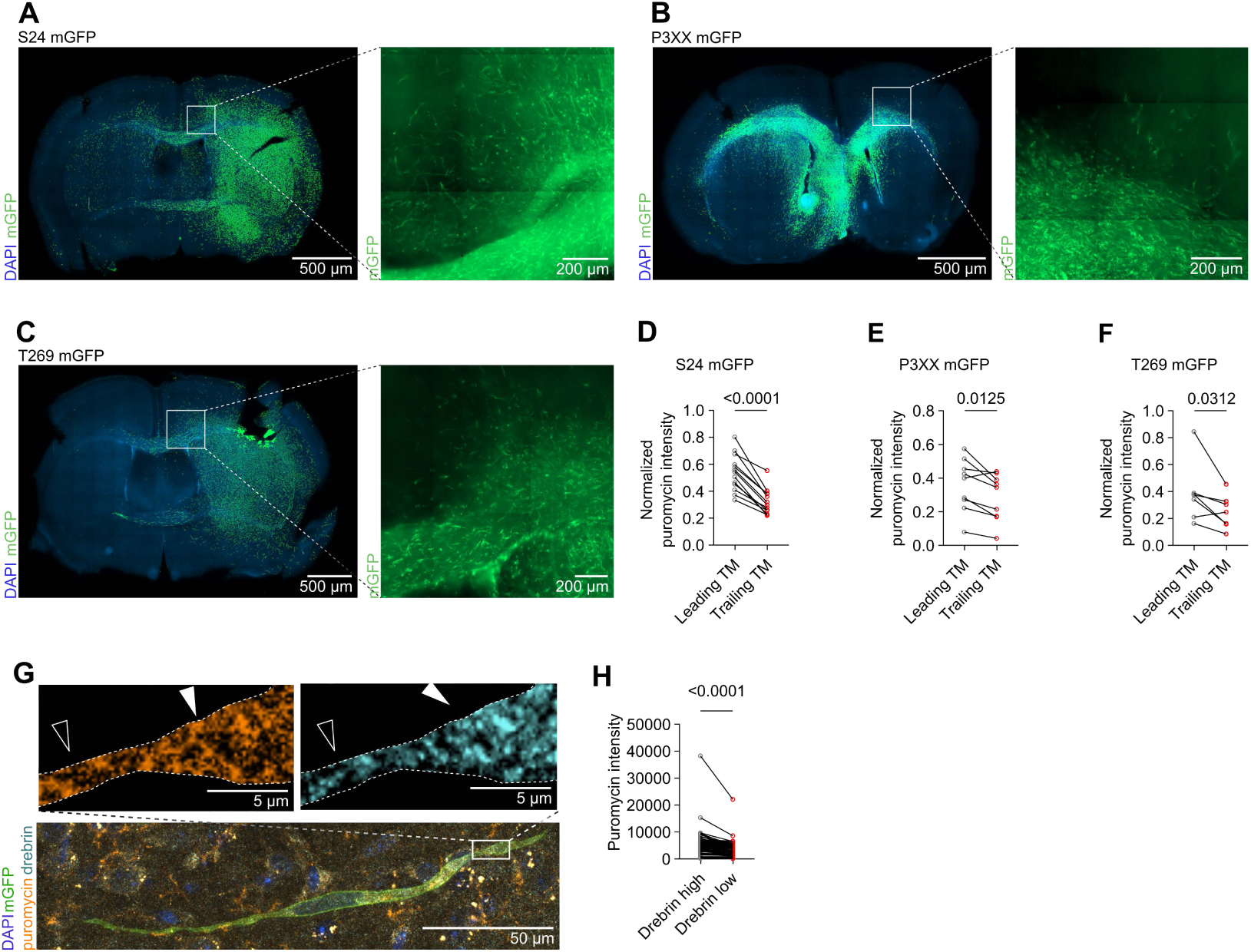
Puromycin intensities correlate with TM protrusion behaviour across patient derived GB cell lines. (A) Representative IHC image of a S24 PDX brain slice, stained for DAPI (blue) and mGFP (green, displayed as a mask). Zoom-in displays native mGFP signal with the characteristic invasion zone at the tumor-brain interface. (B) Representative IHC image of a P3XX PDX brain slice stained against DAPI (blue) and mGFP (green, displayed as a mask). Zoom-in displays native mGFP signal with the characteristic invasion zone at the tumor-brain interface. (C) Representative IHC image of a T269 PDX brain slice stained against DAPI (blue) and mGFP (green). Zoom-in displays the characteristic invasion zone at the tumor-brain interface. (D) Quantification of normalized puromycin intensities in leading and trailing TMs of S24 GB cells in neuron-tumor co-cultures (n = 13 cells). (E) Quantification of normalized puromycin intensities in leading and trailing TMs of T269 GB cells in neuron-tumor co-cultures (n = 7 cells). (F) Quantification of normalized puromycin intensities in leading and trailing TMs of P3XX GB cells in neuron-tumor co-cultures (n = 9 cells). (G) Representative IHC image (standard deviation based projection) of a S24 mGFP GB cell in PDX tissue stained for puromycin (orange), drebrin (turquoise) and GFP (green). Zoom-ins display puromycin or drebrin signal in a high drebrin (filled arrow head) and low drebrin (outlined arrow head) region with a mGFP based crop-out of the cell outline (dashed line). (H) Quantification of soma-normalized puromycin intensity in high drebrin and neighboring low drebrin TM segments of GB cells in PDX tissue (n = 94 segments from 12 GB cells).

**Fig. S9.**
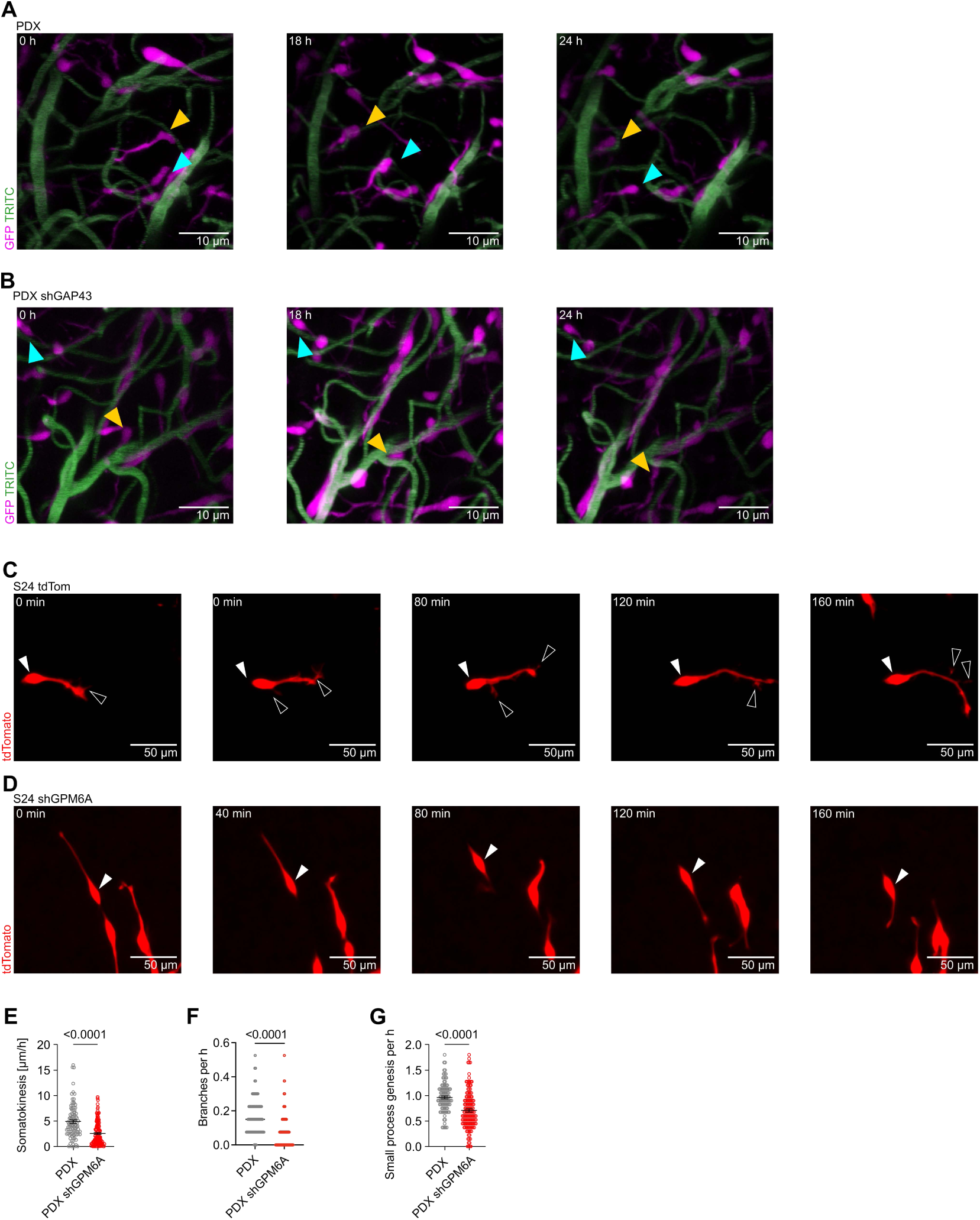
GAP43 intravital imaging and GPM6A knockdown in vitro. (A, B) Representative somatic movement (indicated by arrow heads) of S24 GFP (magenta) and S24 shGAP43 GFP (magenta) GB cells in PDX tissue over a time frame of 24 hours during intravital two-photon imaging (standard deviation-based projection) combined with an angiogram (green). (C, D) Representative motility of S24tdTomato and S24 shGPM6A tdTomato GB cells over the course for live-cell imaging in neuron-tumor co-cultures. White arrow-heads represent the cell soma while black arrow-heads represent the formation of new small processes. (E) Quantification of somatokinesis in S24 tdTomato GB cells compared to S24 shGPM6A tdTomato GB cells during live cell imaging in neuron-tumor co-cultures (n = 96 S24 GB cells, n = 136 S24 shGPM6A GB cells). (F) Quantification of TM-branching events per hour in S24 tdTomato compared to S24 shGPM6A tdTomato GB cells during live cell imaging in neuron-tumor cocultures (n = 96 S24 cells, n = 136 S24 shGPM6AGB cells). (G) Quantification of small process generation in S24 tdTomato compared to S24 shGPM6A tdTomato GB cells during live cell imaging in neuron-tumor cocultures (n = 96 S24 cells, n = 136 S24 shGPM6A GB cells).

**Table S1.** List of Antibodies, Reagents, Bacterial and viral strains, Biological samples, Assays, Cell lines, Software, Chemicals used in this study.

| REAGENT or RESOURCE | SOURCE | IDENTIFIER |
| --- | --- | --- |
| **Antibodies** |  |  |
| Homer 1/2/3 rabbit polyclonal | Synaptic Systems | Cat#160103; RRID: AB_10694096 |
| Mouse monoclonal anti-nestin | Abcam | Cat#ab22035; RRID:AB_446723 |
| Mouse monoclonal Anti-Puromycin | Merck | Cat# MABE343; RRID: AB2566826 |
| Rabbit monoclonal anti-S6 ribosomal protein | Cell Signaling Technology | Cat# 2217; RRID:AB_331355 |
| Rabbit polyclonal anti-Synapsin1/2 | Synaptic Systems | Cat# 106 003; RRID:AB_2619773 |
| Chicken polyclonal anti-GFP | Abcam | Cat# a13970; RRID: AB_300798 |
| Guinea pig polyclonal anti-RFP | Synaptic Systems | Cat# 390004; RRID: AB_2737052 |
| Rabbit monoclonal anti-nestin | Abcam | Cat# ab105389; RRID: AB_10859398 |
| Goat anti-chicken Alexa 488 | Invitrogen | Cat# A32931; PRID:AB_2762843 |
| Goat anti-rabbit Alexa647 | Invitrogen | CAT # A-21245; RRID:AB_2535813 |
| Goat anti-guinea pig Alexa 546 | Invitrogen | Cat# A11011, AB_2534118; Cat# A-11074; RRID:AB_2534118 |
| Goat anti-mouse Alexa 488 | Invitrogen | Cat# A11017, RRID: AB_2534069 |
| Goat anti-rabbit Alexa 488 | Invitrogen | Cat# A11008, RRID: AB_143165 |
| abberior STAR 580, goat anti-rabbit IgG | abberior | ST580-1002-500UG |
| abberior STAR RED, goat anti-mouse IgG | abberior | STRED-1001-500UG |
| Phalloidin Labeling Probe Alexa Fluor™ 488 | Thermo Fisher Scientific | Cat#A12379; RRID: N/A |
| **Bacterial and virus strains** |  |  |
| pLego-SFFV-shGPM6A-tdtomato | This paper | N/A |
| pLKO1.1-puro-CMV-shGAP43-GFP | Osswald et al. 2015 (8) | N/A |
| Biological samples |  |  |
| Human cerebrospinal fluid | This paper | N/A |
| **Chemicals, peptides, and recombinant proteins** |  |  |
| 0,05% Trypsin-EDTA (1x) | Gibco | Cat#25300-054 |
| 2.5% Trypsin (10x) | Gibco | Cat#15090-046 |
| 4-Hydroxy-TEMPO | Merck | Cat#176141 |
| Accutase Solution | Thermo Fisher Scientific | Cat#A1110501 |
| Acheson silver | Plano | Cat#3692 |
| Acrylamide | Sigma-Aldrich | Cat#A9099 |
| AcX | Thermo Fisher Scientific | Cat#A20770 |
| Ammonium persulfate | Sigma-Aldrich | Cat#A3678 |
| Anisomycin | Sigma-Aldrich | Cat#A9789 |
| Aqua | B Braun | Cat#0082479E |
| B-27 supplement for neuronal co-culture | Gibco | Cat#17504044 |
| B27 supplement without Vitamin A for monoculture | Gibco | Cat#12587010 |
| Biotin-PEG(4)-alkyne | Iris Biotech | Cat#PEG4950 |
| CaCl<sub>2</sub> | neoFroxx | Cat#LC-5912.1 |
| Cacodylic acid-Na-salt-3H<sub>2</sub>O | Serva | Cat#15540.1 |
| Cell Conditioning Solution (CC2) | Roche | Cat#5279798001 |
| Choline Bicarbonate | Sigma-Aldrich | Cat#C7519-100ML |
| Cycloheximide | Roth | Cat#8682.1 |
| DAB+Chromogen | Dako | Cat#K3468 |
| DAPI | Sigma-Aldrich | Cat#D9542 |
| DAPI | Invitrogen | Cat#D9542-10MG |
| DBA HardenerHärter | Carl Roth | Cat#8623.232.2 |
| DDSA | Serva | Cat#20755.02 |
| Dimethylsulfoxide | Pan Biotech | Cat#P60-36720100 |
| DMEM F12 Medium | Gibco | Cat#11330032 |
| DMEM High Glucose | anprotec | Cat#AC-LM-0014 |
| DMEM, high glucose, no glutamine, no methionine, no cystine | Gibco | Cat#21013024 |
| DMP-30 | Carl Roth | Cat#8621.1 |
| Dulbecco's Phosphate Buffered Saline | Sigma | Cat#D8537-500ML |
| EDTA | Thermo Fisher Scientific | Cat#AM9260G |
| Ethanol | Sigma-Aldrich | Cat#32221-M |
| Evans Blue | Sigma-Aldrich | Cat#E2129 |
| Fetal Bovine Serum (FBS) | Thermo Fisher Scientific | Cat#A3840401 |
| Fetal Bovine Serum (FBS) | anprotec | Cat#AC-SM-0027 |
| Fluorescein isothiocyanate-dextran (FITC) | Sigma-Aldrich | Cat#FD2000S-1G |
| Glucose | Carl Roth | Cat#X997.2 |
| Glycid Ether 100 for electron microscopy | Serva | Cat#21045.02 |
| HBSS | anprotec | Cat#AC-BS-0009 |
| Heparin | Sigma-Aldrich | Cat#H4784 |
| human EGF recombinant | anprotec | Cat#AC-RH-0633 |
| human FGF recombinant | anprotec | Cat#AC-RH-0794 |
| Insulins solution human | Sigma-Aldrich | Cat#I9278 |
| KCl | Thermo Fisher Scientific | Cat#196770010 |
| KH<sub>2</sub>PO<sub>4</sub> | Sigma-Aldrich | Cat#P0662-500G |
| L-glutamine (GlutaMAX™ - I (100x)) | Gibco | Cat#35050061Cat#11330032 |
| L-Ascorbinsäure | Sigma-Aldrich | Cat#A5690 |
| Magnesium ATP | Sigma-Aldrich | Cat#A9187 |
| MgCl<sub>2</sub> | Carl Roth | Cat#2189.2 |
| Matrigel | Corning | Cat#354230 |
| MNA hardener | Carl Roth | Cat#8639.2 |
| N,N'-Methylenbisacrylamide | Merck | Cat#146072 |
| NaCl | Sigma-Aldrich | Cat#S7653 |
| Nail Polish | Various Suppliers | Cat#N/A |
| Narcoren | Boehringer Ingelheim<br>VETMEDICA | Cat#11336163 |
| Neurobasal medium | Gibco | Cat#11570556 |
| N-Methyl-D-Glucamin | Sigma-Aldrich | Cat#M2004-1KG |
| NucleoSpin RNA Plus kit | Macherey-Nagel | Cat#740984.50 |
| Osmium tetroxide | Serva | Cat#31253.04 |
| PEG-it Virus Precipitation Solution | Systems Biosciences | Cat#LV810A-1 |
| PFA 4% | Roth | Cat#P087.3 |
| Polybrene | Merck | Cat#TR-1003-G |
| Poly-L-Lysine solution | Sigma-Aldrich | Cat#P4832-50ml |
| Potassium ferricyanide | Sigma-Aldrich | Cat#1548258 |
| Propylenoxide | VWR Chemicals | Cat#27165.295 |
| Proteinase-K | New England Biolabs | Cat#P8107 |
| Puromycin-Dihydrochloride | Gibco | Cat#A11138-03 |
| Puroswitch | Ko, T. et al 2022 (43) | N/A |
| SlowFade™ Gold Antifade Mountant | Thermo Fisher Scientific | Cat#S36936 |
| Sodiumacrylate | Sigma-Aldrich | Cat#408220 |
| Sucrose | Sigma-Aldrich | Cat#S0389 |
| Sulforhodamine 101 (SR101) | Sigma-Aldrich | Cat#S7635 |
| Sulforhodamine 101 (SR101) | Invitrogen | Cat#S359 |
| SYTO ® RNaselect™ Green<br>Fluorescent Cell Stain | Thermo Fisher Scientific | Cat#S32703 |
| TEMED | Merck | Cat#T9281 |
| Tetramethylrhodamine isothiocyanate<br>dextran (TRITC) | Sigma-Aldrich | Cat#52194-1G |
| THPTA | Jena Bioscience | Cat#CLK-1010-100 |
| Tris-Cl | Thermo Fisher Scientific | Cat#AM9856-500ml |
| Triton™ X-100 | Sigma-Aldrich | Cat#T9284-100ML |
| TRIzol Reagent | Thermo Fisher Scientific | Cat#15596026 |
| Trypan Blue | Sigma-Aldrich | Cat#T8154 |
| Uranyl acetate | Serva | Cat#77870 |
| Vectastain ABC-Kit | Linaris/Vector | Cat#PK-4000 |
| Xylene | Thermo Fisher Scientific | Cat#L13317.AU |
| **Critical commercial assays** |  |  |
| Xenium Human brain gene expression panel | 10x Genomics | https://www.10xgenomics.com/datasets |
| NucleoSpin RNA kit | Macherey-Nagel | Cat#740984.50 |
| Infinium MethylationEPIC array kit | Illumina | 20087706 |
| High Sensitivity RNA ScreenTape | Agilent | 5067-5579 |
| iTaq™ Universal SYBR®Green One-Step Kit | Biorad | #1725151 |
| Duolink® In Situ Orange Starter Kit Mouse/Rabbit | Sigma-Aldrich | DUO92102 |
| **Experimental models: Cell lines** |  |  |
| S24 | This paper | N/A |
| T269 | This paper | N/A |
| P3XX | This paper | N/A |
| GG16 | This paper | N/A |
| BG5 | This paper | N/A |
| BG7 | This paper | N/A |
| NCH421K | Cytion | 300118 |
| NCH644 | Cytion | 300124 |
| U3047MG | Xie et al., 2015 (121) | N/A |
| U3048MG | Xie et al., 2015 (121) | N/A |
| HF2587 | Giladi et al., 2015 (122) | N/A |
| HF2927 | deCarvalho et al., 2018 (123) | N/A |
| HF3016 | deCarvalho et al., 2018 (123) | N/A |
| HF3253 | deCarvalho et al., 2018 (123) | N/A |
| **Experimental models: Organisms/strains** |  |  |
| NMRI-Foxn1 nu/nu | Charles River and Janvier | BL210203171 |
| WISTAR | Janvier | N/A |
| **Recombinant DNA** |  |  |
| Plasmid: mGFP | De Paola et al., 2003 (124)<br>Dondzillo et al., 2010 (125) | N/A |
| Plasmid: tdTomato | Osswald et al., 2015 (8) | N/A |
| Plasmid: shGAP43 GFP | Osswald et al., 2015 (8) | N/A |
| Plasmid: shGPM6A tdTomato | This paper | N/A |
| Plasmid: GCamp7-tdTomato | Dana et al., 2019 (126) | N/A |
| Software and algorithms |  |  |
| ImageJ/Fiji | Schindelin et al., 2012 (106) | https://imagej.nih.gov/ij/ |
| NIS-Elements AR Analysis 5.30.01 64-bit & 5.34.01 | Nikon | https://www.microscope.healthcare.nikon.com/products/software |
| Ilastik 1.4.0 | Berg et al., 2019 (110) | https://www.ilastik.org/development.html |
| Arivis Vision4D 3.5.0 | Arivis AG, Munich, Germany | https://imaging.arivis.com/en/imaging-science/arivis-vision4d |
| R Studio 1.4 | RStudio Team, 2020 | NA |
| GraphPad Prism Version 10.6.1 | GraphPad | NA |
| Adobe Illustrator 25.4.1 | Adobe | NA |
| Adobe Photoshop | Adobe | NA |
| DaVinciResolve 17 | Blackmagicdesign | https://www.blackmagicdesign.com/de/products/davinciresolve/ |
| Leica Application Suite X | Leica Microsystems CMS GmbH | https://www.leica-microsystems.com/de/produkte/mikroskop-software/p/leica-las-x-ls/ |
| TrakEM2 | Cardona et al., 2012 (127) | N/A |
| Atlas 5.3.0.25 | Zeiss | N/A |
| neuTube | Feng et al., 2015 (111) | https://neutracing.com/ |
| Simple Neurite Tracer (SNT) | Arshadi et al., 2021 (112) | https://www.mdc-berlin.de/research/publications/snt-unifying-toolbox-quantification-neuronal-anatomy/ |
| MTrackJ | Meijering et al., 2012 (108) | https://imagescience.org/meijering/software/mtrackj/ |
| Bio-Formats | Linkert et al., 2010 (128) | N/A |
| AQuA2 | Xuelong Mi et al., 2025 (113) | https://github.com/yu-lab-vt/AQuA2 |
| Zen Blue 3.5, 3.7 and 3.9 | Zeiss | RRID:SCR_013672 |
| MERSCOPE Visualizer | Vizgen, Inc. (2024). | https://vizgen.com/vizualizer-software/ |
| Xenium Explorer | 10x Genomics | https://www.10xgenomics.com/support/software/xenium-explorer/latest |
| MATLAB 9.120.1884302 | MathWorks | RRID:SCR_001622 |
| Kallisto | Bray et al. 2016 (129) | N/A |
| **Other** |  |  |
| Ceramic beads (for RNA extraction) |  |  |
| Glass Bottom Plates for imaging | Cellvis | P06-1.5H-N, P12-1.5H-N, P24-1.5H-N, P96-1.5H-N |
| Millicell Cell Culture Inserts, 0.4µm, 30mm Diameter | Merck | Cat#PICM0RG50 |
| SEM pin stub (0,5"/6mm length) | Agar scientific | Cat#G301F |
| THINCERT cell culture insert | Greiner | Cat#657631 |
| 10µl syringe, Special Cemented Needle | Hamilton | Cat#80308 |

**Table S2:** Patient derived GB-lines and pediatric-type diffuse high grade glioma lines encompassed in this study.

| Patient derived cell line | Sex | Age | MGMT meth. | CDKN2A/B | Treatment |
| --- | --- | --- | --- | --- | --- |
| S24 | female | 44 years | meth. | hom. del. | Rx + TMZ |
| T269 | male | 68 years | meth. | hom. del. | none prior |
| P3XX | male | 64 years | meth. | hom. del. |  |
| GG16 |  |  | meth. | hom. del. |  |
| BG5 | female | 72 years | unmeth. | hom. del. | none prior |
| BG7 |  |  | meth. | hom. del. |  |
| NCH421 | male | 66 years | meth. | het. del |  |
| NCH644 | female | 66 years | meth. | bal. |  |
| U3047 | female | 66 years | meth. | het. del. | none prior |
| U3048 |  | 77 years | meth. | hom. del. | none prior |
| HF2587 | female | 56 years | meth. | hom. del. | none prior |
| HF2927 | female | 55 years | meth. | hom. del. | none prior |
| HF3253 | female | 82 years | meth. | hom. del. | none prior |
| HF3016 | male | 45 years | unmeth. | unaltered | none prior |
| SJ-HGGX97 | male | 7 months |  |  |  |
| OPBG-GBM-001 | male | 12 years |  |  |  |
| B062-039 |  |  |  |  |  |
| HSJD-GBM-002 | male | 14 years |  |  |  |
| RCMB47 | male | 15 years |  |  | Radiotherapy |

meth. = methylated; unmeth. = unmethylated; hom. del. = homozygous deletion; het. del. = heterozygous deletion; bal. =balanced translocation; Rx + TMZ = treatment with Temozolomide

**Table S3:**
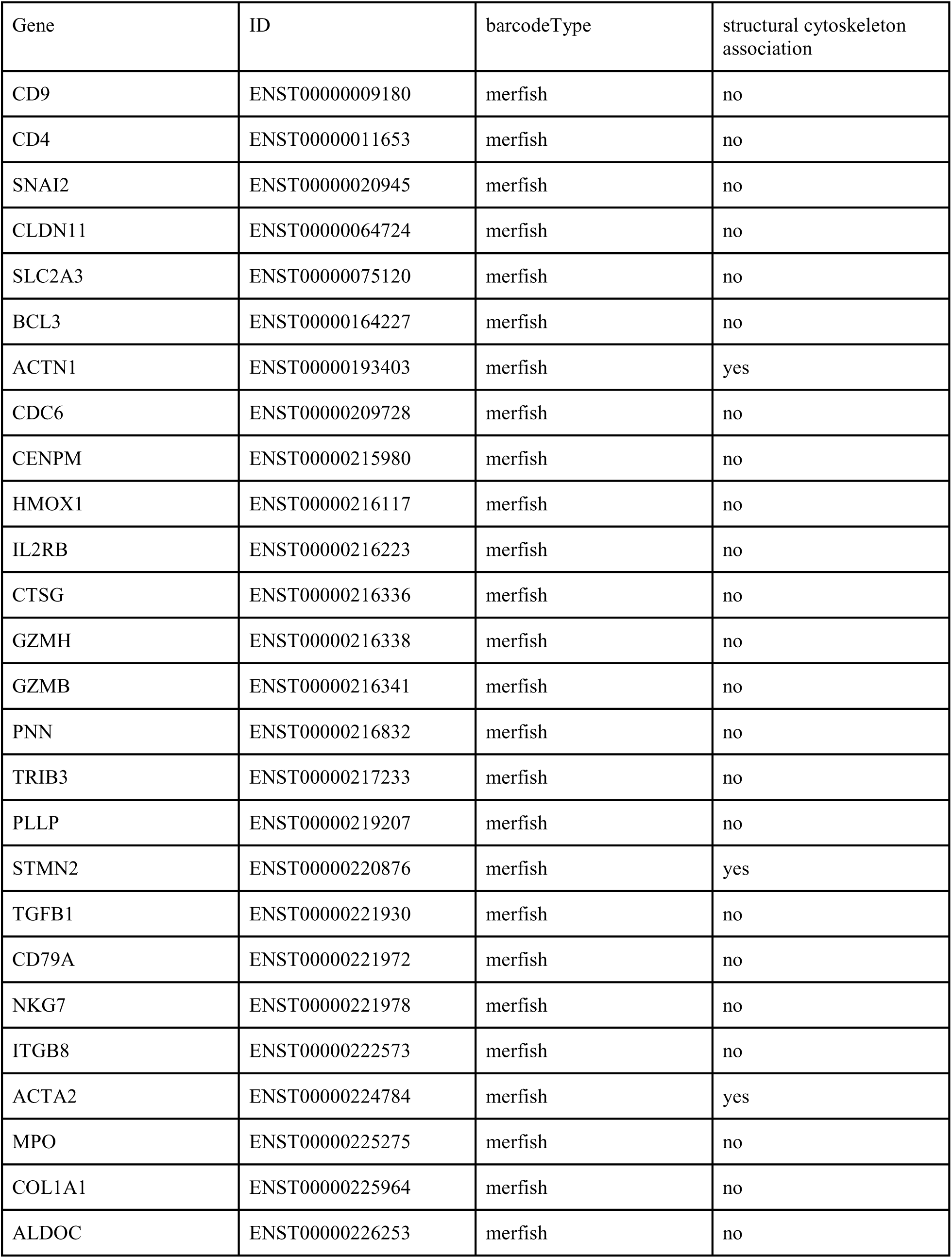
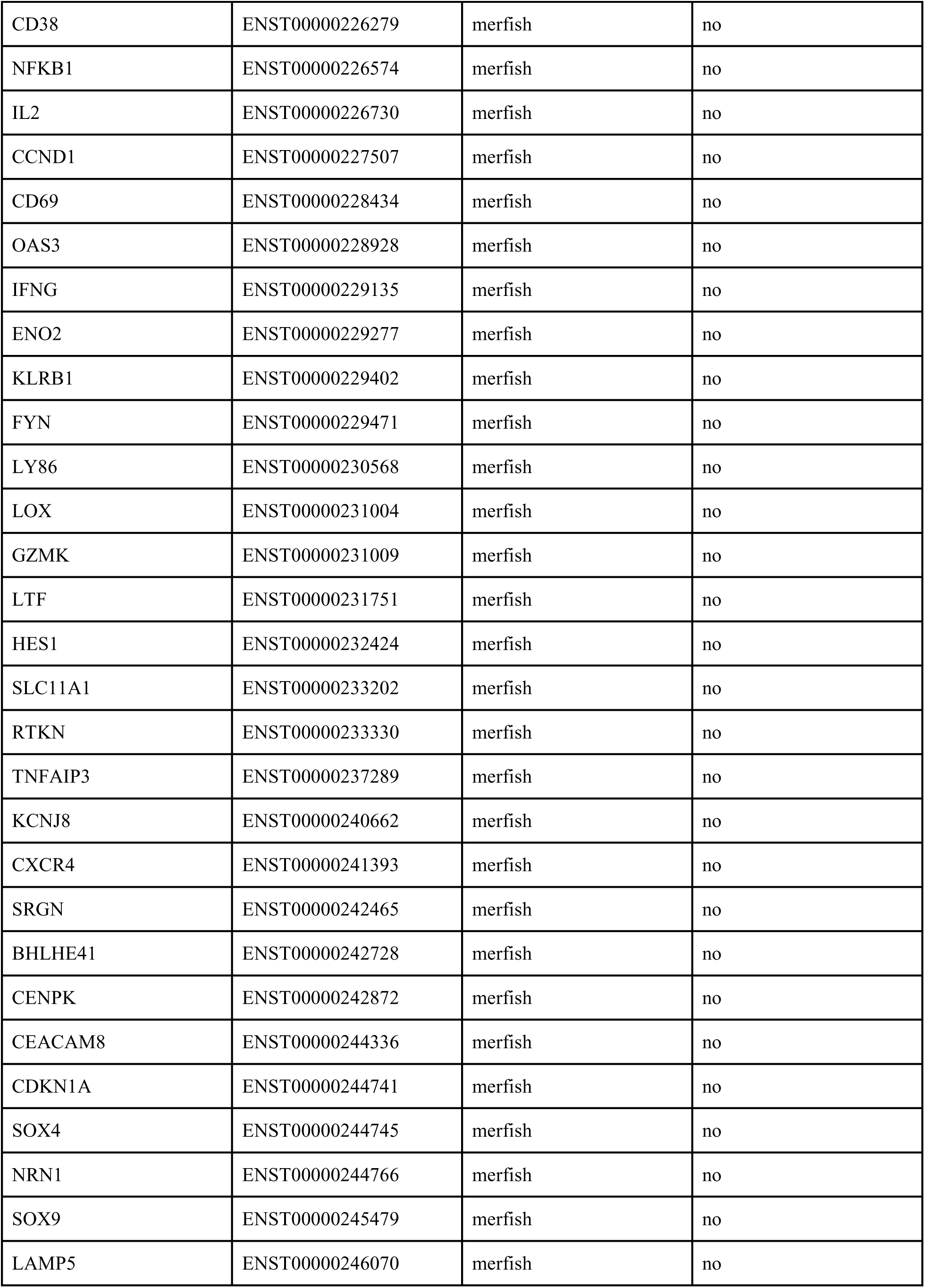
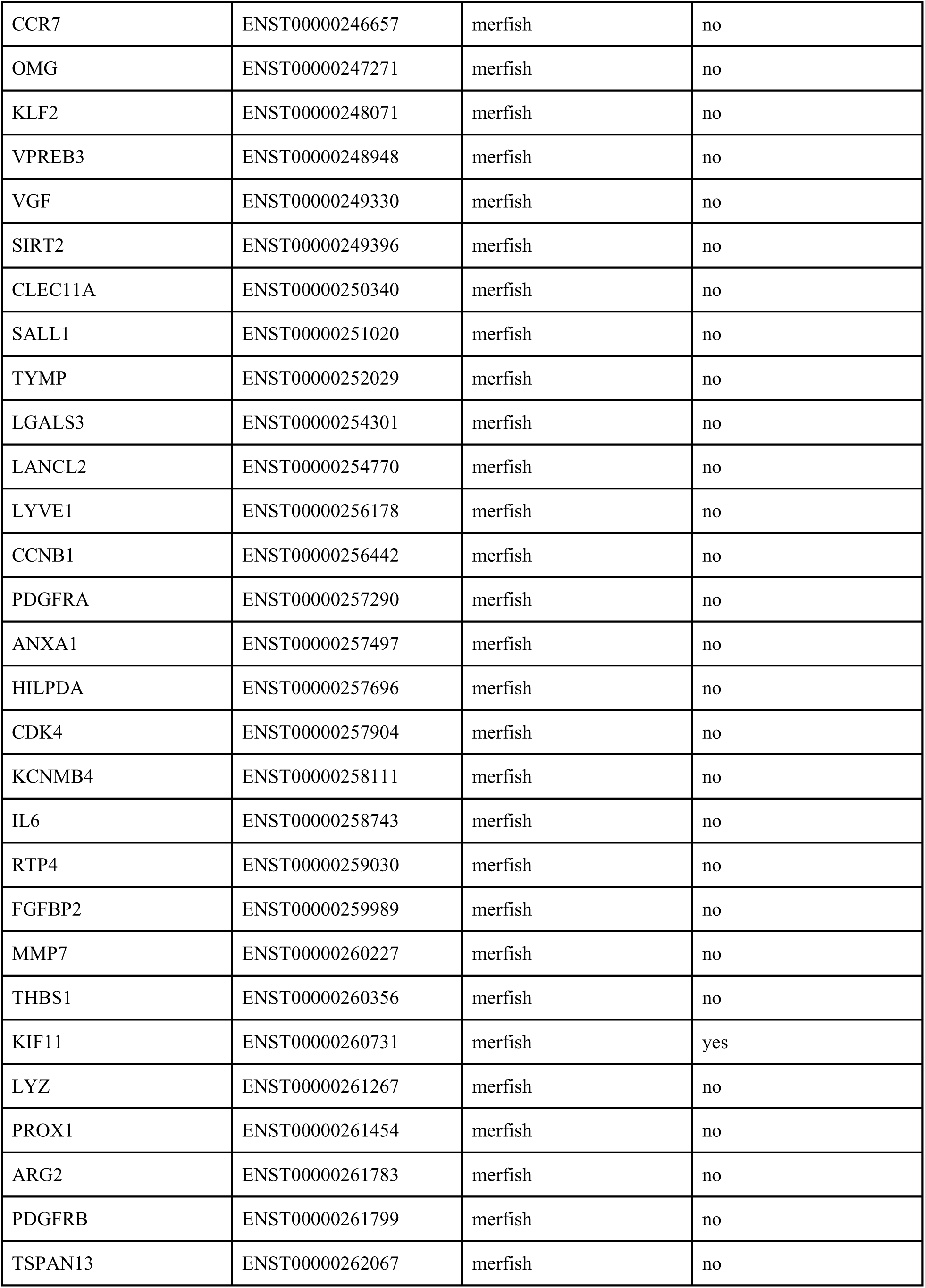
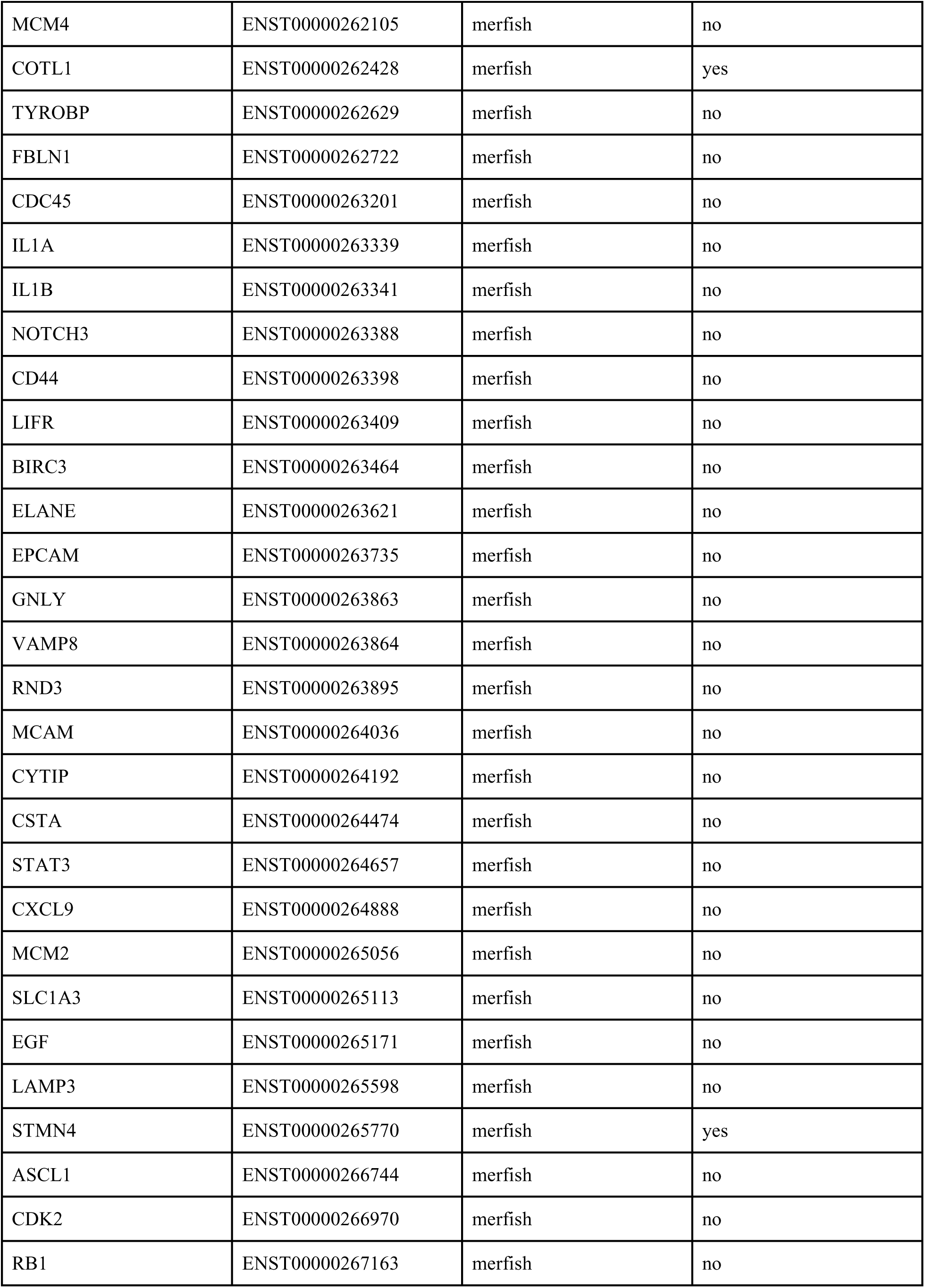
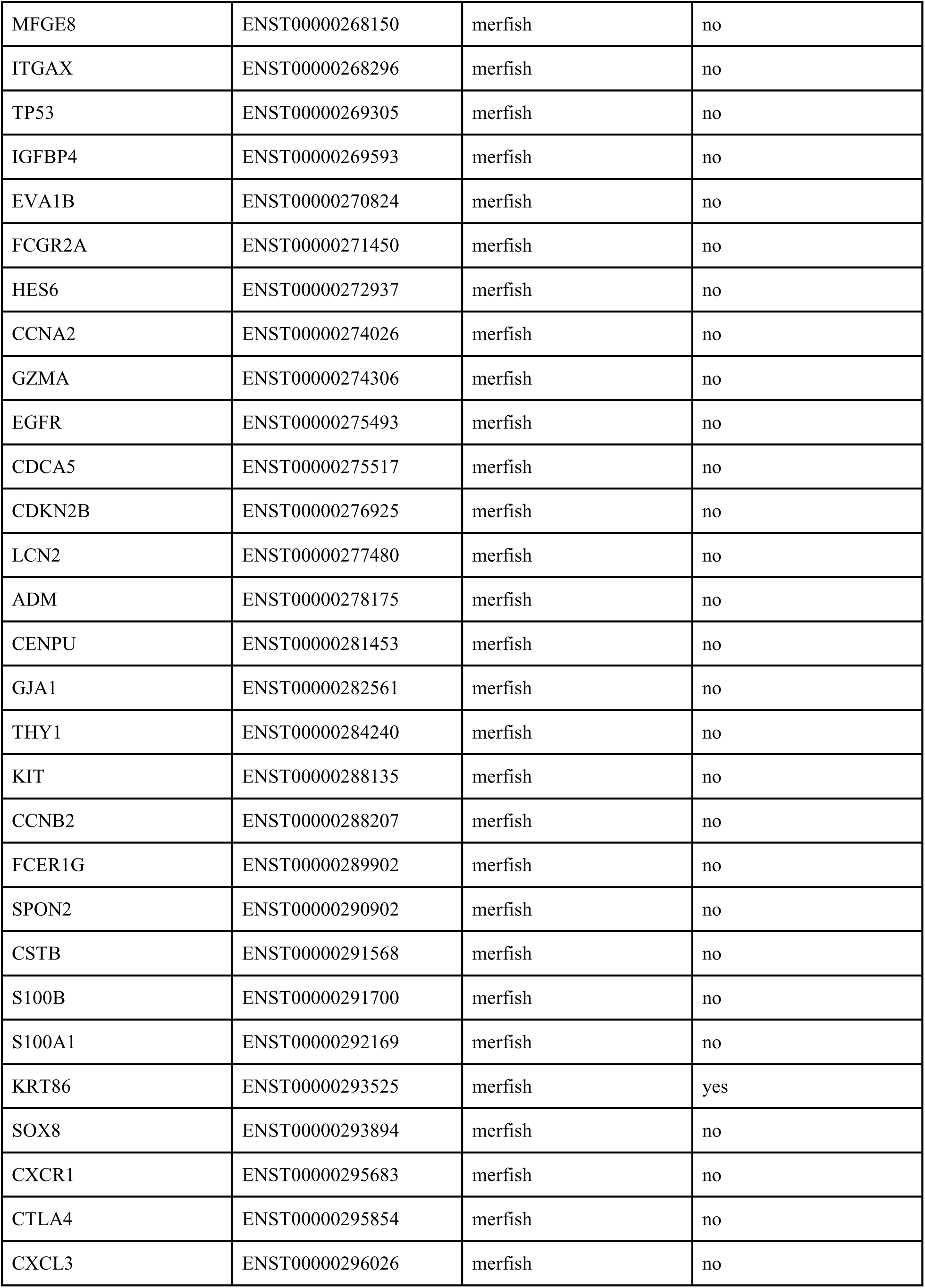
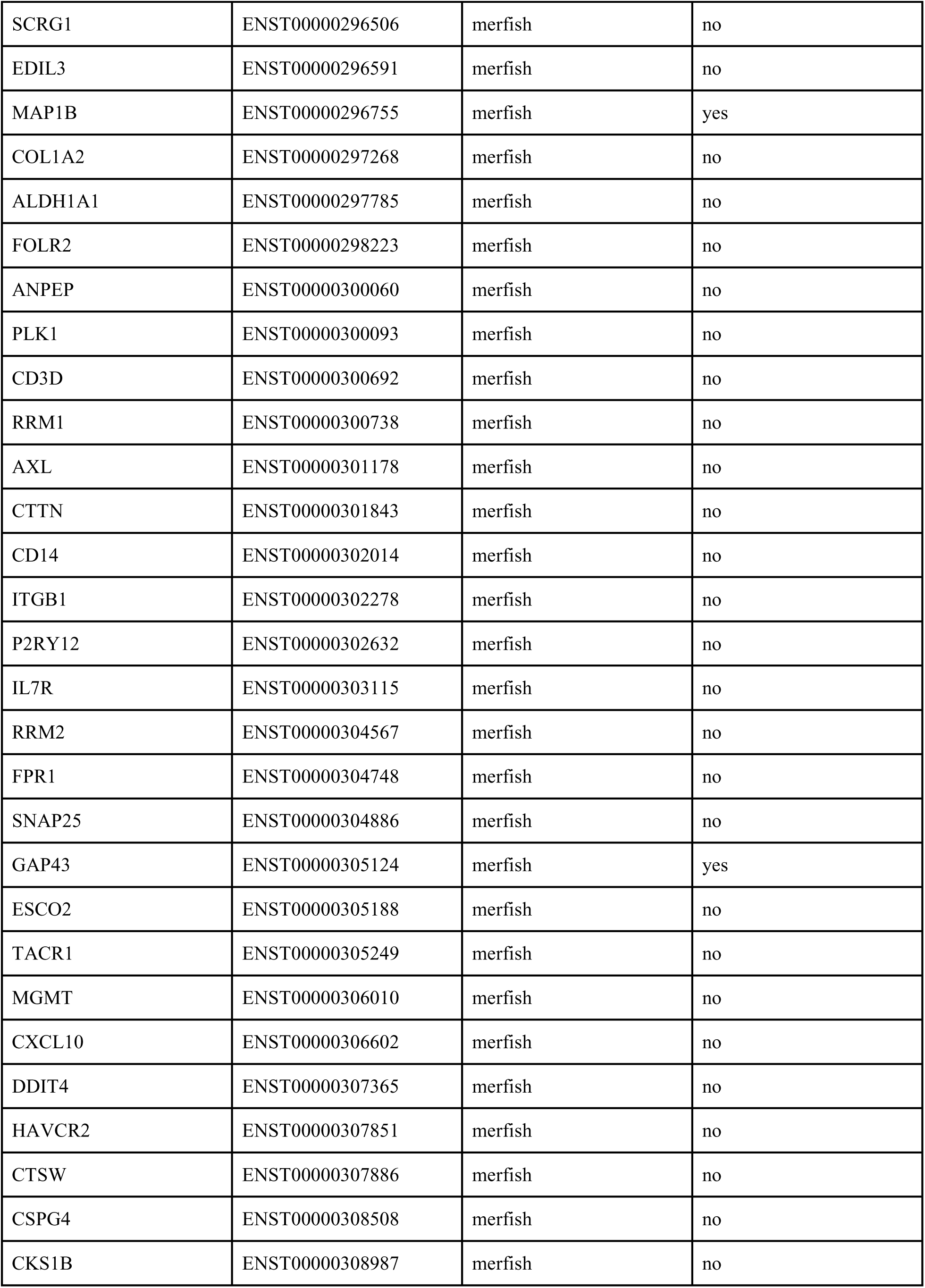
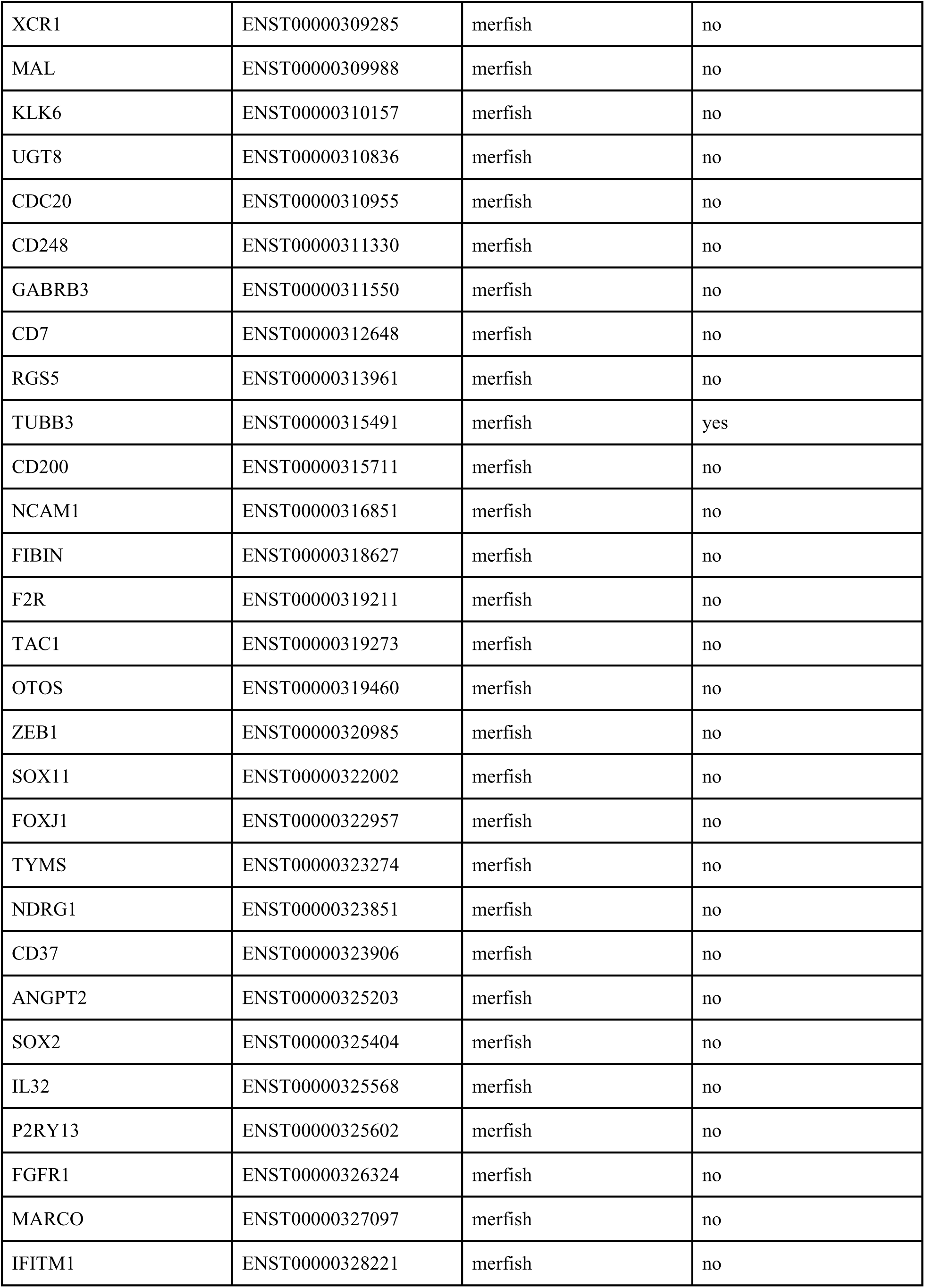
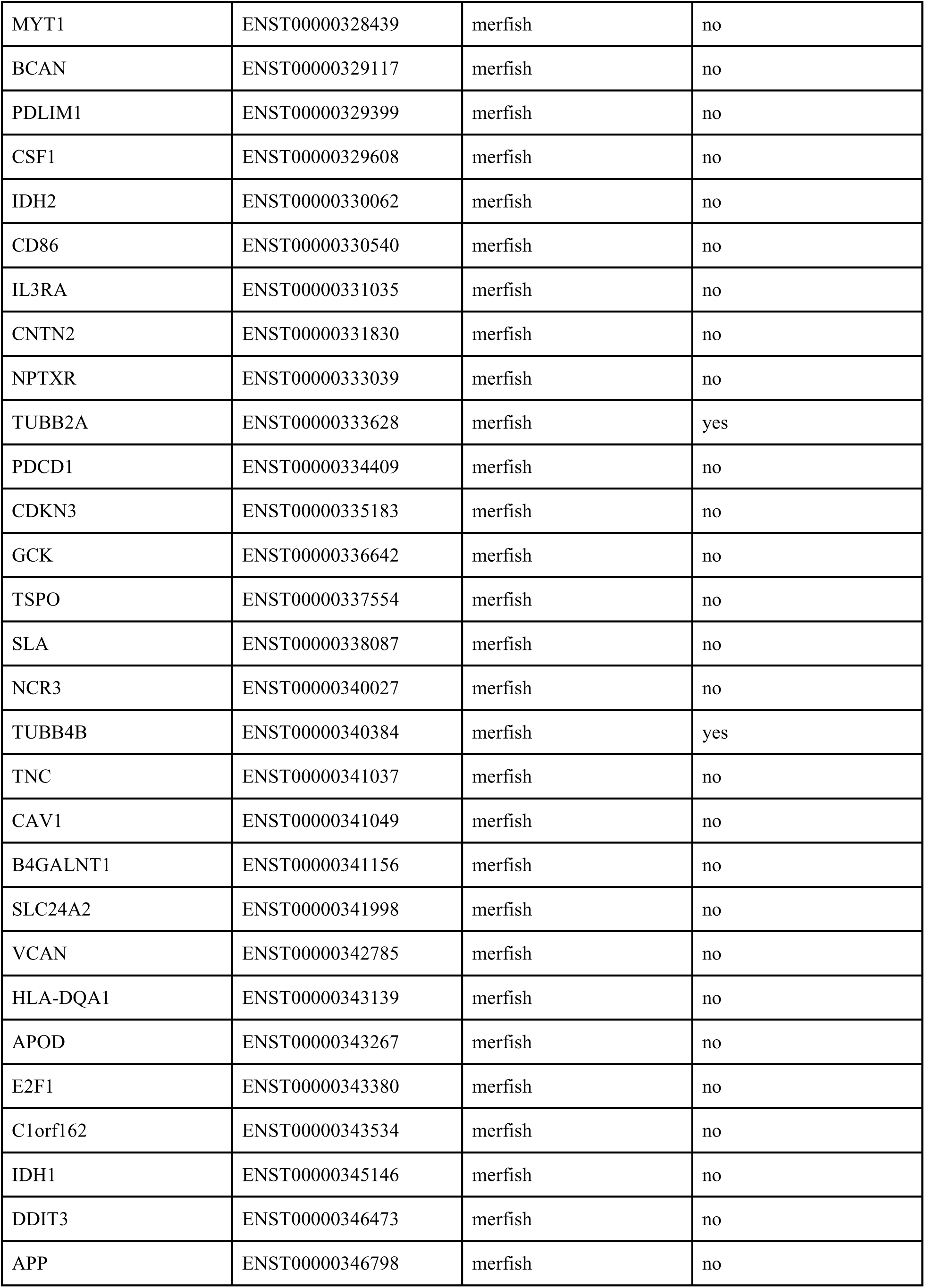
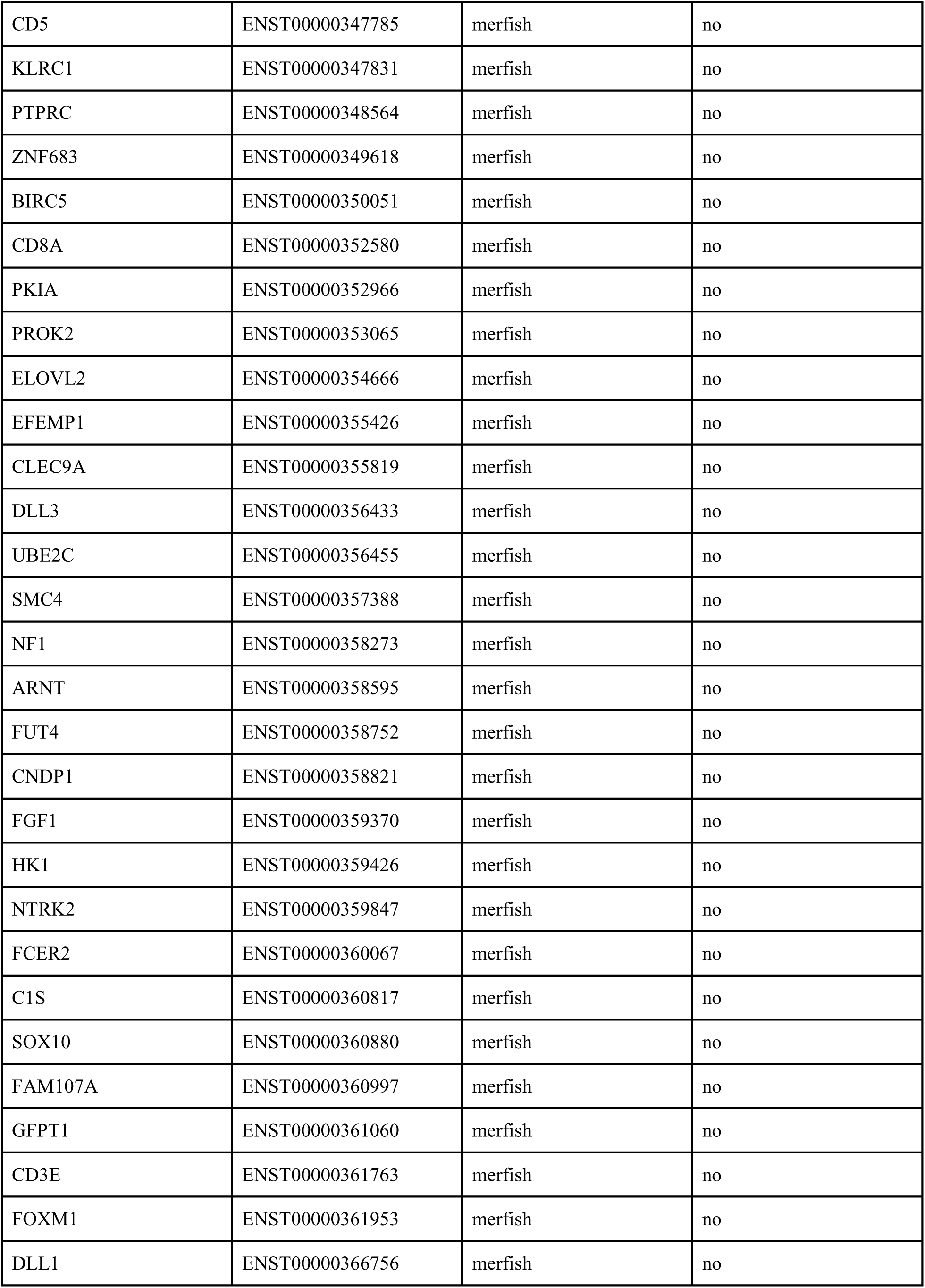
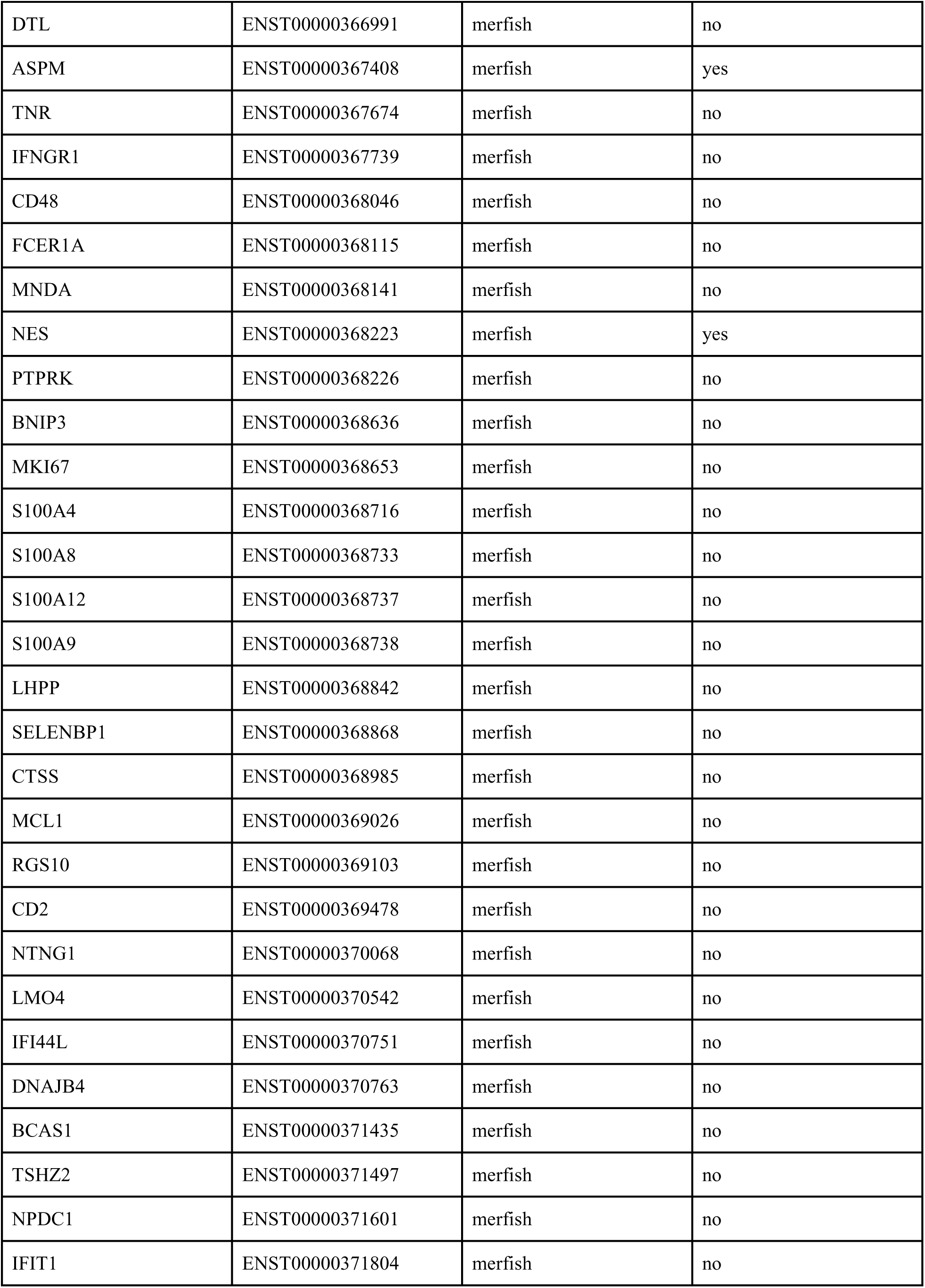
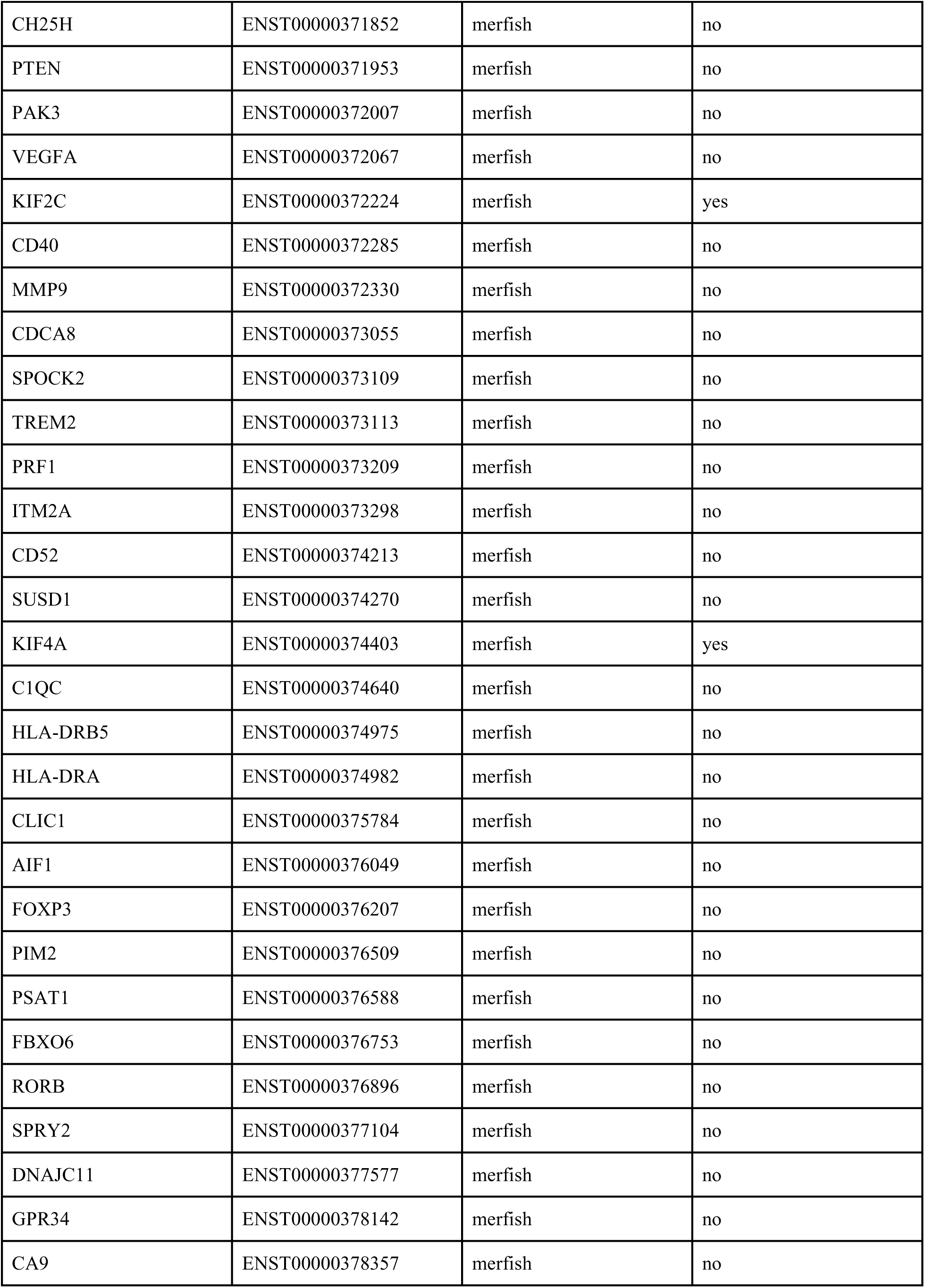
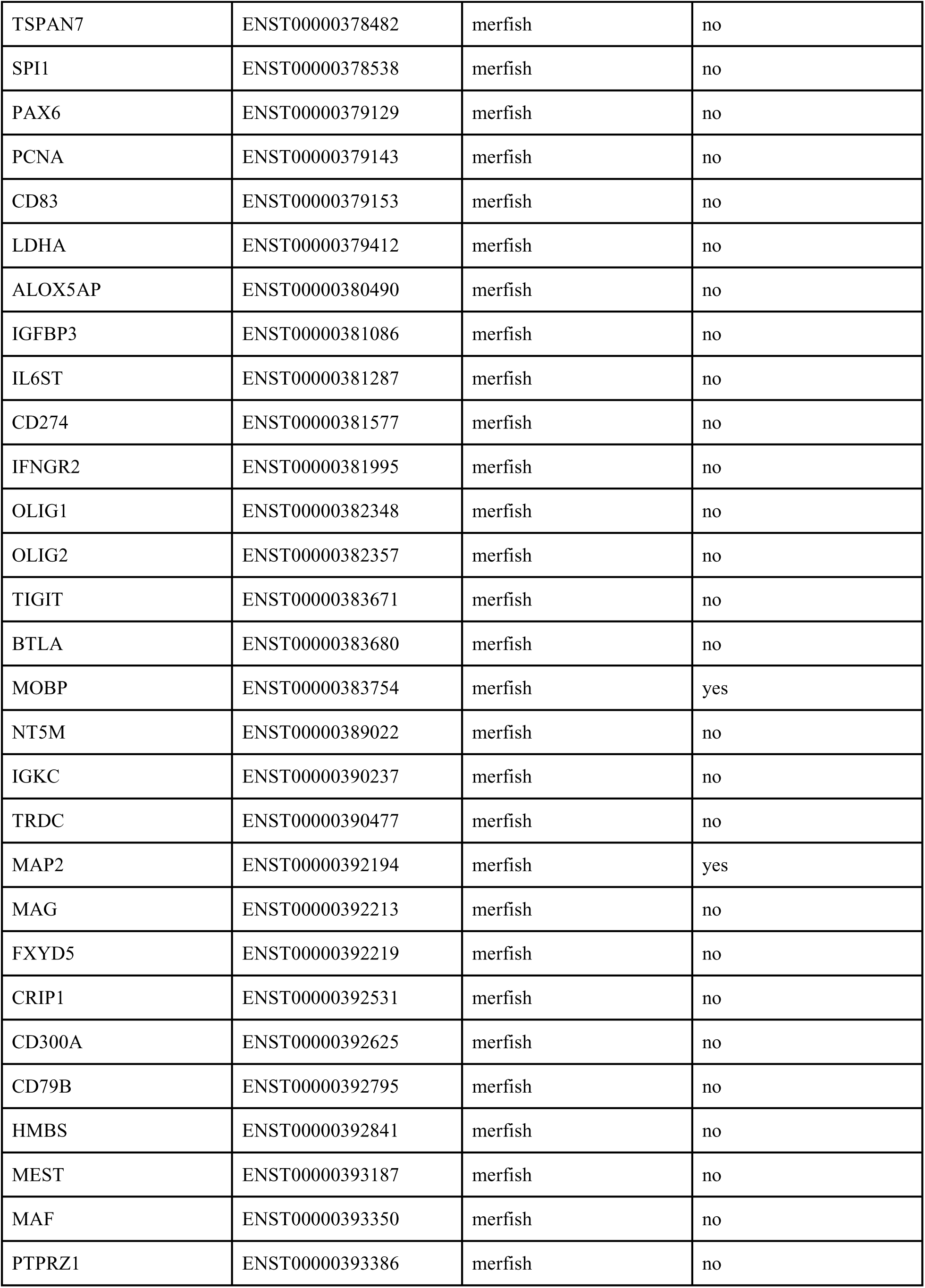
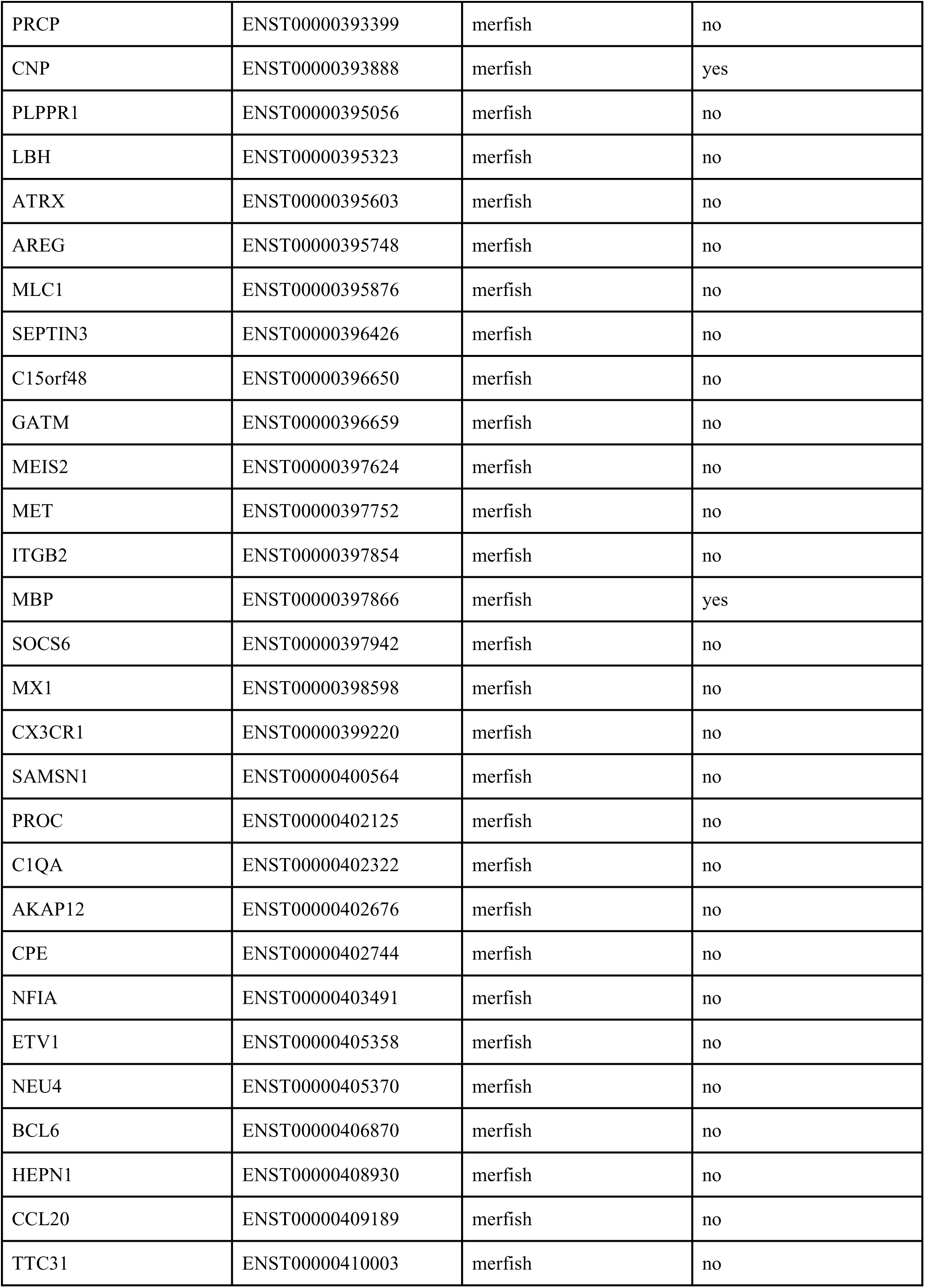
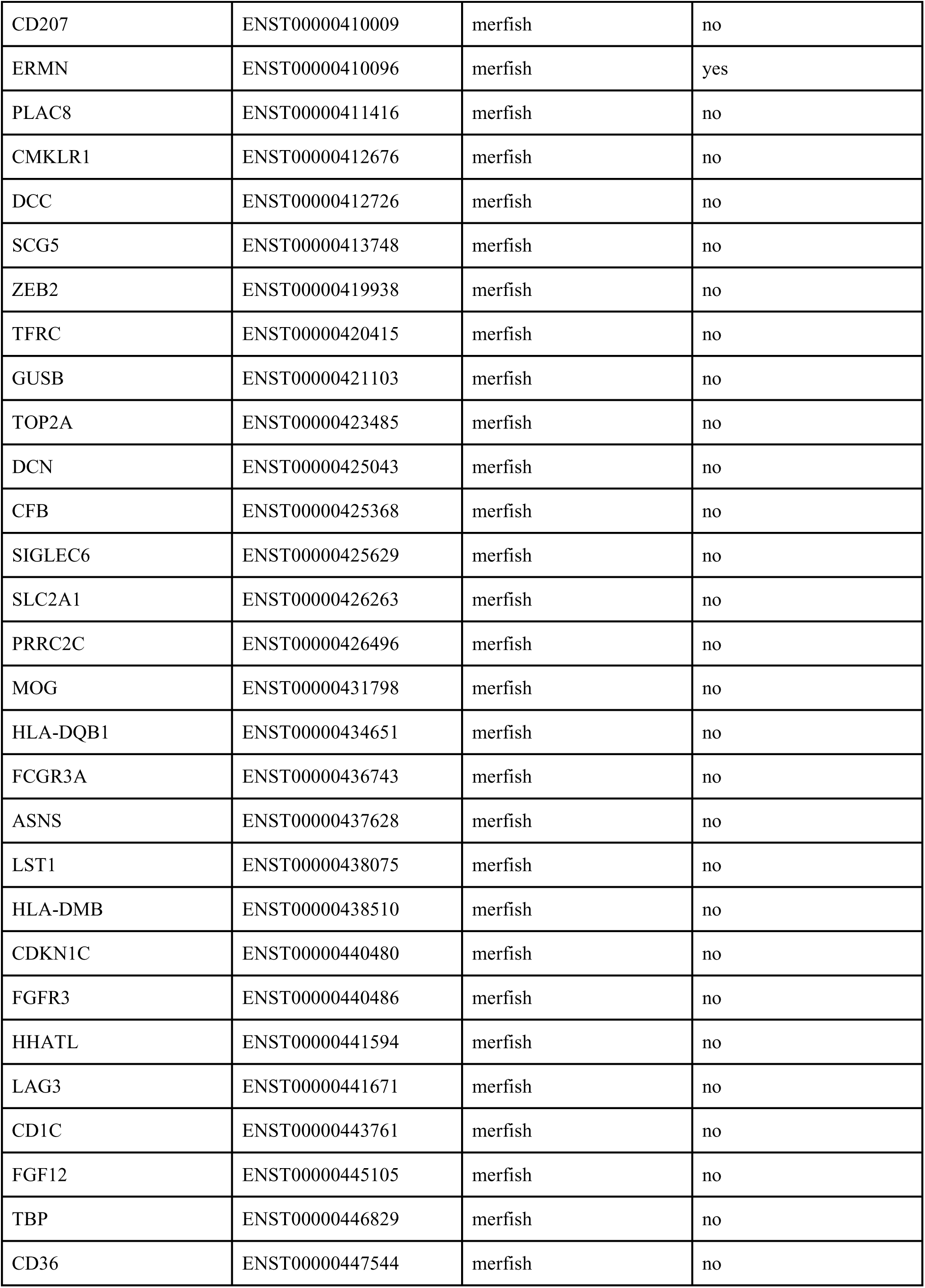
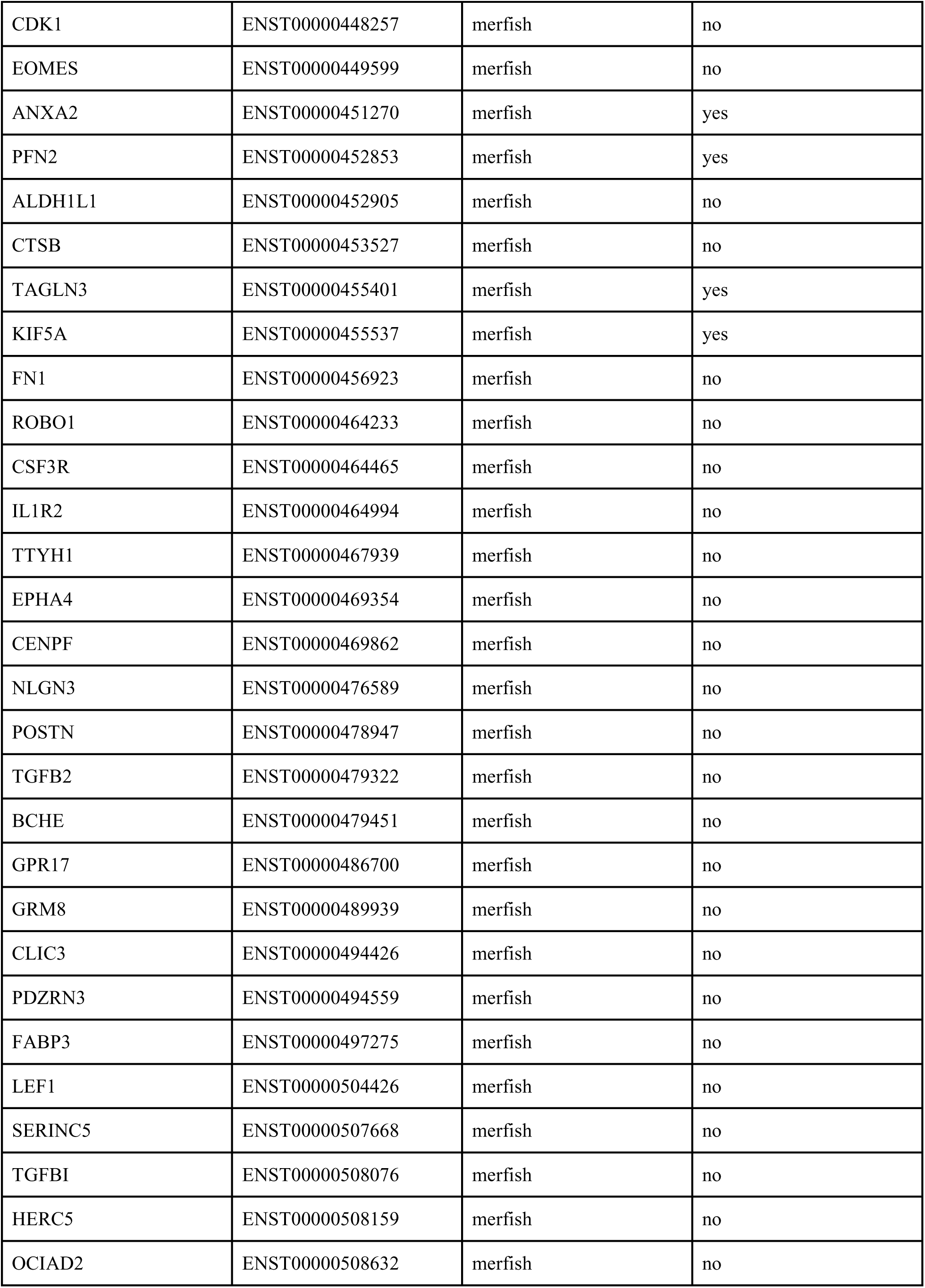
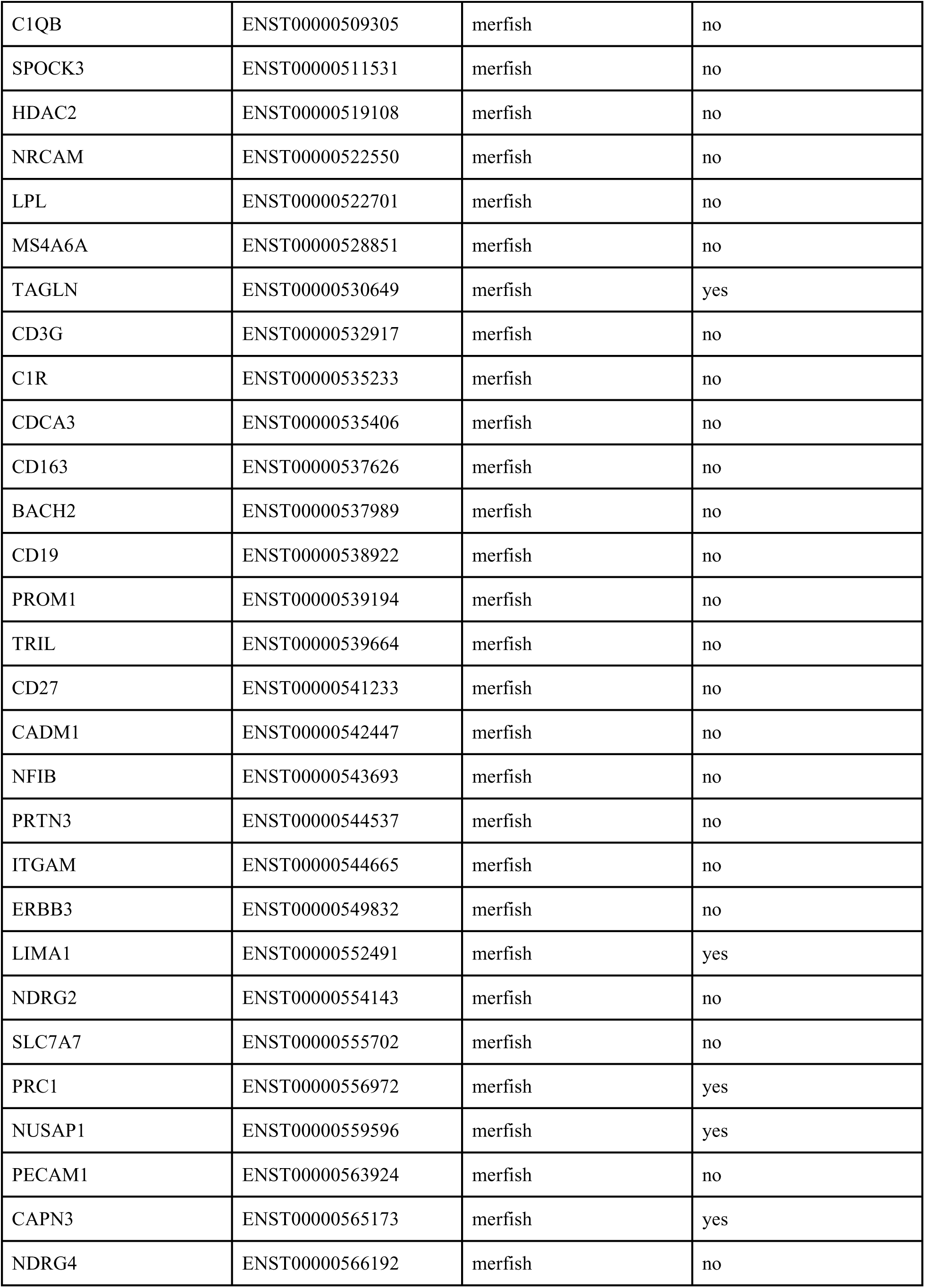
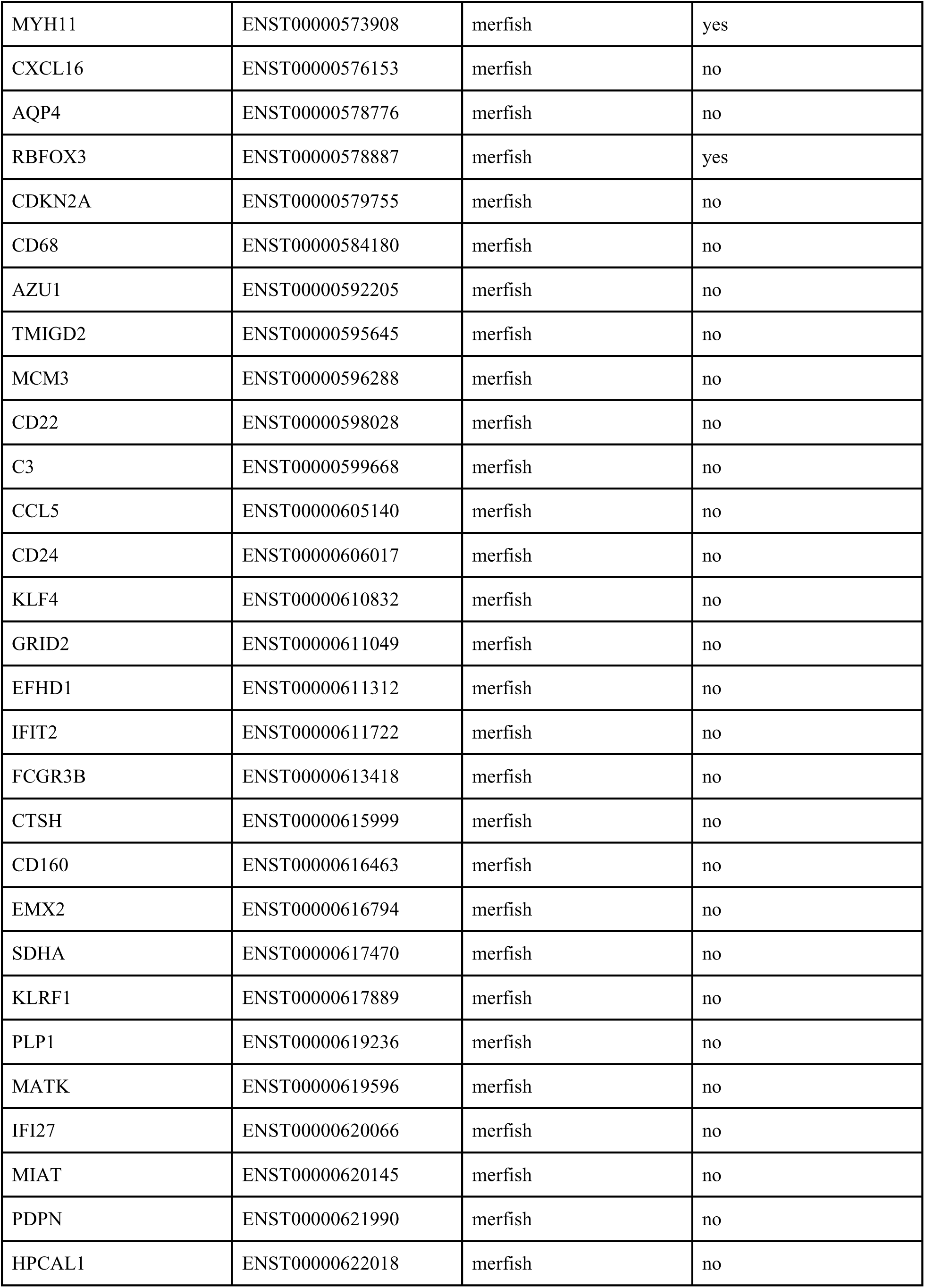
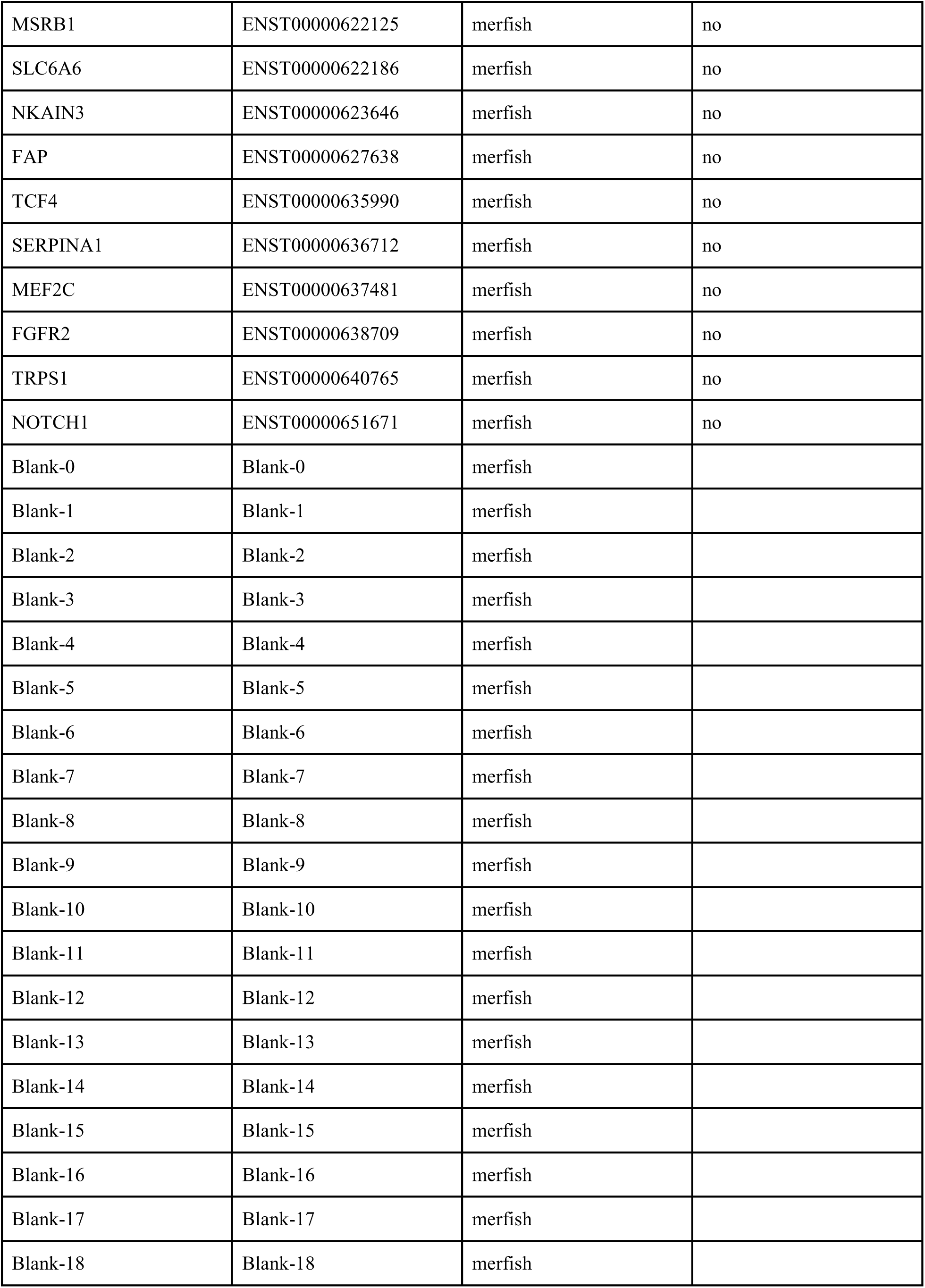
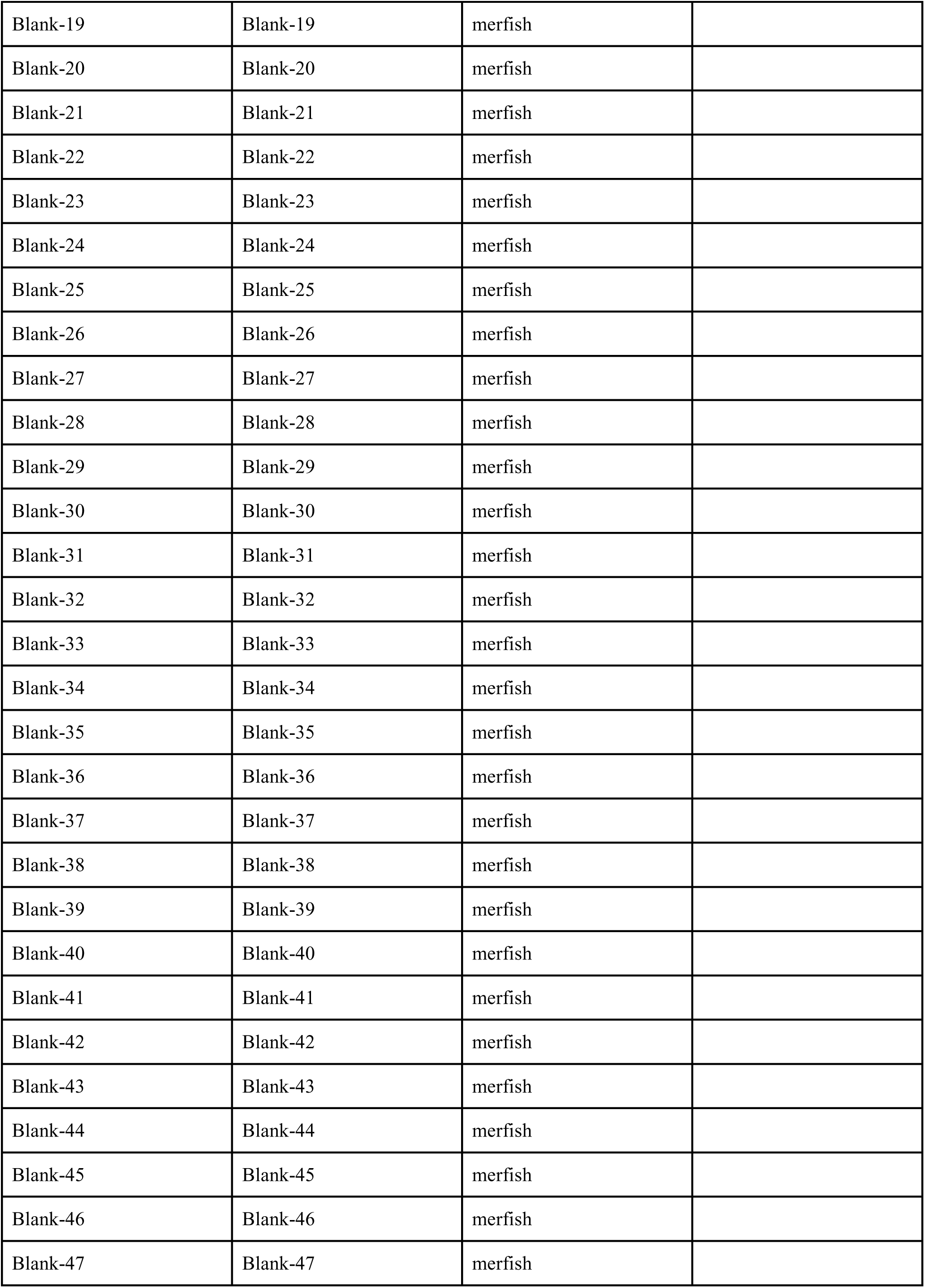
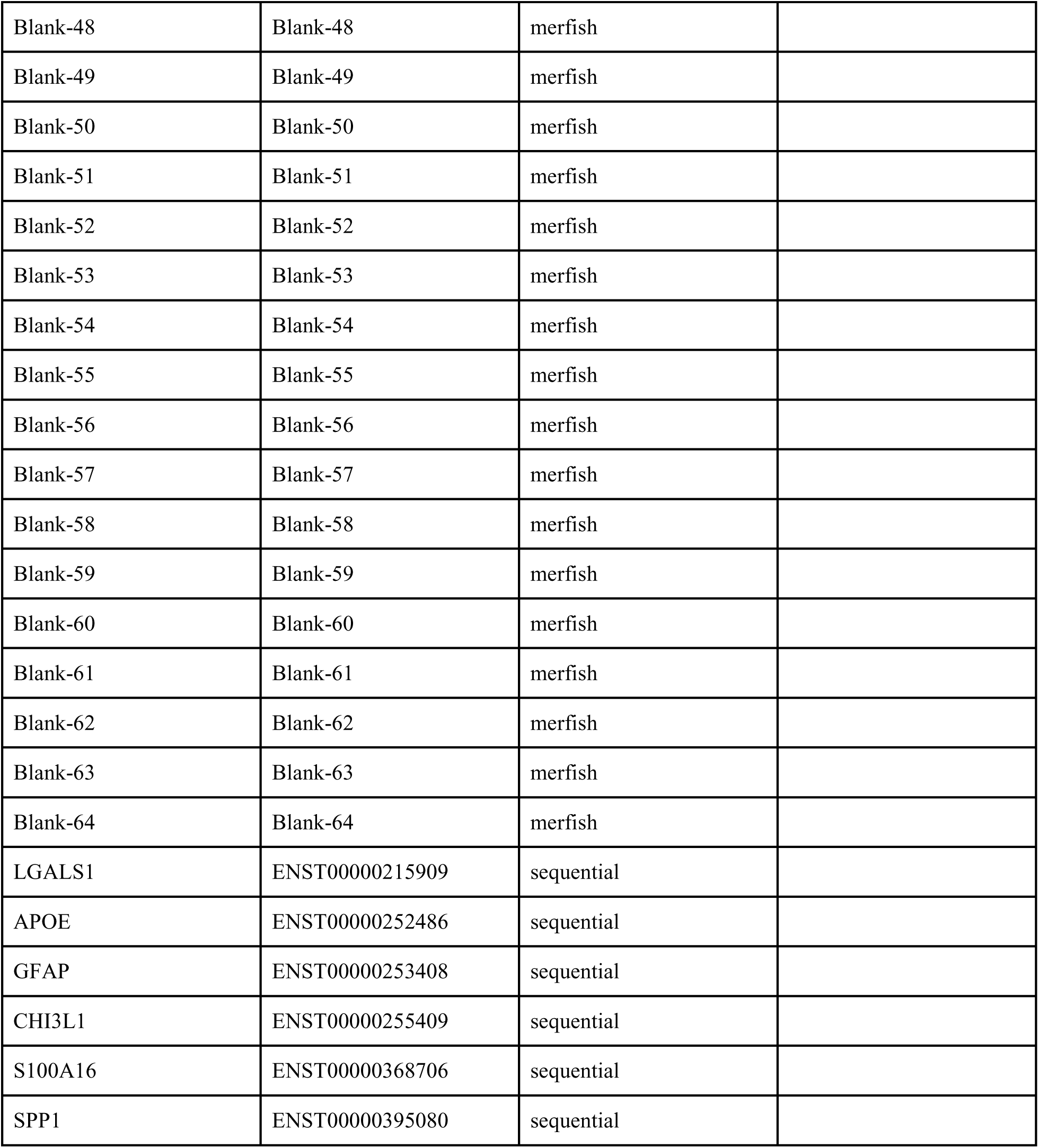
Custom MERFISH gene panel.

